# A small RNA is linking CRISPR-Cas and zinc transport

**DOI:** 10.1101/2021.02.19.431937

**Authors:** Pascal Märkle, Lisa-Katharina Maier, Sandra Maaß, Claudia Hirschfeld, Jürgen Bartel, Dörte Becher, Björn Voß, Anita Marchfelder

## Abstract

The function and mode of action of small regulatory RNAs is currently still understudied in archaea. In the halophilic archaeon *H. volcanii* a plethora of sRNAs have been identified, however, in-depth functional analysis is missing for most of them. We selected a small RNA (s479) from *H. volcanii* for detailed characterization. The sRNA gene is encoded between a CRISPR RNA locus and the Cas protein gene cluster, the s479 deletion strain is viable and was characterized in detail. Transcriptome studies of wild type *Haloferax* cells and the deletion mutant revealed up-regulation of six genes in the deletion strain, showing that the sRNA has a clearly defined function. Three of the six up-regulated genes encode potential zinc transporter proteins (ZnuA1, ZnuB1, ZnuC1) suggesting involvement of s479 in regulation of zinc transport. Upregulation of these genes in the deletion strain was confirmed by northern blot and proteome analyses. Furthermore, electrophoretic mobility shift assays demonstrate a direct interaction of s479 with the target *znu*C1 mRNA. Proteome comparison of wild type and deletion strains further expanded the regulon of s479 deeply rooting this sRNA within the metabolism of *H. volcanii* especially the regulation of transporter abundance. Interestingly, s479 is not only encoded next to CRISPR-*cas* genes but the mature s479 contains a crRNA-like 5’ handle and experiments with Cas protein deletion strains indicate maturation by Cas6 and interaction with Cas proteins. Together this might suggest that the CRISPR-Cas system is involved in s479 function.

## Introduction

Small RNAs have been well established as key-regulators of gene expression in both pro- and eukaryotic species (Storz et al., 2011; Wagner and Romby, 2015; Buddeweg et al., 2018a). But still, understanding of small RNAs (sRNAs) in the archaeal domain lags behind (Gomes-Filho et al., 2018). RNomics as well as more recent high-throughput approaches have been applied to several archaeal species to uncover the wealth of small transcripts found in this domain of life (reviewed in: (Marchfelder et al., 2012; Kliemt and Soppa, 2017; Buddeweg et al., 2018a; Gelsinger and DiRuggiero, 2018a)). With numbers in the hundred (*Archaeoglobus fulgidus*, *Methanosarcina mazei*, *Sulfolobus solfataricus*, *Thermococcus kodakarensis*, *Pyrococcus abyssi, Haloferax mediterranei*) or even thousand (*Haloferax volcanii*, *Methanolobus psychrophilus*), sRNAs are well established as widespread and abundant players within the transcriptome of various archaeal species (reviewed in: (Gelsinger and DiRuggiero, 2018a; Gomes-Filho et al., 2018; Payá et al., 2018)).

sRNAs from the haloarchaeal model organism *H. volcanii* have been studied since more than ten years (Straub et al., 2009; Babski et al., 2011; Fischer et al., 2011; Heyer et al., 2012). Several recent RNA-seq and differential RNA-seq (dRNA-seq) studies explored the small RNome of *H. volcanii* in greater depth and uncovered an unexpected wealth of potential small RNAs expanding the number of sRNA candidates from just about 200 identified in 2009 to now well over 1,000 candidate sRNAs (Heyer et al., 2012: 190 sRNAs; Babski et al., 2016: 1,701 candidates; Laas et al., 2019: 1,635 candidates; Gelsinger and Diruggiero, 2018b: 1,533 candidates) (Heyer et al., 2012; Babski et al., 2016; Gelsinger and DiRuggiero, 2018b; Laass et al., 2019). Depending on their genomic localization small regulatory RNAs are categorized as trans-encoded intergenic sRNAs (sRNAs) or cis-encoded anti-sense RNAs (asRNAs) overlapping with annotated reading frames of the opposite strand. In contrast to asRNAs, for whom targets are readily deducible as they are by default able to extensively base pair with the transcript originating from the opposite strand, trans-acting sRNAs pose quite a challenge as to the identification of targeted mRNAs. More so, as they bind targets via imperfect base-pairing sRNAs may regulate multiple targets as demonstrated in bacteria (Papenfort and Vogel, 2009; Wagner and Romby, 2015) and by the few archaeal examples (reviewed in: (Buddeweg et al., 2018a; Gelsinger and DiRuggiero, 2018a; Gomes-Filho et al., 2018)).

It is well established that intergenic sRNAs of *H. volcanii* are differentially expressed in response to growth phase or environmental stimuli including temperature, salinity, as well as oxidative stress (Straub et al., 2009; Babski et al., 2011; Fischer et al., 2011; Gelsinger and DiRuggiero, 2018a). Their profound biological role is unquestioned as growth phenotypes have been demonstrated for sRNA deletion mutants in response to temperature, salt, alternative carbon-sources, phosphate availability and oxidative stress (Straub et al., 2009; Fischer et al., 2011; Heyer et al., 2012; Jaschinski et al., 2014; Kliemt et al., 2019). Besides growth, swarming behaviour or cell shape have been affected by sRNA deletions as well (Jaschinski et al., 2014). sRNA-mediated regulation may involve a range of sRNAs as differential transcriptome analyses in the context of oxidative stress implies (Gelsinger and DiRuggiero, 2018b). A recent metatranscriptome study highlights the importance of sRNA-based regulation for the archaeal metabolism once more demonstrating differential expression of sRNAs in a halo-extremophile community inside salt rocks of the Atacama Desert in response to environmental changes on a population-wide-scale (Gelsinger et al., 2020). This underpins that sRNAs do not take the side-line in archaeal gene regulation but are central players for stress adaptation in large scale perspective. Despite this immense body of evidence as to the involvement of sRNA mediated regulation in processes of metabolic and stress adaptation, data on sRNA-target pairs is still scarce.

The intergenic sRNA s479 of *H. volcanii* described herein gained our interest, as sRNA resulting in a growth phenotype upon deletion but also as RNA encoded between genes for the CRISPR-Cas system. Differential transcriptome and proteome analysis revealed changes in several zinc-related mRNAs and proteins, respectively. Analyses of the deletion strain in conjunction with electrophoretic mobility shift data and impaired growth under elevated zinc concentrations confirm a role of s479 within the zinc regulon. Differential proteome analysis reveals a role for s479 in the adjustment of a network of ABC transporters. Furthermore s479 seems to depend on Cas proteins for maturation and stability.

## Results

### Characterisation of s479

The s479 RNA was identified in an early RNomics study, sequencing cDNA clones after RNA size-selection (Straub et al., 2009). It is an intergenic sRNA located on the genomic plasmid pHV4 downstream of the CRISPR locus P1 and upstream of the type I-B *cas* gene cassette (Figure 1). A genome-wide high-throughput study analysing transcriptional start sites of *H. volcanii* (Babski et al., 2016) revealed an enrichment of transcript starts at position 207,660.

**Figure 1.**
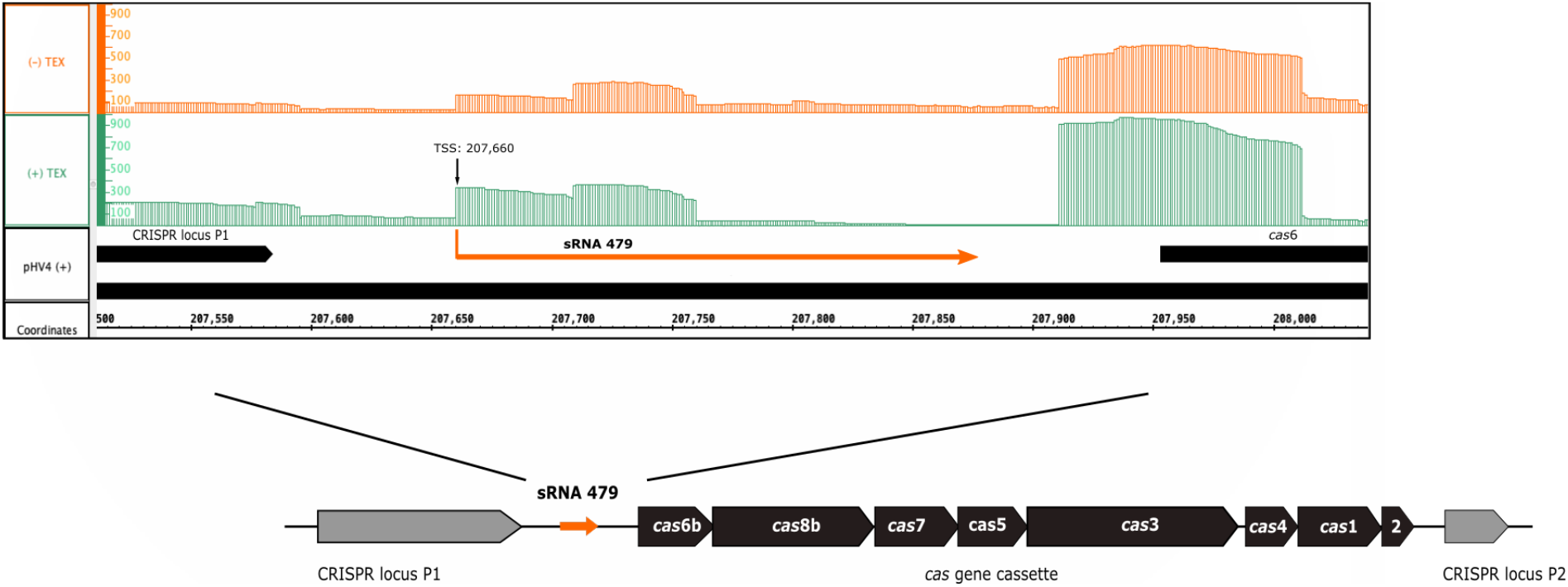
Genomic location of s479. The sRNA 479 is encoded on the genomic plasmid pHV4 and as read abundance shows (Jaschinski et al., 2014; Babski et al., 2016), transcription initiates at position 207,660. Transcription start site data (TSS) together with the previously obtained RNomics data (Straub et al., 2009), show that the s479 gene is 213 bp long. Location of s479 is noteworthy, as it is encoded in the intergenic region between a CRISPR locus and the *cas* gene cassette. Reads obtained from RNA treated with terminal exonuclease (+TEX) are shown in green, reads from untreated RNA (-TEX) are shown in orange. Comparison of reads from both fractions allowed us to determine the TSS (indicated by an arrow). Genome coordinates and annotated genes of the main chromosome plus strand are shown in black at the bottom.

Expression of s479 was confirmed by northern blot analysis revealing long RNAs of about 220 and 160 nucleotides and a very strong cluster of signals of approximately 51 nucleotides (Figure 2), showing, that the primary s479 transcript is processed yielding an RNA of about 51 nucleotides. The northern data confirm the dRNA-seq results presented in Figure 1, which show that RNAs starting at the mature s479 5’ end (position 49 in Figure 3) with about 50 nucleotides length have the most reads.

**Figure 2.**
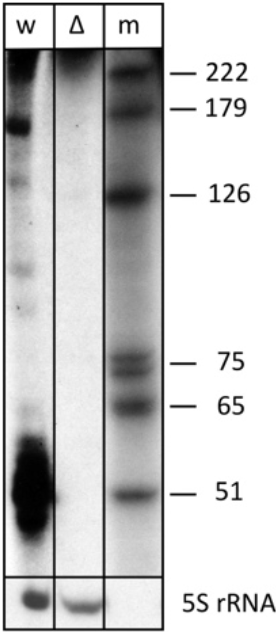
The s479 precursor transcript is processed into a shorter RNA. Total RNA from wild type (lane w) and Δ*s479* (lane Δ) cells was separated on a polyacrylamide gel. After transfer, the membrane was hybridized with a probe against s479 (upper panel, the lower panel was hybridised with a probe against the 5S rRNA). Lane w: RNA from *H. volcanii* wild type strain, lane Δ: RNA from a Δ*s479* strain, lane m: size marker, sizes are given at the right in nucleotides.

**Figure 3.**
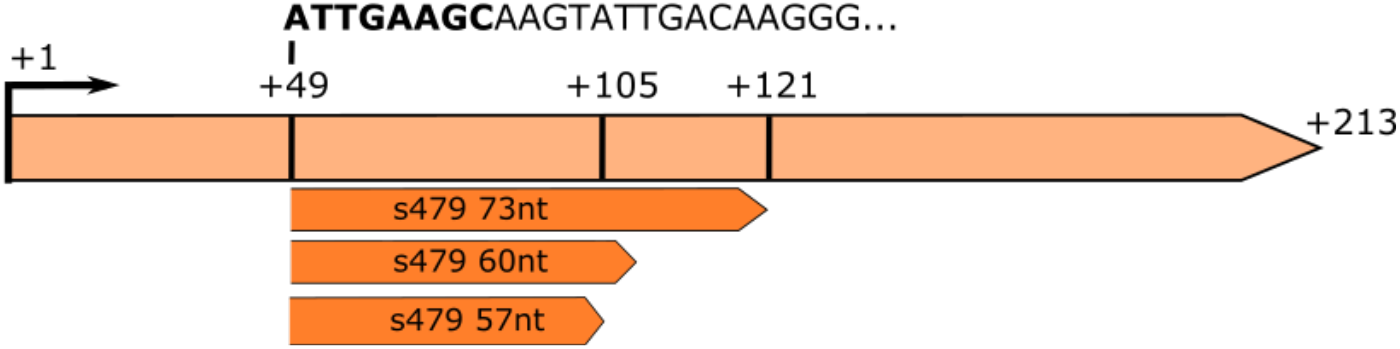
RNA-seq analysis of small RNAs reveals sequences matching the s479 locus. The s479 precursor RNA is 213 nucleotides long, shown in light orange. RNomics analysis of size-selected RNA samples (Maier et al., 2013) reveal shorter versions of s479; 57 nt, 60 nt and 73 nt in length, shown below in dark orange. These data confirm the results of the northern blot (Figure 2). The sequence of the shorter sRNAs starts at nucleotide 49 of the precursor. The first 8 nucleotides are identical to crRNA 5’ handle sequence (shown in bold: ATTGAAGC) (Supplementary Figure 2).

This is also supported by a serendipitous finding: an RNA-seq study of small RNAs to identify CRISPR RNAs revealed reads mapping to the s479 locus; a 57 nt, 60 nt as well as a 73 nt form (Figure 3; Supplementary Figure 1) (Maier et al., 2013). These correspond in size to the cluster of bands visible in the northern blot analysis (Figure 2 and Figure 10).

To elucidate the importance of s479 in the context of *H. volcanii* metabolism in more detail, we first compared growth of the s479 deletion strain (Δ*s479*) obtained in a previous study (Jaschinski et al., 2014) with the H66 wild type strain (Figure 4). Growth curves show a diauxic growth, doubling times of wild type and deletion strains in phase 1 (0.5 h - 10.5 h) are not very different with doubling times of 4.7 h for the wild type strain and 4.5 h for the deletion strain (see also Supplementary Figure 5C for doubling time details). However in phase 2 (14.5 h - 24.5 h) the doubling time of the deletion strain is longer with 24 hours compared to 19 hours of the wild type strain. In addition, the deletion strain reaches a lower OD in stationary phase, showing that the sRNA has an important function in the cell.

**Figure 4.**
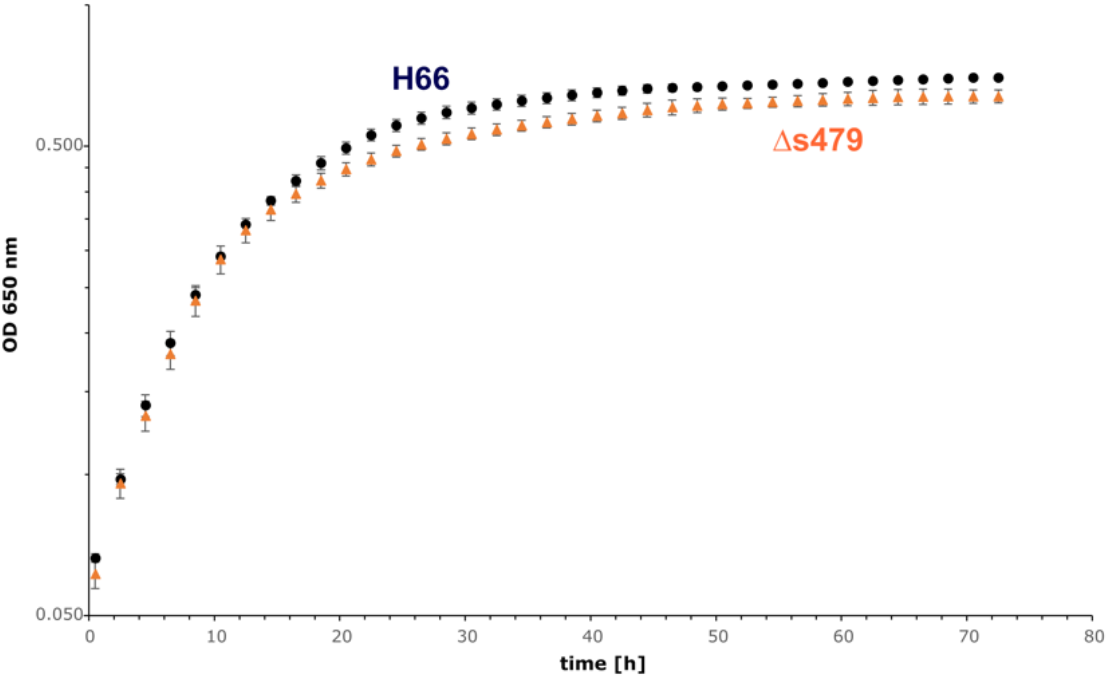
Growth experiment comparing wild type strain and Δ*s479* strain. Both strains were cultivated in triplicate with Hv-Ca medium (supplemented with uracil) in microtiter plates and OD_650nm_ was monitored using a heated plate-reader instrument.

### Influence of s479 on the *H. volcanii* transcriptome

As deletion of s479 led to a modest growth phenotype, we wanted to identify genome wide changes in the *H. volcanii* gene expression profile resulting from loss of sRNA expression. We performed RNA-seq analysis of the sRNA deletion strain Δ*s479* and the wild type strain H66 grown to exponential phase. To pin-point transcripts affected by absence of s479 we applied a stringent differential transcriptome analysis.

Bioinformatical analysis identified one transcript as down-regulated, the s479 RNA, and five transcripts as up-regulated with a log_2_fold-change≥2. As Table 1 shows, two of the up-regulated genes are derived from a single genomic region on the main chromosome comprising amongst others the operon *znu*A1C1B1 encoding a putative zinc-ABC-transporter (Figure 5).

**Figure 5.**
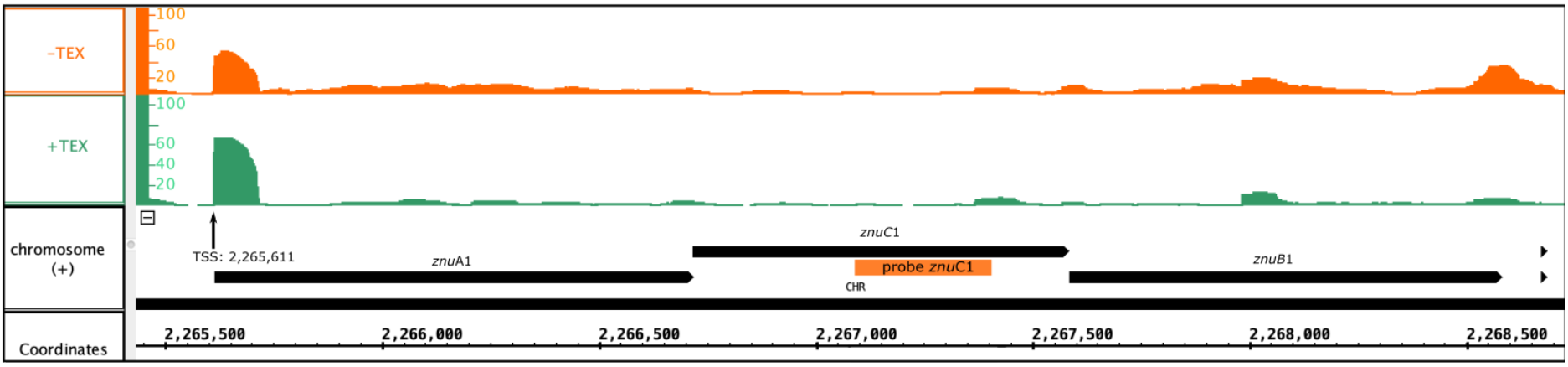
Genomic localization of the *znu* operon. Depicted is the localization of the *znu* operon alongside the read coverage at TSS (Babski et al., 2016) confirming the joint transcription of all three *znu* genes as an approximately 3,000 nt transcript. The location of the PCR probe used for *znu*C1 detection in northern blot hybridization is given as orange bar. Reads obtained from RNA treated with terminal exonuclease (+TEX) are shown in green, reads from untreated RNA (-TEX) are shown in orange. Comparison of reads from both fractions allowed us to determine the TSS (indicated by an arrow). Genome coordinates and annotated genes of the main chromosome plus strand are shown in black at the bottom.

**Table 1.**
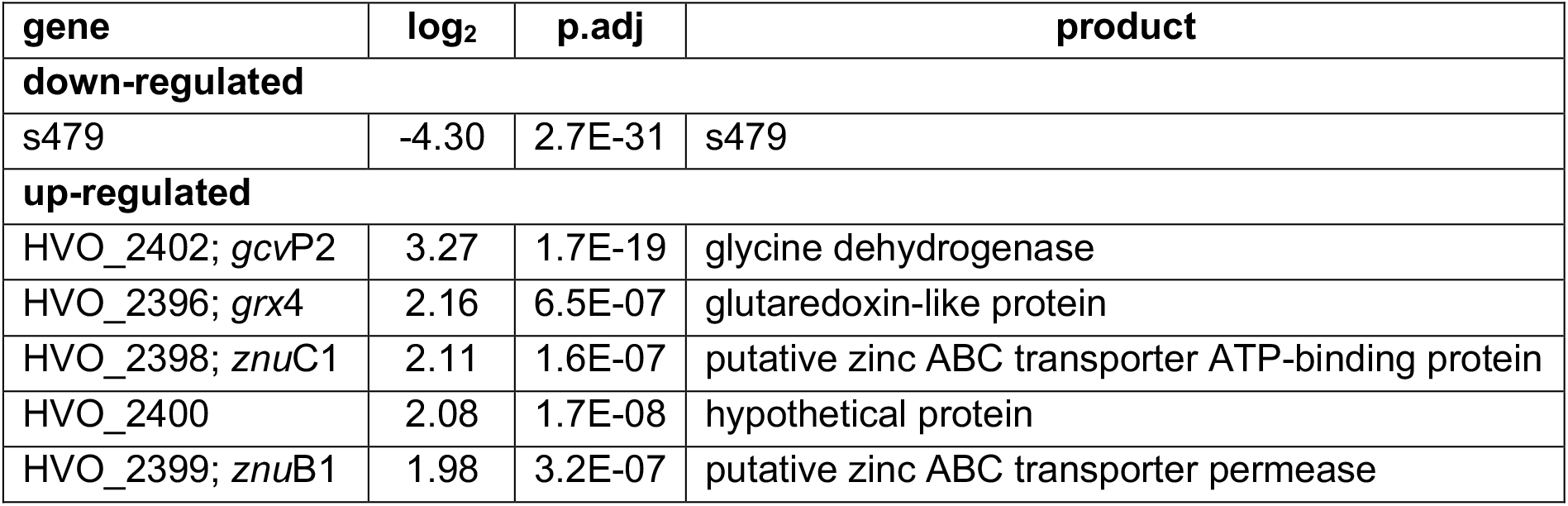
Six genes are differentially expressed in the deletion strain Δ*s479*. The log_2_ fold change (column log_2_) deletion vs. wild type is given alongside the HVO-gene number (column gene), p-value (column p.adj) and gene product name (column product).

| gene | log<sub>2</sub> | p.adj | product |
| --- | --- | --- | --- |
| **down-regulated** |  |  |  |
| s479 | -4.30 | 2.7E-31 | s479 |
| **up-regulated** |  |  |  |
| HVO_2402; *gcvP2* | 3.27 | 1.7E-19 | glycine dehydrogenase |
| HVO_2396; *grx4* | 2.16 | 6.5E-07 | glutaredoxin-like protein |
| HVO_2398; *znuC1* | 2.11 | 1.6E-07 | putative zinc ABC transporter ATP-binding protein |
| HVO_2400 | 2.08 | 1.7E-08 | hypothetical protein |
| HVO_2399; *znuB1* | 1.98 | 3.2E-07 | putative zinc ABC transporter permease |

Abundance of the two transcripts, *znu*C1 and *znu*B1 increases more than four-fold in response to s479 deletion. *znu*A1, encoding the periplasmic substrate-binding protein of said putative ABC transporter, is also present within the set of differentially expressed genes but fell just below the threshold of log_2_FC≥2 with a score of 1.8 (Supplementary Table 1).

As abundance of all three genes of the *znu* operon is altered upon s479 deletion, we further concentrated our analysis on the *znu* operon comprising *znu*A1 (HVO_2397), *znu*B1 (HVO_2399) and *znu*C1 (HVO_2398). TSS analyses (Babski et al., 2016) show that expression is governed by a single TSS four nucleotides upstream of the *znu*A1 start codon, resulting in a multicistronic mRNA of approximately 3,000 nt (Figure 5). Such a short 5’ UTR is typical for *H. volcanii* which has a high percentage of leaderless mRNAs and 5’ UTRs shorter than 6 nucleotides (Babski et al., 2016).

The transcript differences seen for the *znu* operon were validated using northern blot analysis probing for part of the *znu*C1 coding sequence (Figure 6). The *znu* transcript is more abundant in the strain without s479. In addition, results of the northern blot analysis confirm transcription of the *znu* operon as a single polycistronic transcript of about 3,000 nucleotides (Figure 6).

**Figure 6.**
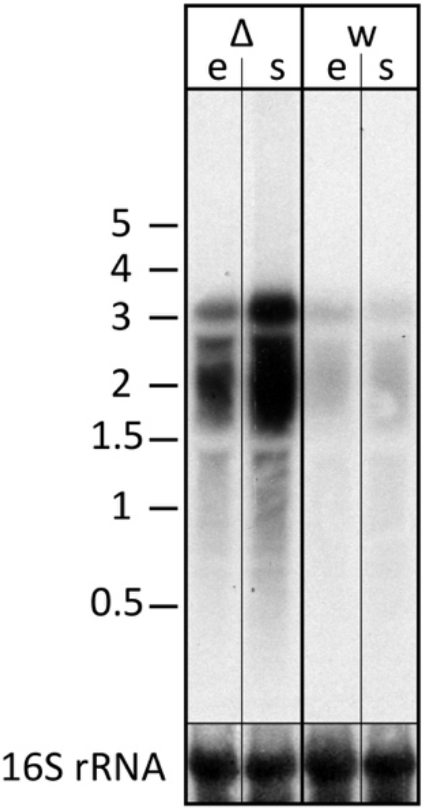
Transcript levels of the *znu* operon in wild type and s479 deletion strain. We compared the transcript levels of *znu* in total RNA of wild type H66 (lanes w) and Δ*s479* (lanes Δ) cells at exponential and stationary phase (lanes e and s, respectively) using northern blot analysis. The probe used for hybridisation in the upper panel is located in the central part of the *znuC1* coding sequence (Figure 5). Signals at about 3,000 nucleotides correspond in length to the complete *znu* operon mRNA, signals at approximately 2,000 correspond in length to a bicistronic mRNA encompassing either *znu*A1 and *znu*C1 or *znu*C1 and *znu*B1. In the lower panel the blot was hybridised with a probe against the 16S rRNA. A size marker is given in kb at the left.

### Influence of s479 on protein abundance

Taking the analysis of the 479 target-sphere a step further, we compared proteomes of the wild type and the s479 deletion strain. Since we saw differential expression of transporter genes in the transcriptome study, we used separate protocols for maximum recovery of soluble as well as membrane-associated proteins for protein extraction and samples were then analysed separately by mass spectrometry (MS). In total, 22 proteins were exclusively present in the s479 deletion strain, whereas 18 proteins were not detected in Δ*s479* (Table 2 and Supplementary Table 2).

**Table 2.** Transporter components exclusively present or accumulated in the deletion strain. Differential proteome analysis comparing the s479 deletion strain and wild type strain was used determining the log_2_ fold change deletion vs. wild type (column on/log_2_). If detected only in the deletion strain the protein is listed as "on". Additionally, HVO-gene numbers (column gene) encoding the proteins are given alongside the gene product (column product) and Kegg/COG assignments (column Kegg pathway/ COG assignment) informing on the metabolic pathway(s). Proteins encoded by the *znu* operon are given in bold.

| gene | product | on/<br>log<sub>2</sub> | Kegg pathway/ COG<br>assignment |
| --- | --- | --- | --- |
| HVO_B0198 | ABC-type transport system<br>periplasmic substrate-binding protein<br>(probable substrate iron-III) | on | hvo02010 - ABC<br>transporters |
| HVO_B0047 | ABC-type transport system<br>periplasmic substrate-binding protein<br>(probable substrate iron-III) | on | hvo02010 - ABC<br>transporters |
| HVO_2375 | putative phosphate ABC transporter<br>periplasmic substrate-binding protein | on | hvo02010 - ABC<br>transporters |
| **HVO_2397** | **putative zinc ABC transporter**<br>**periplasmic substrate-binding**<br>**protein** | **on** | hvo02010 - ABC<br>transporters |
| HVO_1705 | putative iron-III ABC transporter<br>periplasmic substrate-binding protein | on | hvo02010 - ABC<br>transporters |
| **HVO_2398** | **putative zinc ABC transporter**<br>**ATP-binding protein** | **2.5** | hvo02010 - ABC<br>transporters |
| HVO_2324 | pantothenate permease | 2.1 | COG0591 Code<br>ER Na<sup>+</sup>/proline symporter |

For 12 proteins, a significant differential abundance with log_2_ fold change≥2 was measured. Eight of them accumulated and four were depleted in the deletion strain. A summary of the proteomic changes detected with a log_2_ fold change≥0.8 is given in Supplementary Table 2. Comparison with the transcriptome data shows that the increase in transcript level seen for the *znu* operon is paralleled by an increase in protein level (*znu*C1, HVO_2398) or the exclusive detection of the gene product in the deletion strain (*znu*A1, HVO_2397). Analysis of the Kegg pathway assignment of the uniquely or differentially present proteins reveals an enrichment of transporter proteins exclusively present in the deletion strain (5 of 22) (Table 2; Supplementary Table 2). But the majority of proteins present in the deletion strain only are hypothetical proteins (9 of 22). Since deletion of the s479 gene alters expression of zinc transporters we compared growth of wild type and s479 strains in low zinc and high zinc concentrations. Under low zinc concentrations wild type and Δ*s479* show the same growth behaviour (data not shown). However upon addition of high zinc concentrations Δ*s479* shows defects in growth (Supplementary Figure 5).

### *In silico* analysis of the sRNA-target interaction

As little is known about sRNA-target interactions in the archaeal domain and the few examples described so far reveal a diverse set of interaction modes, the potential interaction of s479 with the *znu*C1 coding region was further analysed using bioinformatics. The sequence of the s479 (Figure 7B.) was utilized to predict interactions with the *znu* operon transcript by the IntaRNA suite (Mann et al., 2017). We were able to predict two potential interaction sites, both within the *znu*C1 open reading frame, using standard settings (Figure 7A.). Site 1 is located 120 nt downstream of the first nucleotide of the *znu*C1 coding sequence and site 2 is located 408 nt downstream of it with predicted energy gain of E= −7.82 for site 1 and E=−10.27 for site 2, respectively. The predicted interacting sequence of s479 is situated at the 3’ end.

**Figure 7.**
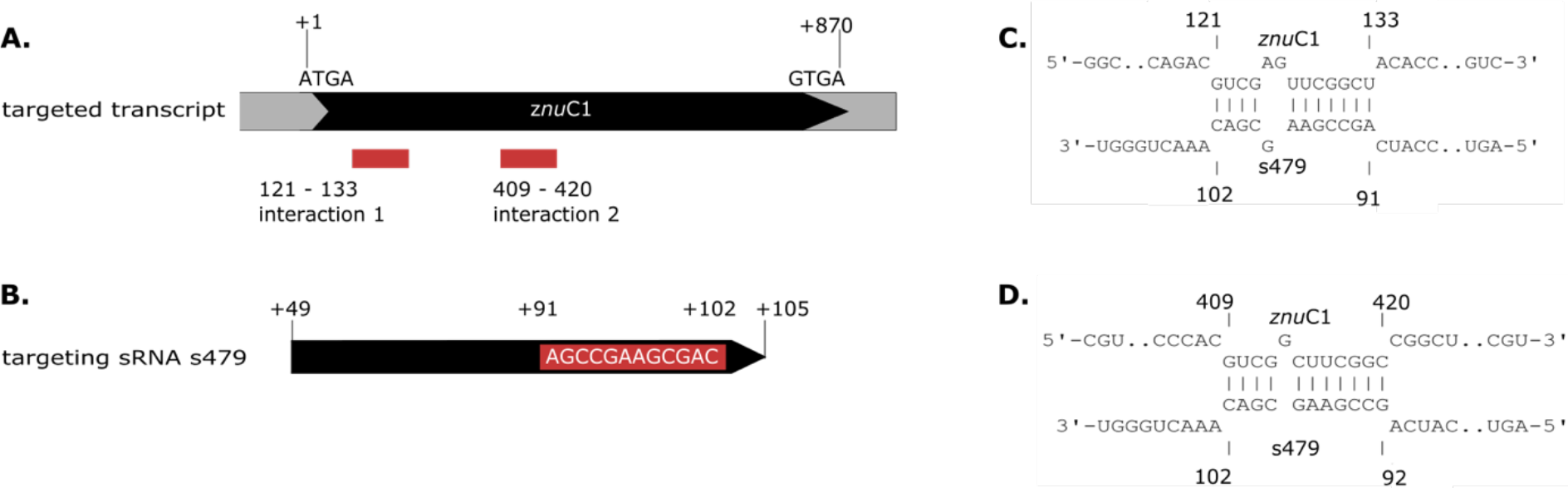
Interacting sequences of the *znu*C1 transcript and s479. **A.** The two potential interaction sites for the *znu*C1 transcript are indicated by red boxes. The coding region is shown in black and numbering starts with the A of the start codon ATG. **B.** The mature s479 is shown, numbering is according to the full primary transcript as shown in Figure 3, where the mature s479 starts at +49. The part of s479 interacting with the *znu*C1 mRNA is shown in the red box, situated at the 3’-end. The predicted target-sRNA binding configuration is depicted in **C.** for interaction 1 and in **D.** for interaction 2.

As proteome analysis revealed additional potential targets of s479, the IntaRNA analysis was extended to the transcripts of these proteins, as well. Since translational regulation commonly involves sequences at or in close proximity to the first codon, we incorporated the sequences from transcriptional start site to 50 nucleotides downstream^1^. Analysis was confined to proteins exclusively present or absent in the deletion strain, as these were the most affected. Using standard parameters, interaction with s479 was predicted for eight transcripts. We then modified the search parameters to also include seed sequences of only five nucleotides and relaxed specificity and this resulted in 21 predicted interactions. All of those interactions involve a similar region within s479 around position 44 to 54 at the 3’-end of the mature s479 (Figure 8). The weblogo (Crooks, 2004) created for the s479 nucleotides involved in contacting all these transcripts highlights an nine nucleotide consensus motif (GCCGAAGCG) corresponding to the interaction surface predicted for the s479-*znu*C1 contact.

**Figure 8.**
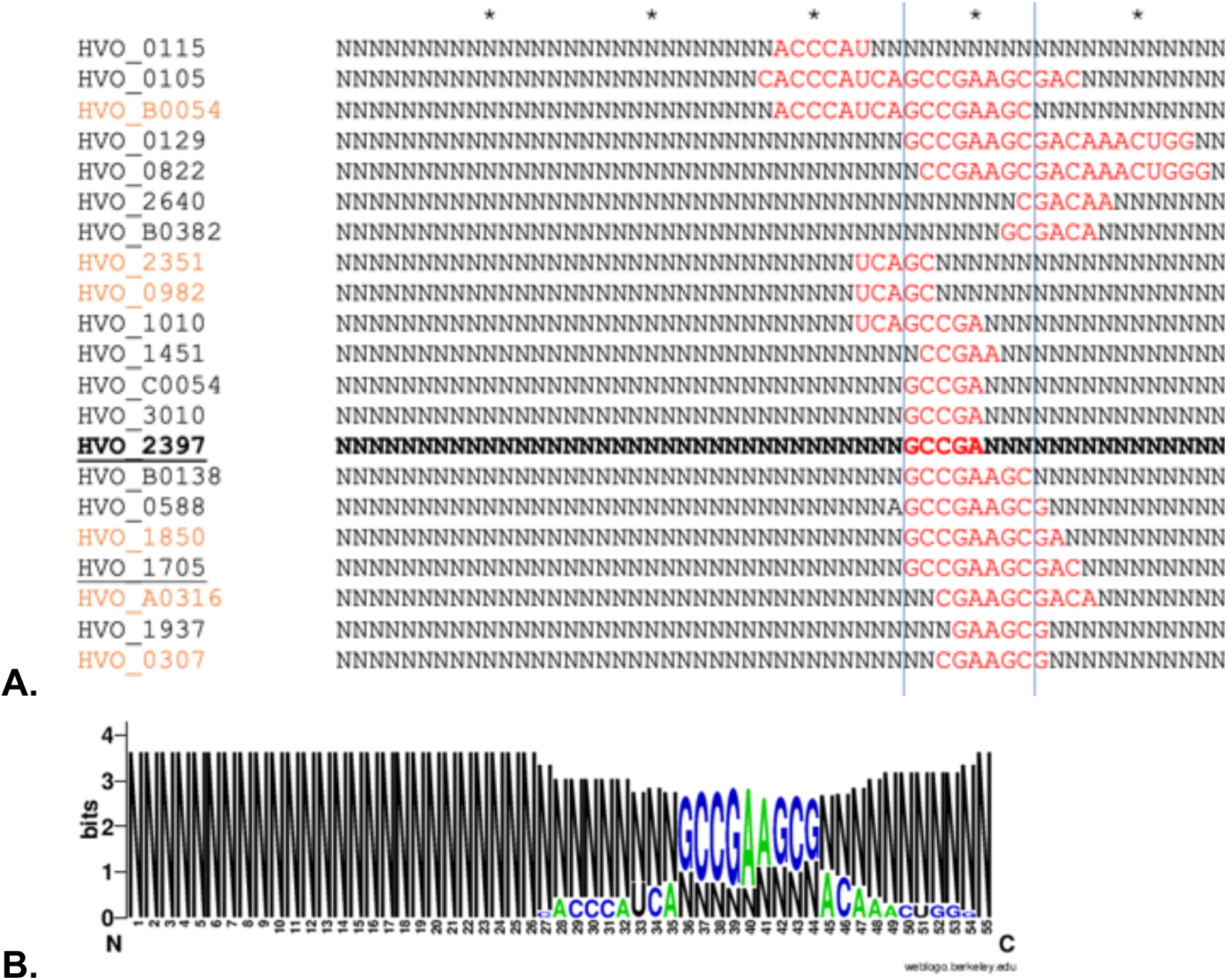
Predicted s479-target interaction sites for proteomic targets. **A.** Results of the IntaRNA prediction of interacting sequences of s479 (Mann et al., 2017). The respective targeted mRNA is given at the left and for the s479 sequence, all nucleotides not taking part in the predicted interaction (red) are depicted as N. The asterisk shown in the upper line marks ten nucleotides. The proteomic targets included are those exclusively present or absent in Δ*s479* in comparison to H66. The *znu*A1 interaction site is given in bold, transporter targets are underlined, and targets predicted with seed-length >5 are given in orange. **B.** The resulting conserved interaction sequence of s479 with its targets is shown as weblogo at the right (Crooks, 2004).

### Verification of the s479-*znu*C1 target interaction

Assessment of the transcriptome by dRNA-seq as well as northern blot analysis confirms that abundance of the *znu*C1 transcript is altered upon deletion of the s479 (Table 1, Figure 6). Moreover, target site prediction suggests two interaction surfaces on the *znu*C1 mRNA (Figure 7). To validate whether the *znu*C1 transcript is a direct interaction partner of s479 and whether the predicted interaction site is correct, we performed an electrophoretic mobility shift assay (Figure 9).

**Figure 9.**
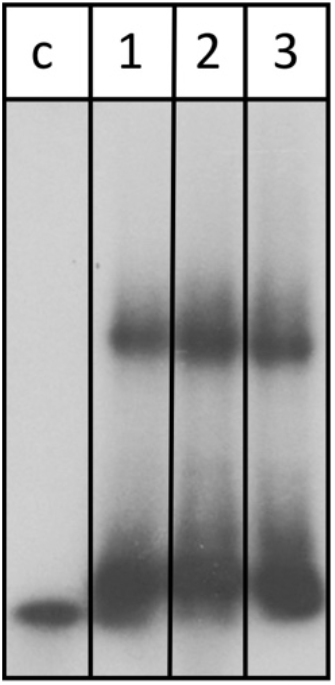
s479 binds to the *znu*C1 mRNA *in vitro*. Labelled s479 was incubated with increasing concentrations of a *znu*C1 mRNA fragment, containing the predicted interaction site 2 (Figure 7D). Reactions were loaded onto a non-denaturing polyacrylamide gel. Lane c: control reaction without addition of *znu*C1, lane1: addition of 50 pmol *znu*C1, lane 2: addition of 100 pmol *znu*C1, lane 3: addition of 200 pmol *znu*C1.

Labelled s479 RNA was incubated with increasing amounts of a unlabelled *znu*C1 mRNA fragment, comprising the predicted interaction site 2 (Figure 7D.) and flanking nucleotides (28 nt upstream, 40 nt downstream), gel shift analysis shows that s479 binds to the *znu*C1 RNA (Figure 9). Competition experiments with unlabelled s479 were also performed (Supplementary Figure 3A.) revealing that the unlabelled s479 competes effectively with the radioactively labelled one for binding to the *znu*C1 RNA. However, sRNA s479 does not bind to a *znu*C1 mutant RNA, which has the s479 interaction site (Figure 7D.) deleted (Supplementary Figure 3B.).

### s479 is bound by Cascade

The s479 precursor RNA contains almost a complete CRISPR repeat sequence (Supplementary Figure 2). Processing at the Cas6 cleavage site of this repeat sequence would yield the mature s479 RNAs containing a 5’ sequence identical to the characteristic eight nucleotide long 5’ handle of the mature *H. volcanii* crRNAs of locus P1 (5’ ATTGAAGC 3’) (Maier et al., 2013). This observation led us to investigate, whether the Cas6 protein is involved in s479 biogenesis. Northern blot analysis shows that the short RNAs with about 50 nucleotides length are lost upon *cas6* deletion (Figure 10), only an intermediate RNA of about 100 nucleotides is detected, suggesting that Cas6 indeed generates the mature sRNA.

**Figure 10.**
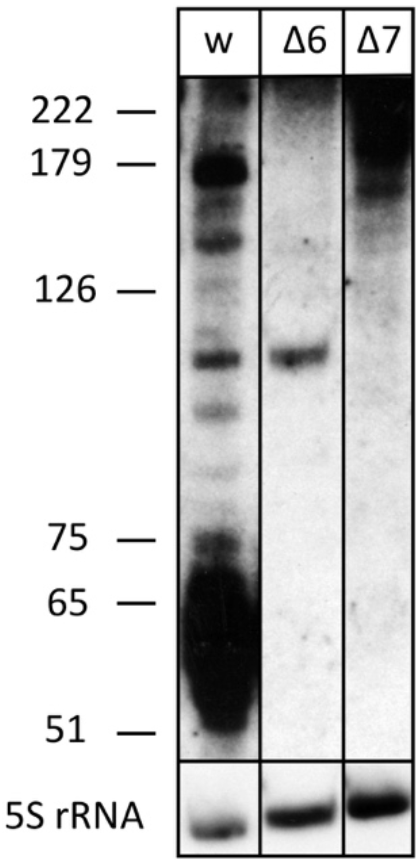
The processed s479 species are lost upon deletion of genes for the Cas6 or Cas7 protein. Total RNA was separated on a polyacrylamide gel. After transfer, a probe against s479 was used for hybridization (upper panel). A size marker is given at the left. Lane wt: RNA from *H. volcanii* H119 wild type strain, lane Δ6: RNA from *H. volcanii cas6* deletion strain Δ*cas6*, lane Δ7: RNA from *H.volcanii cas7* deletion strain Δ*cas7*. In the lower panel hybridisation with a probe against 5S rRNA is shown.

We also analysed whether s479 is present in a *cas7* deletion strain. Earlier investigation showed, that Cas7 is the central subunit of the *Haloferax* Cas protein complex Cascade which binds the crRNAs. In a *cas7* deletion strain, the Cascade complex cannot form anymore, thus crRNAs are not bound to Cascade and therefore are not stable anymore (Brendel et al., 2014). We used RNA from such a *cas7* deletion strain to monitor the s479 interplay with the Cascade complex. Indeed, in a Δ*cas7* strain s479 is not detectable anymore, suggesting that s479 is bound and protected by Cascade as observed for crRNAs. Such an interaction with Cascade would be the first example for a non-crRNA bound by a type I-B Cascade used for internal gene expression regulation.

To determine whether the *znu* operon is up-regulated in Δ*cas6* and Δ*cas7* strains we probed a northern membrane containing RNA from these deletion strains with the *znu*C1 probe (Figure 5, Supplementary Figure 4). Higher concentrations of *znu* mRNA are detected in the Δ*cas7* strain (Supplementary Figure 4), as expected when s479 is missing. Interestingly, in the Δ*cas6* strain the *znu* RNA is not up-regulated although the mature s479 is not present (Figure 10). Thus the longer s479 intermediate with about 100 nucleotides length present in the Δ*cas6* strain seems to be also active in regulating the *znu* RNA.

## Discussion

Despite decades of research archaeal sRNA networks are still enigmatic. It is well established that archaeal sRNAs play crucial roles in gene regulatory networks related to metabolism and therefore are essential players in stress responses, but pin-pointing their interaction partners remained challenging (Buddeweg et al., 2018a). An inventory of small transcripts has been made for more than seven archaeal species, but only for four of them sRNA-target pairs have been identified (Buddeweg et al., 2018b; Gelsinger and DiRuggiero, 2018b; Orell et al., 2018). We report here data for the second sRNA-target RNA pair of *H. volcanii*.

### Deletion of the s479 gene has an impact on growth and the transcriptome

*H. volcanii* encodes two operons for putative zinc ABC transporters (*znu*A1-C1 and *znu*A2-C2 genes), without s479 a higher abundance of transcripts for one of the two operons is observed (*znu*A1-C1), which is confirmed by northern blot analysis. Moreover, proteome analysis comparing s479 deletion and wild type also identifies *znu*1 gene products as differentially regulated. In addition, our data support a direct interaction of s479 with the transcript of the zinc transporter gene *znu*C1. IntaRNA prediction reveals that the 3’ end of the s479 interacts with the coding region of *znu*C1. The energy gain predicted for the interaction implies a stable sRNA-target pairing (Kliemt et al., 2019). The few examples of archaeal sRNA-target pairs described to date suggest a non-universal mode of action and a wide variability in the site of target contact (Buddeweg et al., 2018b; Gomes-Filho et al., 2018). The binding site for s479 within the coding region of the targeted mRNA is a feature shared with the only other sRNA-target pair described for *H. volcanii* (sRNA s132) (Kliemt et al., 2019) and with examples from other species (*M. mazei* s154, *S. acidocaldarius* RrrR(+)) (Prasse et al., 2017; Orell et al., 2018). EMSA analysis revealed s479 binding to the *znu*C1 RNA (Figure 9), confirming a direct interaction of both RNAs. This is the first validation of a sRNA-target pair by gel shift for *H. volcanii*. We hereby also confirm that s479 is exerting direct control of the *znu*C1 transcript. This results in a negative effect on *znu*C1 abundance in the cellular context which can be released by s479 deletion as seen in northern blot analysis. Destabilizing the target by potentially binding within the coding sequence is also suggested for *S. acidocaldarius* RrrR(+) (Orell et al., 2018) but archaeal RNases have not been studied in great depth yet and therefore, no candidate RNase is evident for direct degradation of dsRNA (no RNase III activity has been described in archaea) or for ssRNA cleavage upon dsRNA formation (Clouet-d’Orval et al., 2018).

The s479 deletion strain has a slight growth disadvantage compared to the wild type during late exponential growth. Growth rate of the deletion strain is retarded by 22 % compared to wild type during late exponential phase resulting in a lower end point of growth as well, showing that the sRNA is required for normal growth. The fact that s479 only regulates one of the two *znu* operons might be the reason that deletion of the s479 gene has only a slight impact on growth. The second *znu* operon could be regulated by another sRNA or other factors and concerted regulation of both *znu* operons might require an as yet unknown master regulator.

### Deletion of the s479 gene has a severe impact on the proteome

The differential proteome analysis revealed a much larger regulon for s479 on protein level than seen on the transcript level suggesting that the primary effect of s479 is at the translational level. s479 is severely affecting the presence of more than 50 proteins. In contrast to the transcriptional level, where s479 acts as negative regulator, proteome analysis revealed proteins less abundant in the deletion strain, too. Such a duality in the direction of regulation achieved by a single sRNA has also been described for the other *H. volcanii* sRNA s132 (Kliemt et al., 2019). And like s132, s479 is implicated in both, accumulation and depletion of certain proteins. Whether this effect is direct or indirect via intermediate gene products regulated by s479 must be analysed in future experiments. But it already demonstrates that sRNA-based regulation in *H. volcanii* is a complex and multimodal process in case of both sRNAs as they are addressing a multitude of targets.

### Involvement of s479 in a transporter regulation network

The common theme reflected in the functions assigned to the proteins influenced by s479 reveals a network of transporters and transport related proteins. Amongst the proteins exclusively present in the s479 deletion strain are six components of ABC transporters including proteins encoded by the *znu* operon, which is regulated by s479 on the transcriptional level. The transported substances besides zinc are phosphate and iron. All of them are influx transporters and suggest a role of s479 in regulating the cellular network of metal ion and phosphate transporters. Taking into consideration proteins, that are differentially expressed with log_2_ fold change<2, this is even more pronounced; 13 components of ABC transporters accumulated upon loss of s479 (Supplementary Table 2) including zinc, iron, phosphate and peptide substrates. Therefore, a potential role for s479 might be in regulating metallostasis. Cross-talk between the regulation of metal ion concentrations can be seen in bacteria for instance for the regulatory networks of transcription factors ZuR regulating zinc response and FuR regulating iron homeostasis in *Caulobacter crescentus* (Mazzon et al., 2014) or even within transport itself e.g. *Yersinia pestis* possesses a second Zn^2+^ transporter that engages components of the yersiniabactin (Ybt) siderophore-dependent transport system for iron (Bobrov et al., 2017). Further work is needed, to unravel, whether those translational effects are mediated directly or through secondary effects of other members of the regulon e.g. the translation initiation factor aIF-5A. But direct effect of s479 is plausible for at least a subset of targets as for 31 of the proteins on/off regulated in the s479 deletion strain, a site for physical interaction between the mRNA and the sRNA regulator could be predicted. Interestingly, the majority of interaction sites all map to the same part of the s479 sequence already predicted for the s479::*znu*C1 interaction (Figure 7, 8), which was confirmed as direct contact by gel shift analysis. But as it is not yet resolved entirely how translation initiation ensues on the mostly leaderless transcripts of *H. volcanii* (Babski et al., 2016), future work is needed to unravel how translational control by small RNAs is exerted in this archaeal species. Interestingly, despite the large regulon of s479 especially on protein level, deletion is not deleterious for the cell. Only under high zinc concentrations the deletion strain Δ*s479* shows substantial growth defects. This hints at an interdependent network of regulatory mechanisms that might involve other sRNAs or translational regulators balancing the cost of individual gene losses.

### The CRISPR-Cas connection

The s479 RNA is encoded between a CRISPR locus and the *cas* gene cluster. In addition, the s479 primary transcript contains a sequence highly similar to the CRISPR RNA repeat sequences (Supplementary Figure 2). CRISPR RNAs are processed by the endonuclease Cas6 at these repeat sequences to yield the functional crRNAs (Figure 11, Supplementary Figure 2) (Maier et al., 2019). Northern experiments confirmed that cells without Cas6 cannot generate mature s479. However, a 100 nucleotide RNA is still present and seems to be sufficient for regulating the *znu* mRNA.

**Figure 11.**
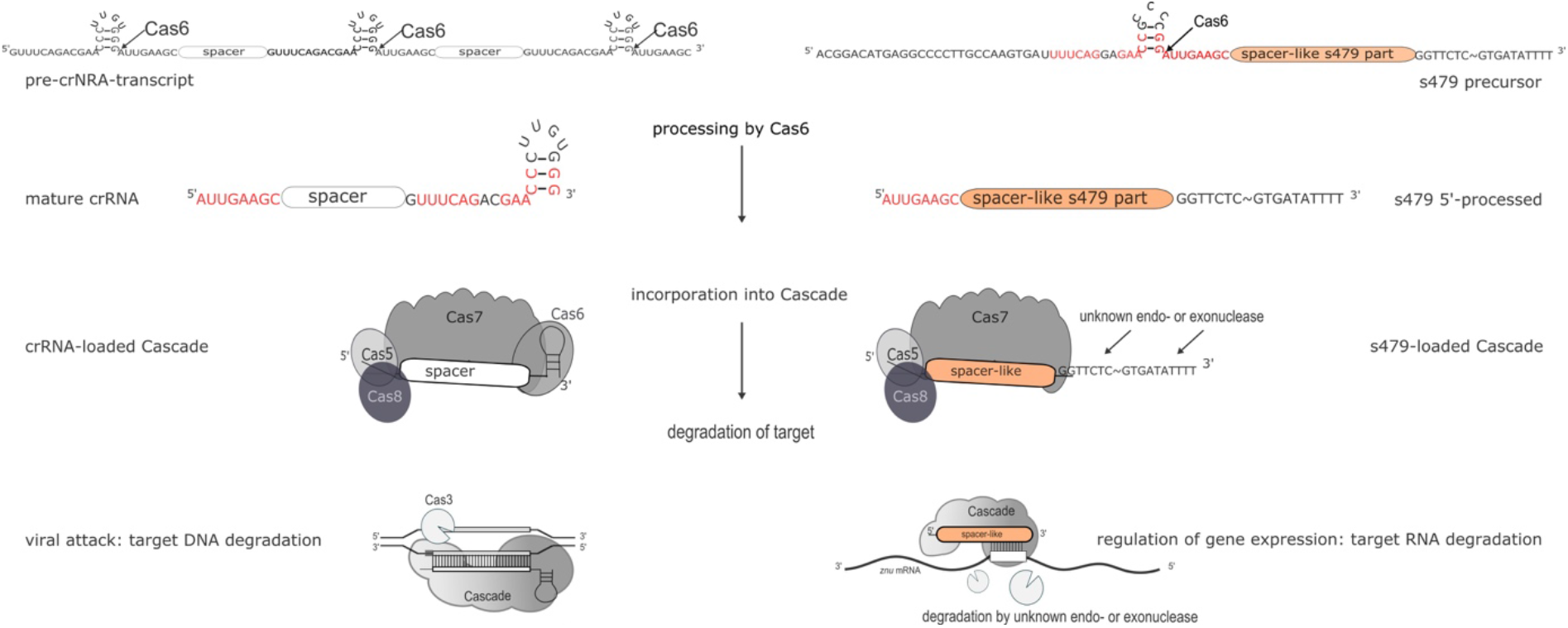
Maturation and function of crRNAs and s479. CRISPR arrays are transcribed into long precursors that are processed by Cas6 to yield the mature short crRNAs (left side). The mature crRNA is then loaded into the Cascade complex and guides Cascade to the invader to degrade the invader DNA. According to data presented here, s479 is also transcribed into a precursor RNA and processed by Cas6. s479 seems to be bound by Cascade via the crRNA-like 5’ handle and s479 might guide Cascade to the target *znu*C1 mRNA triggering its degradation.

In addition, cells without Cas7 and thereby without Cascade contain neither the mature s479 nor the 100 nucleotide RNA (Figure 10), suggesting that s479 is bound and protected by Cascade. The *znu* RNA is clearly up-regulated in a Δ*cas7* strain similar to the up-regulation in the Δ*s479* strain. The fact that the mature s479 contains a typical crRNA 5’ handle and crRNAs are bound to Cascade via the conserved 5’ handle makes it even more likely that s479 is bound by Cascade. Such a dependence and interaction of s479 with Cas6 and Cascade, respectively, would be the first example for a non-crRNA bound by a type I-B Cascade used for internal gene expression regulation, pointing to a connection between the evolutionary origin of this sRNA as a drift from the CRISPR-Cas machinery, evolved to control gene expression at the mRNA level. A similar mode of gene regulation by Cas proteins has so far only been shown in detail for type II systems (Dugar et al., 2018; Ratner et al., 2019). In *Campylobacter jejunii* for instance Cas9 is guided by a crRNA to mRNA targets, inducing RNA cleavage by Cas9 thereby regulating expression of these genes (Dugar et al., 2018).

Further work will show whether s479 guides Cascade to the *znu*C1 mRNA and triggers degradation of the mRNA (Figure 11).

## Conclusion

s479 supports a role for sRNAs as substantial regulatory players within the metabolic networks of *H. volcanii* especially in regulating metabolite availability as s479 seems to harmonize the abundance of several influx transporters of the ABC-type regulating the zinc, iron, peptide and phosphate flux of the cell. We demonstrate the direct interaction of s479 to its target *znu*C1 mRNA. Furthermore our data show that s479 is linked to the CRISPR-Cas system and might act together with Cascade to regulate zinc transport proteins.

## Materials and Methods

### Strains and Growth conditions

Strains and oligonucleotides used in this study are listed in Supplementary Table 3 and 4. A detailed description of the media used can be found in the Supplementary Data. Cloning procedures were performed using *E. coli* strain DH5α and standard culture (aerobically, 37°C, 2YT media) as well as molecular biological techniques (Sambrook, 2006). *H. volcanii* strains were grown aerobically at 45°C in either YPC, Hv-Ca, Hv-MM with appropriate supplements (Allers et al., 2004; Duggin et al., 2015; de Silva et al., 2020). For in-depth comparison of growth, transcriptome and proteome, H66 (Δ*pyr*E2, Δ*leu*B) was used as wild type, since in the s479 deletion strain, the s479 gene is replaced by a tryptophan marker in the genome of the parent strain H119 (Δ*pyr*E2, Δ*trp*A, and Δ*leu*B). Thus Δ*s479* and H66 require both addition of uracil and leucine to media for growth (Allers et al., 2004; Jaschinski et al., 2014).

### Growth Experiments

Growth experiments were carried out in microtiter plates using a heated plate reader (Epoch 2 NS Microplate Spectrophotometer, BioTek Instruments). Strains H66 (wild type) and Δ*s479* were precultured in Hv-Ca medium supplemented with uracil to OD_650nm_= 0.4-0.7 and then diluted to OD_650nm_=0.05 and transferred to microtiter plates in triplicates. These were then cultured (aerobically, orbital shaking, 45 °C) while OD_650nm_ was measured every 30 min. Outer wells were filled with salt water as evaporation barriers (Jaschinski et al., 2014; Liao et al., 2016). For stress conditions, adjusted media preparations were used (see above "Strains and Growth conditions"). Doubling time (d [h]) and growth rate (μ [h^−1^]) were calculated as growth rate μ = (ln(x_t_) – ln (x_0_)) / (t – t_0_) and doubling time d= ln(2) / μ. Calculations were carried out separately for all replicates before calculating mean value and standard deviation. Phases of exponential growth were identified using fitted trendlines and corresponding R^2^-values (Supplementary Figure 5B. and 5C.)

### Northern Blot Analysis

For analysis of s479 transcripts (Figure 2) *Haloferax* strains H119 and Δ*s479* were cultivated in Hv-MM supplemented with leucin and uracil (for H119 tryptophan was also added); for detection of *znu* mRNAs (Figure 6) *Haloferax* strains H66 and Δ*s479* were grown in YPC. For detection of s479 transcripts in Figure 10 *Haloferax* strain H119 was cultivated in Hv-MM supplemented with leucin, tryptophan and uracil; deletion strains Δ*cas6* and Δ*cas7* were cultivated in YPC. For investigation of *znu* mRNAs in Cas protein deletion strains Δ*cas6* and Δ*cas7* (Supplementary Figure 4) H119, Δ*cas6* and Δ*cas7* were grown in YPC. TRIzol™ Reagent (Invitrogen™, ThermoFisher Scientific) or NucleoZOL™ (Machery and Nagel) was used to isolate total RNA from *H. volcanii* cells. Ten microgram total RNA was separated using an 1.5 % agarose (transcript size >500 nt) or 8 % denaturing polyacrylamide gel (PAGE) and then transferred to a nylon membrane (Biodyne® A, PALL or Hybond-N+, GE Healthcare). After transfer, the membrane of PAGE blots was hybridized with oligonucleotide s479spacerpart (primer sequences are listed in Supplementary Table 4) to detect the s479 transcript, the membrane was subsequently hybridised with an oligonucleotide against the 5S rRNA, both radioactively labelled with [γ-^32^P]-ATP via polynucleotide kinase treatment. Membranes of agarose blots were hybridized with a probe against *znu*C1 generated by PCR using primers "probe znuC1 Hvo_2398 fw/rev" and genomic DNA as template and the product was labelled using [α-^32^P]-dCTP and the random primed DNA labelling kit DECAprime™II (Invitrogen). In addition, the membrane was hybridised with a probe against the 16S rRNA. The probe was generated with PCR using primers 16Sseqf and 16Sseqrev and genomic DNA from *H. volcanii* as template. Using the DECAprime™II kit (Invitrogen) the PCR fragment was radioactively labelled with α-^32^P-dCTP. Oligonucleotide probes and PCR primers see Supplementary Table 4.

### Sample Preparation for transcriptome analysis and RNA-seq analysis

Three replicates of wild type (H66) and deletion strain (Δ*s479*) were cultured in Hv-Ca medium supplemented with uracil at 45°C and grown to OD_650nm_=0.6-0.7. Total RNA was isolated using NucleoZOL™ (Machery and Nagel) and RNA samples were sent to vertis Biotechnologie AG (Martinsried, Germany) for further treatment. Total RNA was treated with T4 Polynucleotide Kinase (NEB) and rRNA depleted using an in-house protocol and cDNA library preparation was preceded by ultrasonic fragmentation. After 3’ adapter ligation, first-strand cDNA synthesis was performed using 3’-adapter primer and M-MLV reverse transcriptase. After cDNA purification, the 5’ Illumina TruSeq sequencing adapter was ligated to the 3’ end of the antisense cDNA and the sample amplified to 10-20 ng/μl using a high-fidelity DNA polymerase. Finally, cDNA was purified using the Agencourt AMPure XP kit (Beckman Coulter Genomics), samples were pooled (equimolar), the pool size fractionated (200 – 550 bp) by preparative agarose gel electrophoresis and sequenced on an Illumina NextSeq 500 system using 1×75 bp read length. TruSeq barcode sequences which are part of the 5’ TruSeq sequencing adapter are included in Supplementary Table 4. Sequencing reads are deposited at the European Nucleotide Archive (ENA) under the study accession number PRJEB41379. For data analysis, reads were mapped to the genome using bowtie2 (version2.3.4.1) with the „--very-sensitive” option and defaults otherwise (Langmead et al., 2009). Then reads per feature were counted using featureCounts (version 1.6.4) and analysed for differential expression with DeSeq2 (version 1.2.11) (Liao et al., 2014; Love et al., 2014).

### Sample Preparation for MS/MS Analysis

Three biological replicates (250 ml) of wild type (H66) and deletion strain (Δ*s479*) were cultivated in Hv-Ca media supplemented with uracil, 45°C and grown to OD_650nm_= 0.6-0.74. Cells were harvested and washed in enriched PBS buffer (2.5 M NaCl, 150 mM MgCl_2_, 1× PBS (137 mM NaCl, 2.7 mM KCl, 8 mM Na_2_HPO_4_, 2 mM K_2_HPO_4_, pH 7.4). After cell lysis by ultrasonication in 10 ml lysis buffer (1 M NaCl, 100 mM Tris/HCl, pH 7.5, 1 mM EDTA, 10 mM MgCl_2_, 1 mM CaCl_2_, 13 μl/ml protease inhibitor mixture (Sigma)) cytosolic and membrane fraction were separated by ultracentrifugation at 100.000 x g and treated as separate samples. The cytosolic protein sample was directly used for 1D SDS PAGE whereas the pelleted membrane protein fraction was solubilized in 2 ml HTH buffer ((6 M Thiourea/2 M Urea); 10 min 37°C; 10 min 37°C ultrasonication). 20 μg of both samples were separated by 1D SDS PAGE and in gel digested as previously described (Bonn et al., 2014). Briefly, Coomassie stained gel lanes were cut resulting in ten gel pieces per sample before gel pieces were cut into smaller blocks and transferred into low binding tubes. Samples were destained and dried in a vacuum centrifuge before being covered with trypsin solution. Digestion was carried out at 37 °C overnight before peptides were eluted in water by ultrasonication. The peptide-containing supernatant was transferred into a fresh tube, desiccated in a vacuum centrifuge and peptides were resolubilized in 0.1% (v/v) acetic acid for mass spectrometric analysis.

### MS/MS Analysis

LC-MS/MS analyses were performed on an LTQ Orbitrap Velos Pro (ThermoFisher Scientific, Waltham, Massachusetts, USA) using an EASY-nLC II liquid chromatography system. Tryptic peptides were subjected to liquid chromatography (LC) separation and electrospray ionization-based mass spectrometry (MS) applying the same injected volumes in order to allow for label-free relative protein quantification. Therefore, peptides were loaded on a self-packed analytical column (OD 360 μm, ID 100 μm, length 20 cm) filled with 3 μm diameter C18 particles (Dr. Maisch, Ammerbuch-Entringen, Germany) and eluted by a binary nonlinear gradient of 5 - 99 % acetonitrile in 0.1 % acetic acid over 86 min with a flow rate of 300 nL/min. For MS analysis, a full scan in the Orbitrap with a resolution of 30,000 was followed by collision-induced dissociation (CID) of the twenty most abundant precursor ions. MS2 experiments were acquired in the linear ion trap.

### MS Data Analysis

Database search was performed with MaxQuant 1.6.17.0 against a *H. volcanii* database (Jevtić et al., 2019) containing 4106 entries. Max Quant’s generic contamination list as well as reverse entries were added during the search. The following parameters were applied: digestion mode, trypsin/P with up to 2 missed cleavages; variable modification, methionine oxidation and N-terminal acetylation, and maximal number of 5 modifications per peptide; activated LFQ option with minimal ratio count of 2 and ‘match-between runs’ feature. The false discovery rates of peptide spectrum match and protein level were set to 0.01. A protein was considered to be identified if two or more unique peptides were identified in a biological replicate. Only unique peptides were used for protein quantification.

The comparative proteome analyses based on MaxQuant LFQ values were performed separately for cytosolic and membrane protein samples. Proteins were considered to be quantified if a quantitative value based on at least two unique peptides was available in at least two biological replicates. LFQ values as proxy for protein abundance were used for statistical analysis. A Student’s t-tests was performed to analyse changes in protein amounts between wild type and mutant. Proteins with significantly changed amount exhibited a p-value < 0.01 and an average log_2_ fold change >|0.8|

### Electrophoretic mobility shift assay

For electrophoretic mobility shift assays (EMSA) RNAs were obtained from Biomers (Biomers, Ulm, Germany) (sequences are listed in Supplementary Table 4). The s479 RNA was labelled at the 3’ end using [α-^32^P]pCp and T4 RNA ligase (Fermentas, Thermo Fisher Scientific). For the EMSA in Figure 9, 100 cps labelled s479-RNA was mixed with 50, 100 or 200 pmol the unlabelled *znu*C1 RNA fragment encompassing the interaction site 2 (Figure 7D.). For the EMSA in Supplementary Figure 3A., 100 cps labelled s479 RNA was mixed with 50 pmol unlabelled *znu*C1 RNA encompassing the interaction site 2 (Figure 7D.), in addition 0, 50, 200 or 400 pmol of unlabelled s479 were added. For the EMSA in Supplementary Figure 3B., 100 cps labelled s479 RNA was mixed with 50 pmol unlabelled *znu*C1 RNA encompassing the interaction site 2 (Figure 7D.) or 50, 200 or 400 pmol of unlabelled *znu*C1 RNA mutant, which has the s479 interaction site deleted (Figure 7D.) All reactions were performed in 20 μl reaction volume containing 10 mM Tris-Cl pH 7.5, 5 mM MgCl_2_ and 100 mM KCl. After incubation at 37°C for 30 min 1 μl 50% (vol/vol) glycerol containing 0.1% (w/vol) bromphenol blue was added and samples separated on a native 8% (w/vol) polyacrylamide gel at 4°C which was subsequently analysed by autoradiography.

### In silico target site prediction

To predict target sites of s479 *in silico*, we applied IntaRNA (version 3.2.0) (Mann et al., 2017). For prediction of potential s479 interaction sites, we used the s479 sequence corresponding to pHV4: 207,716-770. This corresponds to the start point of the potential spacer sequence of the shortened s479 versions (Figure 3). The spacer is the sequence located downstream of the 5’ handle sequence within crRNAs. In crRNAs, the spacer sequence is the sequence used for target recognition. Therefore we chose this part of the sequence for analysis and set the spacer length to 55 nt. IntaRNA was used with default settings for prediction of the s479::*znu*C1 interaction sites. For prediction of the interaction sites on the proteome-targets, we first used default settings and then also included predictions for a seed sequence of five nucleotides as this is the increment of protein contacts seen for spacer sequences within Cascade complexes (Maier et al., 2019).

## Data availability statement

The transcriptome data were deposited in the European Nucleotide Archive (ENA) at EMBL-EBI under study accession number: PRJEB41379.

Proteome data was deposited to the ProteomeXchange Consortium via the PRIDE partner repository (Vizcaíno et al., 2016) with the dataset identifier PXD022750.

## Author contributions

PM, LKM, SM, CH, JB and BV did the experiments, PM, LKM, AM, SM, DB and BV performed data curation. AM conceptualised the project. LKM, PM, and AM wrote the original draft. SM, JB, DB, BV, LKM and AM reviewed and edited the draft, DB, BV, and AM provided the resources and funding. All authors contributed to the article and approved the submitted version.

## Acknowledgments

We thank Susanne Schmidt for expert technical assistance. We would like to thank the reviewers for their constructive comments to improve the manuscript.

## Footnotes

1 For targets without a predicted TSS (Babski et al., 2016) or targets located in operons, we included 100 nt upstream of the first coding nucleotide.

