## SupplementaryData for "A small RNA is linking CRISPR-Cas and zinc transport"

**Supplementary data**


| | | |
|---|---|---|
| Supplementary Figures | page | 2 |
| Supplementary Methods and Materials | page | 8 |
| Supplementary Tables | page | 9 |

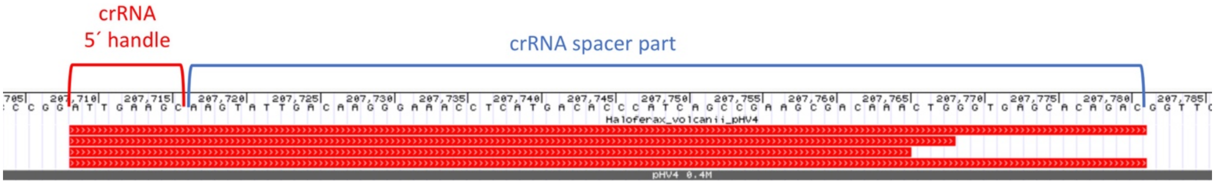


**Supplementary Figure 1. RNA-seq analysis of small RNAs reveals shortened s479 variants.** Sequencing data from an RNA-seq study dedicated to identification of small RNAs revealed reads mapping to the s479 locus all starting at position pHV4: 207,708. The reads extend to s479 length variants of 57, 60 and 73 nt. The sequence that is identical to the crRNA 5´ handle is indicated with a red bracket; this is the part that interacts with the Cas5 protein in the Cas protein complex Cascade. The sequence downstream of the 5´ handle corresponds to the crRNA spacer sequence and is indicated with a blue bracket.

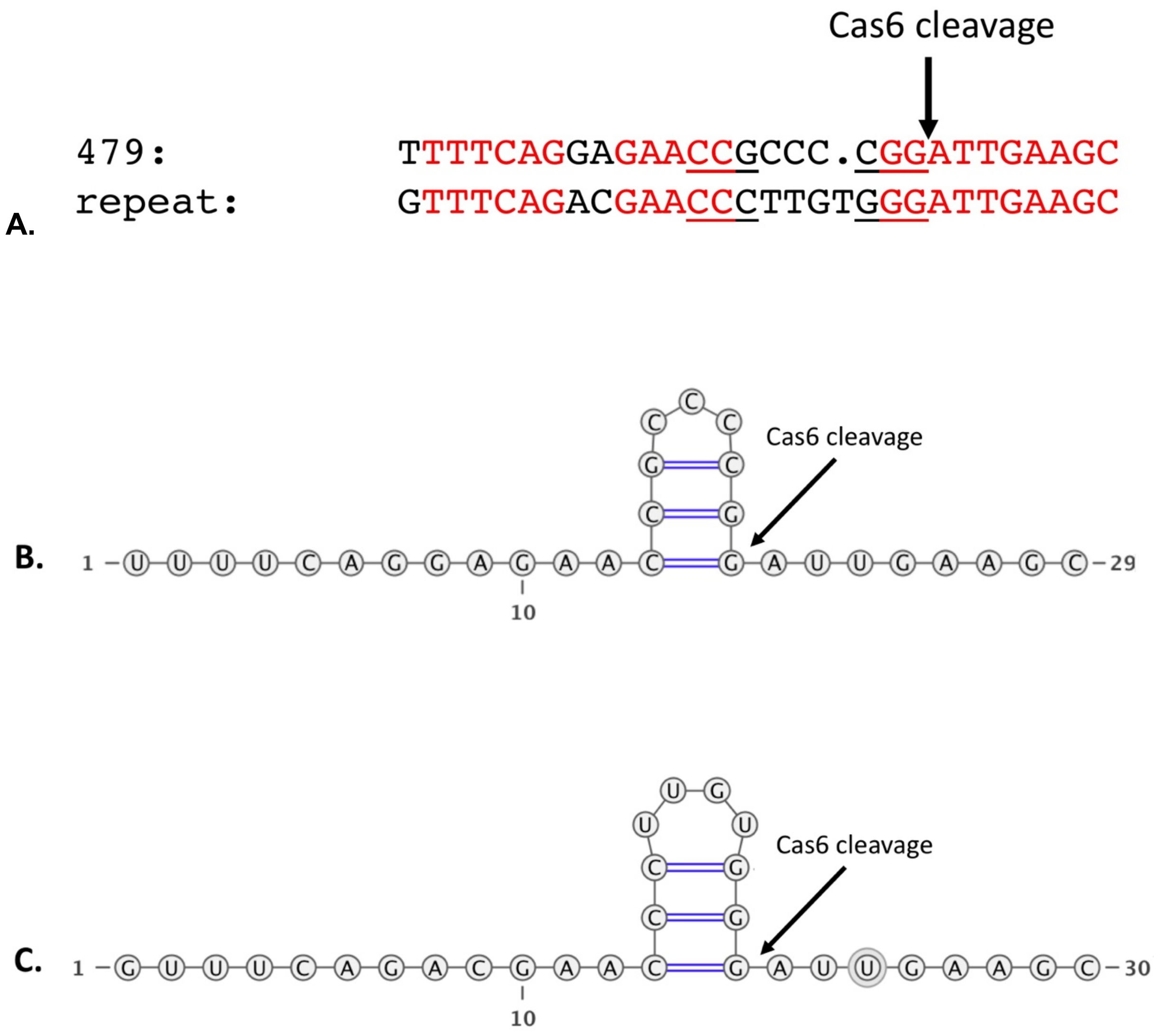


**Supplementary Figure 2. Similarity of the s479 precursor and crRNA precursors (pre-crRNA). A.** The s479 coding sequence of the precursor is highly similar (red nucleotides) to the repeat sequence of the CRISPR locus P1 flanking it upstream. The s479 precursor repeat-like sequence shows the potential to fold into a stem-loop structure (flanking nucleotides underlined). The Cas6 processing site is indicated by an arrow. **B.** The resulting stem-loop of the s479 precursor lies just upstream of the future handle sequence (AUUGAAGC) as seen for *H. volcanii* crRNAs (Maier et al., 2019). If the s479 precursor is processed by Cas6 like the CRISPR pre-crRNA, an endonucleolytic cut at the base of this stem-loop (shown by an arrow) yields the 5´-handle as seen in mature crRNAs of locus P1 and as seen in RNA-Seq. **C.** The structure of the CRISPR crRNA is shown for comparison.

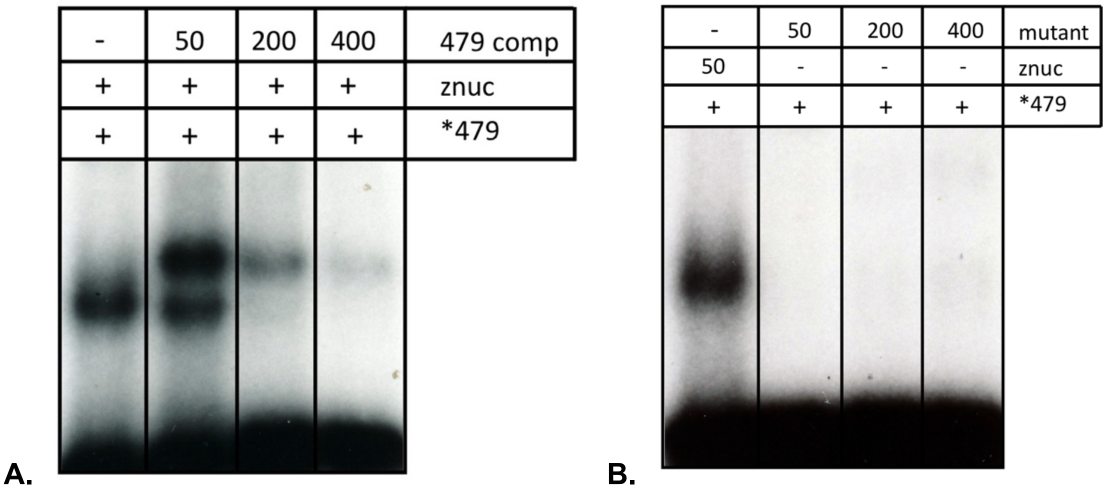


**Supplementary Figure 3. Gel shift experiments with s479.** Labelled s479 was incubated with different RNAs. Reactions were loaded onto a non-denaturing polyacrylamide gel to separate the molecules.

**A. Addition of unlabelled sRNA s479 competes with binding of radioactively labelled sRNA s479 to *znu*C1 RNA.** Radioactively labelled s479 was incubated with the *znu*C1 mRNA fragment (as shown in Figure 9.). Increasing amounts of unlabelled s479 were added (lanes 479 comp: 50, 200 and 400 pmol). The unlabelled s479 clearly competes with binding of the labelled s479. Addition of unlabelled s479 results in an additional bigger complex, which runs slower in the gel. This might be due to binding of more than one s479 molecules to the *znu*C1 mRNA.

**B. s479 does not bind to a mutant *znu*C1 RNA without the binding site.** Radioactively labelled s479 was incubated with a *znu*C1 RNA mutant, which has the s479 interaction site deleted. s479 does not bind to the mutant *znu*C1 RNA. Different amounts of the *znu*C1 mutant RNA were incubated with the radioactively labelled s479 (lanes mutant: 50 pmol, 200 pmol and 400 pmol). As positive control the radioactively labelled s479 was incubated with 50 pmol wild type *znu*C1 RNA (as shown in Figure 9) (lane znuc).

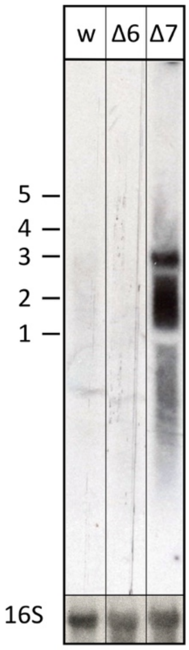


**Supplementary Figure 4. Transcript levels of the *znu* operon in wild type and Cas protein deletion strains.** We compared the transcript levels of *znu* in total RNA of wild type H119 (lane w), Δ*cas6* (lane Δ6) as well as Δ*cas7* cells (lane Δ7) using northern blot analysis. The probe used for hybridisation in the upper panel is located in the central part of the *znuC1* coding sequence (Figure 5). Signals at about 3,000 nucleotides correspond in length to the complete *znu* operon mRNA, signals at approximately 2,000 correspond in length to a bicistronic mRNA encompassing either *znu*A1 and *znu*C1 or *znu*C1 and *znu*B1. In the lower panel the blot was hybridised with a probe against the 16S rRNA. A size marker is given in kb at the left.

The wild type strain has almost no detectable *znu* mRNA, while high concentrations of the *znu* mRNA are visible in the Δ*s479* strain (Figure 6). Similar high concentrations are found in the Δ*cas7* strain confirming that Cas7 is not only important for a stable s479 as shown in Figure 10 but also for regulation of the *znu* mRNA. Interestingly, although Δ*cas6* is important for generating the mature s479 (Figure 10), the 100 nucleotide s479 intermediate seems sufficient for the regulation of the *znu* mRNA, since *znu* mRNA is not detected in a Δ*cas6* strain.

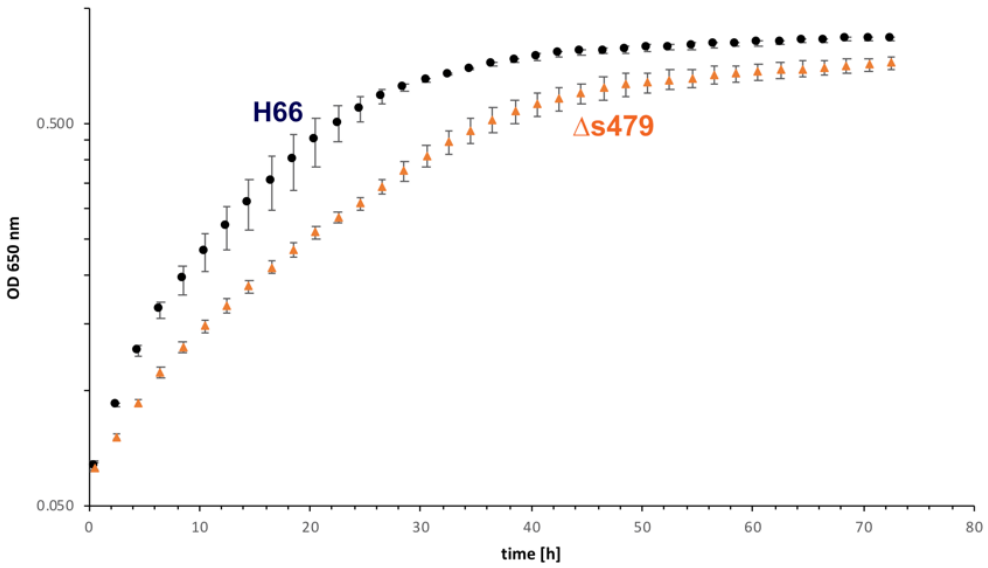

OD 650 nm
0.500
0.050
H66
Δs479
0
10
20
30
40
50
60
70
80
time [h]


**A.**

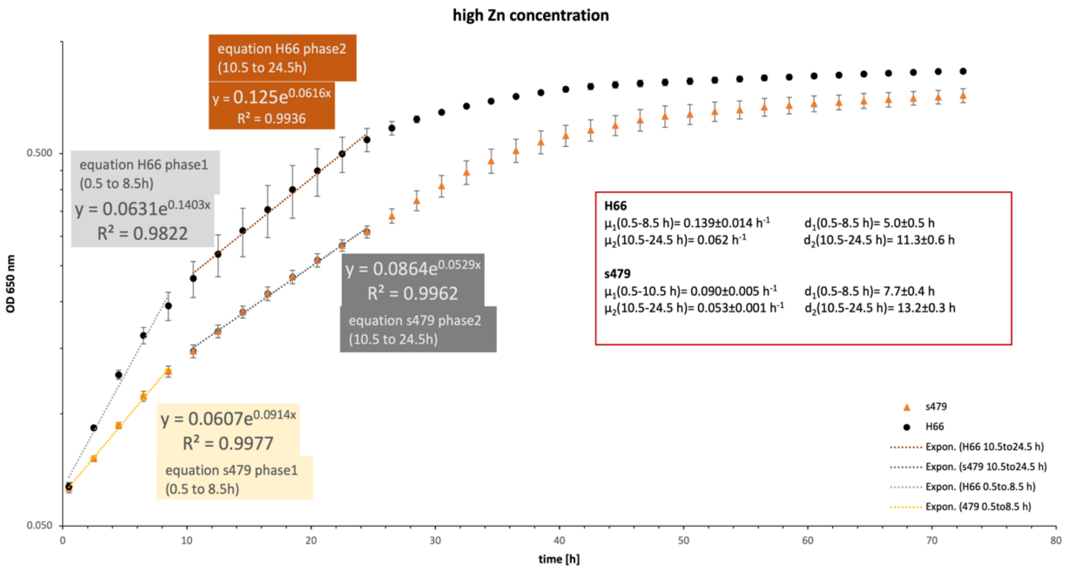

high Zn concentration
equation H66 phase2
(10.5 to 24.5h)
y = 0.125e^0.0616x
R² = 0.9936
equation H66 phase1
(0.5 to 8.5h)
y = 0.0631e^0.1403x
R² = 0.9822
y = 0.0864e^0.0529x
R² = 0.9962
equation s479 phase2
(10.5 to 24.5h)
y = 0.0607e^0.0914x
R² = 0.9977
equation s479 phase1
(0.5 to 8.5h)
H66
μ1(0.5-8.5 h)= 0.139±0.014 h-1
d1(0.5-8.5 h)= 5.0±0.5 h
μ2(10.5-24.5 h)= 0.062 h-1
d2(10.5-24.5 h)= 11.3±0.6 h
s479
μ1(0.5-10.5 h)= 0.090±0.005 h-1
d1(0.5-8.5 h)= 7.7±0.4 h
μ2(10.5-24.5 h)= 0.053±0.001 h-1
d2(10.5-24.5 h)= 13.2±0.3 h
s479
H66
Expon. (H66 10.5to24.5 h)
Expon. (s479 10.5to24.5 h)
Expon. (H66 0.5to.8.5 h)
Expon. (479 0.5to8.5 h)
OD 650 nm
0.500
0.050
0
10
20
30
40
50
60
70
80
time [h]


**B.**

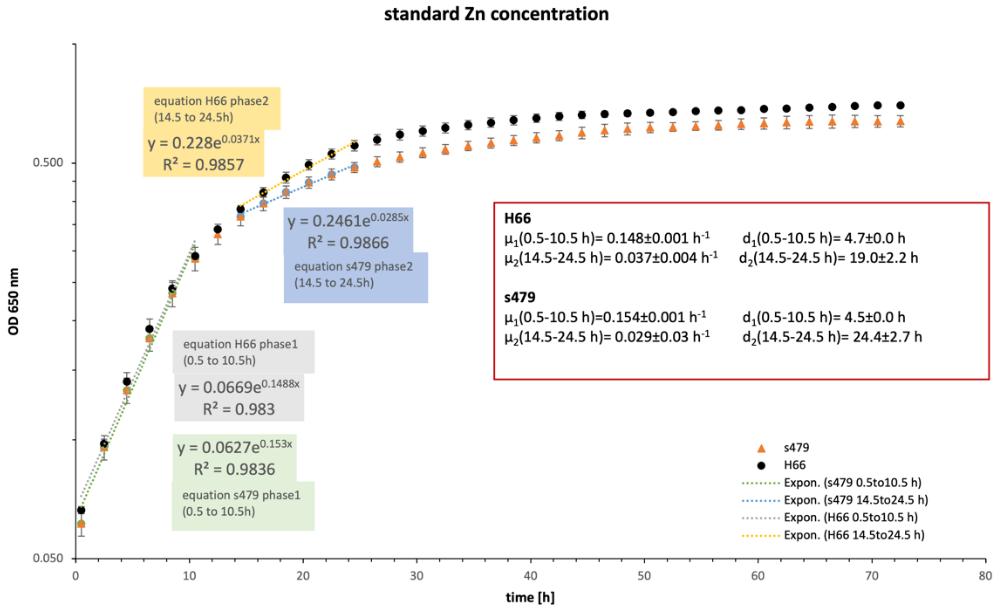


**C.**

**Supplementary Figure 5. Growth of wild type and Δ*s479* strains in different media.**

**A. Growth in medium with high zinc concentrations.** The deletion mutant has a delayed growth compared to the wild type strain. Growth curves show a diauxic growth behaviour, doubling times for phase 1 (0.5 h- 8.5 h) are 5 hours for wild type cells and 7.7 hours for the deletion strain. In phase 2 (10.5 h - 24.5 h) the doubling time for the wild type strain is 11.3 and for the deletion strain 13.2.

**B. Determination of doubling times and growth rates.** Slope analysis of the growth curves reveals a biphasic exponential phase. The trendline, line equation and R2 are given for phase 1 (0.5 h to 8.5 h) and phase 2 (10.5 h to 24.5 h) separately for each growth curve. The red box gives the calculated growth rate (µ) and doubling time (d) for each phase and strain as mean of the three replicates together with the respective standard deviation. Raw data and calculation are listed in Supplementary Table 6.

**C. Determination of doubling times and growth rates for growth in standard medium (growth curve shown in Figure 4).**
Slope analysis of the growth curves reveals a biphasic exponential phase. The trendline, line equation and R2 are given for phase 1 (0.5 h to 10.5 h) and phase 2 (14.5 h to 24.5 h) separately for each growth curve. The red box gives the calculated growth rate (µ) and doubling time (d) for each phase and strain as mean of the three replicates together with the respective standard deviation. Raw data and calculation are listed in Supplementary Table 5.

**Supplementary Material and Methods**

**Media composition**

Hv-YPC, Hv-Ca, and Hv-MM medium were used to grow *H. volcanii* (Allers *et al.*, 2004). Due to the sensitivity of *H. volcanii* to different ingredients Bacto yeast extract, casamino acids and peptone were used only from Oxoid. Unless otherwise stated, all components were autoclaved at 121 °C for 20 minutes.

**YPC.** For one liter of YPC liquid medium 600 ml 30% salt water (SW) (4.1 M NaCl, 148 mM $MgCl_2$, 142 mM $MgSO_4$, 94 mM KCl, 20 mM Tris/HCl pH 7.5), 300 ml double-distilled water and 99 ml 10x YPC stock solution (0.52% (w/v) Bacto yeast extract (Oxoid), 0.1% (w/v) peptone (Oxoid), 0.1% (w/v) casamino acids (Oxoid), adjust pH to 7.5 with KOH) were mixed and autoclaved. After cooling and before use 3 ml 1 M $CaCl_2$ (3 mM) was added.

**Hv-Ca.** For one liter of Hv-Ca medium, 600 ml SW, 225 ml double-distilled water and 30 ml 1M Tris-HCl/pH 7.5 were mixed and autoclaved. After cooling, 99 ml sterile-filtered 10x casamino acids (6% (w/v) casamino acids, 29.5 mM KOH.), 25.5 ml Hv-MM carbon source (10% sodium-DL-lactate, 9% succinic acid, 1% glycerol, adjust pH to 7.5 with NaOH), 12 ml Hv-MM salts (30 ml 1M $NH_4Cl$, 18 ml 1M $CaCl_2$, 6 ml trace elements, 18 ml double-distilled water); (trace elements: 0.0036 mg/ml $MnCl_2x4H_2O$, 0.0044 mg/ml $ZnSO_4x7H_2O$, 0.023 mg/ml $FeSO_4x7H_2O$, 0.0005 mg/ml $CuSO_4x5H_2O$), 1.95 ml 0.5 M $KPO_4$-buffer (83.4 ml 1 M $K_2HPO_4$, 16.6 ml 1 M $KH_2PO_4$, 100 ml double-distilled water) and 900 µl vitamin solution (1 mg/ml thiamine and 0.1 mg/ml biotin) were added to the medium. In addition, 1.02 ml (50 µg/ml) of sterile filtered uracil was added to the medium before use. To increase the zinc concentration for high-zinc medium, double the amount of $ZnSO_4$ was added in the trace elements. Please note that Hv-Ca medium contains a source of leucin.

**Hv-MM.** For one liter of Hv-MM medium, 600 ml of SW and 325 ml double-distilled water and 30 ml 1M Tris-HCl/pH 7.5 were autoclaved. After autoclaving, the following sterile filtered components were added, 25.5 ml Hv-MM carbon source (10% sodium-DL-lactate, 9% succinic acid, 1% glycerol, adjust pH to 7.5 with NaOH), 12 ml Hv-MM salts (30 ml 1 M $NH_4Cl$, 18 ml 1 M $CaCl_2$, 6 ml trace elements, 18 ml double-distilled water); (trace elements: 0.0036 mg/ml $MnCl_2x4H_2O$, 0.0044 mg/ml $ZnSO_4x7H_2O$, 0.023 mg/ml $FeSO_4x7H_2O$, 0.5 µg/ml $CuSO_4x5H_2O$), 1.95 ml 0.5 M $KPO_4$-buffer (83.4 ml 1 M $K_2HPO_4$, 16.6 ml 1 M $KH_2PO_4$, 100 ml double-distilled water) and 900 µl vitamin solution (9.6 ml thiamine 1 mg/ml, 1.2 ml biotin 1 mg/ml,) were added to the medium. In addition, if required 1.02 ml (50 µg/ml) of sterile filtered uracil, 5.1 ml (50 µg/ml) of sterile filtered tryptophan and 5.1 ml (50 µg/ml) of sterile filtered leucine was added to the medium before use.

**Supplementary Table 1. Transcripts that are up- and down-regulated in the Δ*s479* strain.** The $log_2$ fold change (column $log_2$) deletion vs. wild type is given alongside the HVO- gene number (column gene), p-value (column p.adj) and gene product name (column product). Green and red: up- and down-regulation, respectively.

| gene | product | $log_2$ | p.adj |
|---|---|---|---|
| s479 | sRNA s479 | -4.30 | 2.70E-31 |
| HVO_B0158 | hypothetical protein | -1.36 | 0.0439214 |
| HVO_0943 | ba3-type terminal oxidase subunit CbaD | -1.30 | 0.0083068 |
| HVO_0930 | hypothetical protein | -1.29 | 0.0351432 |
| HVO_B0371 | aldehyde dehydrogenase, AldH1 | 1.18 | 0.0227347 |
| HVO_2395 | ATP:cob(I)alamin adenosyltransferase, PduO | 1.29 | 0.0439214 |
| HVO_B0248 | short-chain family oxidoreductase | 1.47 | 0.0439214 |
| HVO_B0071 | alcohol dehydrogenase, Adh2 | 1.53 | 0.0164927 |
| HVO_2469 | SNF family transport protein | 1.54 | 0.0017482 |
| HVO_2397 | ABC-type transport system periplasmic substrate-binding protein (probable substrate zinc), ZnuA1 | 1.81 | 5.62E-05 |
| HVO_2399 | ABC-type transport system permease protein (probable substrate zinc), ZnuB1 | 1.98 | 3.15E-07 |
| HVO_2400 | small CPxCG-related zinc finger protein | 2.08 | 1.73E-08 |
| HVO_2398 | ABC-type transport system ATP-binding protein (probable substrate zinc), ZnuC1 | 2.11 | 1.58E-07 |
| HVO_2396 | glutaredoxin, Grx4 | 2.16 | 6.51E-07 |
| HVO_2401 | glycine cleavage system protein P beta subunit, GcvP2 | 3.27 | 1.66E-19 |

**Supplementary Table 2. Proteins that accumulated or were depleted in the $\Delta$*s479* strain.** Differential proteome analysis comparing the s479 deletion strain and wild type H66 was used determining the $\log_2$ fold change (column on/off/$\log_2$). If detected only in the deletion strain the protein is listed as "on", if detected only in the wild type strain the protein is listed as "off". Additionally, HVO-gene numbers (column gene) encoding the proteins are given alongside the gene product (column gene product) and Kegg/COG assignments (columns Kegg pathway/ COG) informing on the metabolic pathway(s). The MS sample the protein was measured in (column S; c: cytoplasmic, m: membrane) and the predicted localization (column local; C: cytoplasm, M: membrane) are also included. Orange: off or depleted in $\Delta$*s479* samples. Green: on or enriched in $\Delta$*s479* samples. Italic: association with tryptophan metabolism.

| S | gene product | gene | loc | cn/ off/ $\log_2$ ($\Delta$s479 / WT) | Kegg pathway | COG |
|---|---|---|---|---|---|---|
| c | A-type ATP synthase subunit E | HVO_0313 | C | off | hvo00190 - Oxidative phosphorylation; hvo00680 - Methane metabolism; | |
| c | acid phosphatase | HVO_1314 | C | off | hvo00230 - Purine metabolism; hvo00240 - Pyrimidine metabolism; hvo00760 - Nicotinate and nicotinamide metabolism; hvo01110 - Biosynthesis of secondary metabolites; | |
| c | glutamate dehydrogenase | HVO_1451 | C | off | hvo00250 - Alanine, aspartate and glutamate metabolism; hvo00330 - Arginine and proline metabolism; hvo00910 - Nitrogen metabolism; | |
| *c* | *N-(5phosphoribosyl)anthranilate isomerase* | *HVO_2455* | *C* | *off* | *hvo00400 - Phenylalanine, tyrosine and tryptophan biosynthesis /hvo01110 - Biosynthesis of secondary metabolites* | |
| c | 5,10-methylenetetrahydromethanopterin reductase | HVO_1937 | C | off | hvo00680 - Methane metabolism; hvo01120 - Microbial metabolism in diverse environments; | |
| m | cobalamin 5-phosphate synthase | HVO_0588 | C/M | off | hvo00860 - Porphyrin and chlorophyll metabolism | |
| c | 50S ribosomal protein L39.eR | HVO_0115 | C | off | hvo03010 - Ribosome; | |
| c | endonuclease IV | HVO_0573 | C | off | vo03410 - Base excision repair | |
| c | acyl-CoA synthetase | HVO_A0551 | C | off | | COG0318 Code IQ Acyl-CoA synthetases (AMP-forming)/AMP-acid ligases II |
| c | cytochrome P450 | HVO_1558 | C | off | | OG2124 Code Q Cytochrome P450 |
| c | deoxyhypusine synthase | HVO_2297 | C | off | | OG1899 Code O Deoxyhypusine synthase |
| c | DnaJ domain-containing protein | HVO_1040 | C | off | | COG0484 Code O DnaJ-class molecular chaperone with C-terminal Zn finger domain |
| c | ExsB protein | HVO_1716 | Unknown | off | | COG0603 Code R Predicted PP-loop superfamily ATPase |
| c | hypothetical protein | HVO_2024 | C | off | | |
| c | hypothetical protein | HVO_2640 | C | off | | |

| | | | | | | |
|---|---|---|---|---|---|---|
| c | colbalt chelase thioredoxin | HVO_B0054 | C | off | | COG2138 Code S Uncharacterized conserved protein |
| c | PQQ repeat protein | HVO_B0138 | C | off | | COG1520 Code S FOG: WD40-like repeat |
| c | Xaa-Pro aminopeptidase, M24 family protein | HVO_0414 | Unknown | off | | COG0006 Code E Xaa-Pro aminopeptidase |
| c | 3-dehydroquinate synthase | HVO_0822 | C | on | | COG0371 Code C Glycerol dehydrogenase and related enzymes |
| c | NADH dehydrogenase-like complex subunit I | HVO_0982 | C | on | hvo00190 - Oxidative phosphorylation; | |
| m | glycerol-3-phosphate dehydrogenase subunit B | HVO_1539 | C/M | on | hvo00564 - Glycerophospholipid metabolism | |
| m | ABC-type transport system periplasmic substrate-binding protein (probable substrate iron-III) | HVO_B0198 | Unknown | on | hvo02010 - ABC transporters | |
| m | ABC-type transport system periplasmic substrate-binding protein (probable substrate iron-III) | HVO_B0047 | Extra-cellular | on | hvo02010 - ABC transporters | |
| m | putative phosphate ABC transporter periplasmic substrate-binding protein | HVO_2375 | C/M | on | hvo02010 - ABC transporters | |
| m | putative zinc ABC transporter periplasmic substrate-binding protein | HVO_2397 | C/M | on | hvo02010 - ABC transporters | |
| m | putative iron-III ABC transporter periplasmic substrate-binding protein | HVO_1705 | Unknown | on | hvo02010 - ABC transporters; | |
| c | TATA-binding transcription initiation factor | HVO_B0382 | C | on | hvo03022 - Basal transcription factors; | |
| m | ATP-dependent RNA helicase/nuclease Hef | HVO_3010 | Unknown | on | | COG1111 Code L ERCC4-like helicases |
| m | flavoprotein reductase-like protein | HVO_0105 | C/M | on | | COG: COG1252 Code C NADH dehydrogenase, FAD-containing subunit |
| m | FxsA-like protein | HVO_2351 | C/M | on | | COG3030 Code R Protein affecting phage T7 exclusion by the F plasmid |
| m | hypothetical | HVO_0307 | Unknown | on | | |
| m | hypothetical | HVO_0129 | C/M | on | | |
| m | hypothetical | HVO_1010 | C/M | on | | |
| m | hypothetical | HVO_1307 | C/M | on | | |
| m | hypothetical | HVO_1850 | C/M | on | | |
| m | hypothetical | HVO_2513 | C/M | on | | |
| m | hypothetical | HVO_A0316 | C/M | on | | |
| m | Tat (twin-arginine translocation) pathway signalsequence domain protein | HVO_C0054 | Unknown | on | | |
| c | UspA domain protein | HVO_A0047 | C | on | | COG0589 Code T Universal stress protein UspA and related nucleotide-binding proteins |

| | | | | | | |
|---|---|---|---|---|---|---|
| c | UspA domain-containing protein | HVO_1198 | C | on | | COG0589 Code T Universal stress protein UspA and related nucleotide-binding proteins |
| m | cationic amino acid transporter | HVO_A0175 | C/M | **-3.5** | | COG0531 Code E Amino acid transporters |
| c | DNA polymerase D polymerase subunit DP2 | HVO_0065 | C | **-3.5** | hvo00230 - Purine metabolism;<br>hvo00240 - Pyrimidine metabolism | |
| c | PQQ repeat protein | HVO_B0052 | Unknown | **-2.7** | | COG1520 Code S FOG: WD40-like repeat |
| *c* | *tryptophan synthase subunit beta* | *HVO_0788* | *C* | **-2.2** | *hvo00260 - Glycine, serine and threonine metabolism; hvo00400 - Phenylalanine, tyrosine and tryptophan biosynthesis;<br>hvo01110 - Biosynthesis of secondary metabolites;* | |
| c | acyl-CoA dehydrogenase | HVO_1199 | C | -1.8 | | COG1960 Code I Acyl-CoA dehydrogenases |
| m | cytochrome d ubiquinol oxidase subunit II | HVO_0461 | C/M | -1.8 | hvo00190 - Oxidative phosphorylation | |
| m | hypothetical | HVO_B0359 | C/M | -1.8 | | |
| *c* | *tryptophan synthase subunit alpha* | *HVO_0789* | *C* | *-1.8* | *hvo00260 - Glycine, serine and threonine metabolism; hvo00400 - Phenylalanine, tyrosine and tryptophan biosynthesis;<br>hvo01110 - Biosynthesis of secondary metabolites; hvo00400 - Phenylalanine, tyrosine and tryptophan biosynthesis;<br>hvo01110 - Biosynthesis of secondary metabolites;* | |
| c | isopentenyl-diphosphate delta-isomerase | HVO_2506 | C | -1.8 | hvo00900 - Terpenoid backbone biosynthesis; hvo01110 - Biosynthesis of secondary metabolites; | |
| c | 50S ribosomal protein L1 | HVO_2757 | C | -1.8 | hvo03010 - Ribosome | |
| c | 50S ribosomal protein L15.eR | HVO_0561 | C | -1.8 | hvo03010 - Ribosome | |
| m | PQQ repeat-containing protein | HVO_2606 | Unknown | -1.8 | | COG1520 Code S FOG: WD40-like repeat |
| c | 3-ketoacyl-CoA thiolase | HVO_1914 | C | -1.7 | hvo00071 - Fatty acid metabolism; hvo00280 - Valine, leucine and isoleucine degradation; hvo00281 - Geraniol degradation; hvo000362 - Benzoate degradation; hvo00592 - alpha-Linolenic acid metabolism; hvo01110 - Biosynthesis of secondary metabolites; | |
| c | CbiG | HVO_B0059 | C | -1.7 | hvo00860 - Porphyrin and chlorophyll metabolism; | |
| m | hypothetical | HVO_1103 | C/M | -1.7 | | |
| c | hypothetical protein | HVO_2203 | Unknown | -1.7 | | |
| c | DNA N-glycosylase | HVO_1681 | C | -1.6 | hvo03410 - Base excision repair; | |
| c | propionyl-CoA carboxylase complex B chain | HVO_1447 | C | -1.6 | hvo00280 - Valine, leucine and isoleucine degradation; hvo00630 - Glyoxylate and dicarboxylate metabolism;<br>hvo00640 - Propanoate metabolism; hvo01120 - Microbial metabolism in diverse environments; | |
| m | heavy-metal transporting CPx-type ATPase | HVO_0940 | C/M | -1.6 | | OG2217 Code P Cation transport ATPase |
| m | SpoIVFB-type metallopeptidase | HVO_0285 | C/M | -1.5 | | OG1994 Code R Zn-dependent proteases |
| c | carbamoyl-phosphate synthase large subunit | HVO_2361 | C | -1.4 | hvo00240 - Pyrimidine metabolism; hvo00250 - Alanine, aspartate and glutamate metabolism; | |
| c | acyl-CoA synthetase | HVO_0894 | C | -1.4 | hvo00010 - Glycolysis / Gluconeogenesis; hvo00620 - Pyruvate metabolism; hvo00640 - Propanoate metabolism;<br>hvo00680 - Methane metabolism; hvo01110 - Biosynthesis of secondary metabolites | |

| | | | | | | |
|---|---|---|---|---|---|---|
| c | ATP-dependent DNA helicase | HVO_1333 | C | -1.4 | | COG1201 Code R Lhr-like helicases |
| c | L-fuculose phosphate aldolase | HVO_A0268 | C | -1.4 | hvo00051 - Fructose and mannose metabolism | |
| c | precorrin-3B C17-methyltransferase | HVO_B0057 | C | -1.3 | hvo00860 - Porphyrin and chlorophyll metabolism; | |
| c | diphthamide synthase subunit | HVO_1631 | C | -1.3 | | COG0411 Code E ABC-type branched-chain amino acid transport systems, ATPase component |
| m | NADH dehydrogenase-like complex subunit B | HVO_0979 | C/M | -1.3 | hvo00190 - Oxidative phosphorylation | |
| c | ornithine carbamoyltransferase | HVO_0041 | C | -1.3 | | COG0078 Code E Ornithine carbamoyltransferase |
| c | GTP cyclohydrolase III 1 | HVO_1284 | C | -1.3 | hvo00740 - Riboflavin metabolism | |
| m | ba3-type terminal oxidase subunit II | HVO_0944 | C/M | -1.3 | hvo00190 - Oxidative phosphorylation | |
| c | acyl-CoA synthetase | HVO_1374 | C | -1.3 | | COG0318 Code IQ Acyl-CoA synthetases (AMP-forming)/AMP-acid ligases II |
| m | Manganese transport protein mntH | HVO_2674 | C/M | -1.3 | | COG1914 Code P Mn2+ and Fe2+ transporters of the NRAMP family |
| m | putative dipeptides/oligopeptides ABC transporter ATP-binding protein | HVO_0627 | C/M | -1.3 | hvo02010 - ABC transporters | |
| c | glycosyltransferase AglI | HVO_1528 | C | -1.3 | | COG1215 Code M Glycosyltransferases, probably involved in cell wall biogenesis |
| *c* | *acyl-CoA dehydrogenase* | *HVO_0209* | *C* | *-1.3* | *hvo00071 - Fatty acid metabolism; hvo00310 - Lysine degradation; hvo00362 - Benzoate degradation; hvo00380 - Tryptophan metabolism; hvo01120 - Microbial metabolism in diverse environments;* | |
| m | FAD-linked oxidase domain-containing protein | HVO_1697 | C/M | -1.2 | | COG0277 Code C FAD/FMN-containing dehydrogenases |
| c | GTP-binding protein Era | HVO_3014 | C | -1.2 | | COG1100 Code R GTPase SAR1 and related small G proteins |
| c | cystathionine gamma-synthase | HVO_2946 | C | -1.2 | hvo00260 - Glycine, serine and threonine metabolism; hvo00270 - Cysteine and methionine metabolism; hvo00450 - Selenoamino acid metabolism; hvo00910 - Nitrogen metabolism; | |
| c | farnesyl-diphosphate farnesyltransferase | HVO_1139 | C | -1.2 | ; hvo01110 - Biosynthesis of secondary metabolites | |
| m | hypothetical | HVO_0453 | C/M | -1.2 | | |
| c | putative inositol-1(or 4)-monophosphatase / fructose-1,6-bisphosphatase, archaeal type | HVO_2857 | C | -1.2 | hvo00521 - Streptomycin biosynthesis; hvo00562 - Inositol phosphate metabolism; hvo01110 - Biosynthesis of secondary metabolites | |
| c | glutamate dehydrogenase | HVO_B0266 | C | -1.2 | hvo00250 - Alanine, aspartate and glutamate metabolism; hvo00330 - Arginine and proline metabolism; hvo00910 - Nitrogen metabolism; | |
| c | phosphoribosylamine--glycine ligase | HVO_1657 | C | -1.2 | hvo00230 - Purine metabolism; hvo01110 - Biosynthesis of secondary metabolites | |
| c | malate synthase | HVO_B0200 | C | -1.1 | | COG2301 Code G Citrate lyase beta subunit |
| c | DJ-1/PfpI/ThiJ superfamily protein | HVO_1073 | C | -1.1 | | COG0693 Code R Putative intracellular protease/amidase |
| *c* | *anthranilate phosphoribosyltransferase* | *HVO_2456* | *C* | *-1.1* | *hvo00400 - Phenylalanine, tyrosine and tryptophan biosynthesis; hvo01110 - Biosynthesis of secondary metabolites* | |
| c | MiaB-like tRNA modifying enzyme, archaeal-type | HVO_2605 | C | -1.1 | | COG0621 Code J 2-methylthioadenine synthetase |

| | | | | | | |
|---|---|---|---|---|---|---|
| m | PBS lyase HEAT-like repeat domain-containing protein | HVO_1020 | Unknown | -1.1 | | COG1413 Code C FOG: HEAT repeat |
| c | precorrin-3B C17-methyltransferase | HVO_B0058 | Unknown | -1.1 | hvo00860 - Porphyrin and chlorophyll metabolism; | |
| c | cobyric acid synthase CobQ | HVO_A0553 | C | -1.1 | hvo00860 - Porphyrin and chlorophyll metabolism; | |
| c | repair helicase | HVO_0415 | C | -1.1 | hvo03420 - Nucleotide excision repair; hvo03430 - Mismatch repair | |
| c | uridine phosphorylase | HVO_2614 | C | -1.1 | hvo00240 - Pyrimidine metabolism | |
| m | hypothetical | HVO_0941 | C/M | -1.0 | | |
| c | acyl-CoA synthetase | HVO_1585 | C | -1.0 | hvo00010 - Glycolysis / Gluconeogenesis; hvo00620 - Pyruvate metabolism; hvo00640 - Propanoate metabolism; hvo00680 - Methane metabolism; hvo01110 - Biosynthesis of secondary metabolites | |
| c | methylmalonyl-CoA mutase subunit A | HVO_0893 | C | -1.0 | hvo00280 - Valine, leucine and isoleucine degradation; hvo00640 - Propanoate metabolism; hvo01120 - Microbial metabolism in diverse environments; | |
| c | DNA binding protein | HVO_0662 | C | -1.0 | | OG1992 Code S Uncharacterized conserved protein |
| c | phosphoribosylaminoimidazole-succinocarboxamide synthase | HVO_2193 | C | -1.0 | hvo00230 - Purine metabolism; hvo01110 - Biosynthesis of secondary metabolites | |
| c | pyrroline-5-carboxylate reductase | HVO_1372 | C | -1.0 | hvo00330 - Arginine and proline metabolism; hvo01110 - Biosynthesis of secondary metabolites | |
| c | quinolinate synthetase complex, A subunit | HVO_2581 | C | -1.0 | hvo00760 - Nicotinate and nicotinamide metabolism | |
| c | possible polygalacturonase, putative | HVO_B0085 | C | -1.0 | | COG5434 Code M Endopolygalacturonase |
| m | hypothetical | HVO_1927 | C/M | -1.0 | | |
| c | ferredoxin--nitrite reductase | HVO_1788 | C | -1.0 | hvo00910 - Nitrogen metabolism; hvo01120 - Microbial metabolism in diverse environments | |
| c | aminotransferase class V | HVO_A0485 | C | -1.0 | | COG0075 Code E Serine-pyruvate aminotransferase/archaeal aspartate aminotransferase |
| c | FolD bifunctional protein | HVO_2865 | C | -1.0 | hvo00670 - One carbon pool by folate; hvo01120 - Microbial metabolism in diverse environments | |
| m | cyanide insensitive terminal oxidase chain cioA | HVO_0462 | C/M | -1.0 | hvo00190 - Oxidative phosphorylation | |
| c | diphosphomevalonate decarboxylase | HVO_1412 | C | -0.9 | hvo00900 - Terpenoid backbone biosynthesis; hvo01110 - Biosynthesis of secondary metabolites | |
| c | 50S ribosomal protein L10.eR | HVO_0484 | C | -0.9 | hvo03010 - Ribosome | |
| c | Fic protein family, putative | HVO_A0401 | C | -0.9 | | **COG:** COG3177 Code S Uncharacterized conserved protein |
| c | 50S ribosomal protein L18 | HVO_2545 | Unknown | -0.9 | hvo03010 - Ribosome | |
| c | ArcR family transcription regulator | HVO_A0266 | Unknown | -0.9 | ArcR transcriptional regulator | COG1414 Code K Transcriptional regulator |
| c | FAD binding domain | HVO_2580 | C | -0.9 | hvo00250 - Alanine, aspartate and glutamate metabolism; hvo00760 - Nicotinate and nicotinamide metabolism | |
| c | citrate synthase | HVO_0466 | C | -0.9 | hvo00020 - Citrate cycle (TCA cycle); hvo00630 - Glyoxylate and dicarboxylate metabolism; hvo01110 - Biosynthesis of secondary metabolites | |
| c | 30S ribosomal protein S4 | HVO_2783 | C | -0.9 | hvo03010 - Ribosome | |

| | | | | | | |
|---|---|---|---|---|---|---|
| c | hypothetical protein | HVO_2080 | Unknown | -0.9 | | |
| m | HTR-like protein | HVO_A0160 | C/M | -0.8 | | COG0642 Code T Signal transduction histidine kinase |
| c | aspartyl-tRNA(Asn) amidotransferase subunit A | HVO_1054 | C | -0.8 | hvo00970 - Aminoacyl-tRNA biosynthesis | |
| m | cation-transporting ATPase | HVO_0933 | C/M | -0.8 | | OG0474 Code P Cation transport ATPase |
| c | cell surface glycoprotein | HVO_2072 | Cellwall | -0.8 | | |
| *c* | *anthranilate synthase component I* | *HVO_2454* | *C* | *-0.8* | *hvo00400 - Phenylalanine, tyrosine and tryptophan biosynthesis; hvo01110 - Biosynthesis of secondary metabolites* | |
| c | deoxyhypusine synthase | HVO_B0182 | C | -0.8 | | COG1899 Code O Deoxyhypusine synthase |
| *c* | *tryptophanase* | *HVO_0009* | *C* | ***4.3*** | *hvo00380 - Tryptophan metabolism; hvo00910 - Nitrogen metabolism;* | |
| m | hypothetical | HVO_B0064 | C/M | **4.2** | | |
| c | putative zinc ABC transporter ATP-binding protein | HVO_2398 | C | **2.5** | hvo02010 - ABC transporters | |
| m | cox-type terminal oxidase subunit I | HVO_0907 | C/M | **2.4** | hvo00190 - Oxidative phosphorylation | |
| c | halocyanin | HVO_1228 | C | **2.4** | | COG3794 Code C Plastocyanin |
| c | hypothetical protein | HVO_0377 | Unknown | **2.4** | | |
| m | pantothenate permease | HVO_2324 | C/M | **2.1** | | COG0591 Code ER Na+/proline symporter |
| c | translation initiation factor aIF-5A | HVO_2300 | C | 2.0 | | COG0231 Code J Translation elongation factor P (EF-P)/translation initiation factor 5A (eIF-5A) |
| m | hypothetical | HVO_0229 | C/M | 1.8 | | |
| c | sugar nucleotidyltransferase | HVO_A0586 | C | 1.8 | hvo00520 - Amino sugar and nucleotide sugar metabolism | |
| c | coenzyme PQQ synthesis protein E-like protein | HVO_1121 | C | 1.8 | | COG0535 Code R Predicted Fe-S oxidoreductases |
| m | preprotein translocase Sec61 alpha subunit | HVO_A0174 | C/M | 1.8 | hvo03060 - Protein export; hvo03070 - Bacterial secretion system | |
| c | putative iron-III ABC transporter periplasmic substrate-binding protein | HVO_1705 | Unknown | 1.7 | hvo02010 - ABC transporters | |
| c | pyridine nucleotide-disulfide oxidoreductase, class II, putative | HVO_A0501 | Unknown | 1.7 | | COG2226 Code H Methylase involved in ubiquinone/menaquinone biosynthesis |
| c | UspA domain protein | HVO_A0496 | C | 1.7 | | COG0589 Code T Universal stress protein UspA and related nucleotide-binding proteins |
| c | hypothetical protein | HVO_A0633 | Unknown | 1.7 | | |
| m | anion permease | HVO_2996 | C/M | 1.7 | | COG0306 Code P Phosphate/sulphate permeases |
| c M S | ABC-type transport system periplasmic substrate-binding protein (probable substrate dipeptides/oligopeptides) | HVO_A0339 | Cellwall | 1.6 | hvo02010 - ABC transporters | |

| | | | | | | |
|---|---|---|---|---|---|---|
| c | hypothetical protein | HVO_2327 | C | 1.6 | | |
| m | preprotein translocase Sec61 subunit gamma | HVO_0718 | C/M | 1.6 | hvo03060 - Protein export | |
| c | dioxgenase | HVO_2662 | C | 1.6 | | COG0346 Code E Lactoylglutathione lyase and related lyases |
| c | cobyrinic acid ac-diamide synthase | HVO_0999 | C | 1.5 | | COG0857 Code R BioD-like N-terminal domain of phosphotransacetylase |
| *c* | *anthranilate phosphoribosyltransferase* | *HVO_2226* | *C* | *1.5* | *hvo00400 - Phenylalanine, tyrosine and tryptophan biosynthesis; vo01110 - Biosynthesis of secondary metabolites;* | |
| c | deoxyribodipyrimidine photolyase | HVO_2911 | C | 1.5 | | COG0415 Code L Deoxyribodipyrimidine photolyase |
| m | 1,4-dihydroxy-2-naphthoateoctaprenyltransferase | HVO_1462 | C/M | 1.5 | hvo00130 - Ubiquinone and other terpenoid-quinone biosynthesis; hvo01110 - Biosynthesis of secondary metabolites; | |
| m | A-type ATP synthase subunit I | HVO_0311 | C/M | 1.4 | hvo00190 - Oxidative phosphorylation; hvo00680 - Methane metabolism; | |
| m | hypothetical | HVO_1182 | C/M | 1.4 | | |
| m | glycosyltransferase AglD | HVO_0798 | C/M | 1.4 | | COG0463 Code M Glycosyltransferases involved in cell wall biogenesis |
| c | thioredoxin reductase | HVO_1758 | C | 1.4 | | COG0492 Code O Thioredoxin reductase |
| c | HpcH/HpaI aldolase family protein | HVO_2665 | C | 1.4 | | OG3836 Code G 2,4-dihydroxyhept-2-ene-1,7-dioic acid aldolase |
| m | putative dipeptides/oligopeptides ABC transporter permease | HVO_0629 | C/M | 1.4 | hvo02010 - ABC transporters | |
| m | hypothetical | HVO_1274 | C/M | 1.4 | | |
| m | xanthine/uracil permease family protein | HVO_0335 | C/M | 1.3 | | COG2252 Code R Permeases |
| m | dolichyl-phosphate-mannose-proteinmannosyltransfe rase | HVO_0975 | C/M | 1.3 | | COG4745 Code O Predicted m-bound mannosyltransferase |
| c | O-acetylhomoserine aminocarboxypropyltransferase | HVO_2997 | C | 1.3 | hvo00270 - Cysteine and methionine metabolism; | |
| c | 4-hydroxythreonine-4-phosphate dehydrogenase | HVO_2111 | C | 1.3 | hvo00750 - Vitamin B6 metabolism | |
| c | NADH dehydrogenase/oxidoreductase-like protein | HVO_2205 | C | 1.3 | hvo00190 - Oxidative phosphorylation; | |
| c | molybdopterin biosynthesis protein moeA | HVO_2304 | C | 1.3 | | COG0303 Code H Molybdopterin biosynthesis enzyme |
| c | NADH-quinone oxidoreductase chain c/d | HVO_0968 | C | 1.3 | hvo00190 - Oxidative phosphorylation; | |
| c | chlorite dismutase family protein | HVO_1871 | Unknown | 1.2 | | COG3253 Code S Uncharacterized conserved protein |
| c | cetyl-CoA synthetase | HVO_1000 | C | 1.2 | | COG1042 Code C Acyl-CoA synthetase (NDP forming) |
| c | SAM-dependent methyltransferase | HVO_1389 | Unknown | 1.2 | | COG2226 Code H Methylase involved in ubiquinone/menaquinone biosynthesis |
| c | ba3-type terminal oxidase subunit CbaD | HVO_0943 | Unknown | 1.2 | | |

| | | | | | | |
|---|---|---|---|---|---|---|
| c | molybdopterin biosynthesis protein moeA | HVO_2305 | C | 1.2 | | COG0303 Code H Molybdopterin biosynthesis enzyme |
| c M S | hypothetical | HVO_1431 | Extra-cellular | 1.2 | | |
| m | gluconate permease GntP | HVO_2190 | C/M | 1.2 | | COG2610 Code GE H+/gluconate symporter and related permeases |
| c | hypothetical protein | HVO_2770 | Unknown | 1.1 | | |
| m | ABC-type transport system permease protein (probable substrate dipeptides/oligopeptides) | HVO_B0081 | C/M | 1.1 | hvo02010 - ABC transporters | |
| c | HydD | HVO_1568 | Unknown | 1.1 | | COG0596 Code R Predicted hydrolases or acyltransferases (alpha/beta hydrolase superfamily) |
| m | twin arginine translocation system subunit TatAt | HVO_1162 | Unknown | 1.1 | hvo03060 - Protein export;<br>hvo03070 - Bacterial secretion system | |
| c | UspA domain-containing protein | HVO_2156 | C | 1.1 | | COG0589 Code T Universal stress protein UspA and related nucleotide-binding proteins |
| c | CBS domain pair | HVO_2384 | C | 1.1 | | COG3448 Code T CBS-domain-containing m protein; homolog=chloride channel |
| c | hypothetical protein | HVO_1749 | Unknown | 1.1 | | |
| c | threonine ammonia-lyase | HVO_0464 | C | 1.0 | hvo00260 - Glycine, serine and threonine metabolism; hvo00290 - Valine, leucine and isoleucine biosynthesis; hvo01110 - Biosynthesis of secondary metabolites | |
| m | putative phosphate ABC transporter ATP-binding protein | HVO_2378 | C/M | 1.0 | hvo02010 - ABC transporters | |
| m | ABC-type transport system permease protein (probable substrate dipeptides/oligopeptides) | HVO_B0080 | C/M | 1.0 | hvo02010 - ABC transporters | |
| c | ATP-binding protein Mrp | HVO_2790 | C | 1.0 | | COG0489 Code D ATPases involved in chromosome partitioning |
| c | sugar nucleotidyltransferase | HVO_1076 | C | 1.0 | hvo00521 - Streptomycin biosynthesis; hvo00523 - Polyketide sugar unit biosynthesis; hvo01110 - Biosynthesis of secondary metabolites | |
| c M S | hypothetical | HVO_0972 | Extra-cellular | 1.0 | | |
| c | hypothetical protein | HVO_1660 | C | 0.9 | | |
| c | dihydrolipoyl dehydrogenase | HVO_2961 | C | 0.9 | hvo00010 - Glycolysis / Gluconeogenesis; hvo00020 - Citrate cycle (TCA cycle); hvo00260 - Glycine, serine and threonine metabolism; hvo00280 - Valine, leucine and isoleucine degradation; hvo00620 - Pyruvate metabolism; hvo01110 - Biosynthesis of secondary metabolites | |
| c | glutamate dehydrogenase | HVO_1453 | C | 0.9 | hvo00250 - Alanine, aspartate and glutamate metabolism; hvo00330 - Arginine and proline metabolism; hvo00910 - Nitrogen metabolism; | |
| c | ATP phosphoribosyltransferase | HVO_0161 | C | 0.9 | hvo00340 - Histidine metabolism; hvo01110 - Biosynthesis of secondary metabolites | |
| c | peptidyl-prolyl cis-trans isomerase | HVO_1637 | C | 0.9 | | COG1047 Code O FKBP-type peptidyl-prolyl cis-trans isomerases 2 |
| c | indole-3-acetyl-L-aspartic acid hydrolase | HVO_1395 | C | 0.9 | | COG1473 Code R Metal-dependent amidase/aminoacylase/carboxypeptidase |

| | | | | | | |
|---|---|---|---|---|---|---|
| c M S | aminopeptidase | HVO_0836 | Extra-cellular | 0.9 | | OG2234 Code R Predicted aminopeptidases |
| m | hypothetical | HVO_0958 | C/M | 0.9 | | |
| c | TATA-binding transcription initiation factor | HVO_1727 | C | 0.9 | hvo03022 - Basal transcription factors | |
| m | ABC transporter permease | HVO_2084 | C/M | 0.9 | | COG4591 Code M ABC-type transport system, involved in lipoprotein release, permease component |
| c | hypothetical protein | HVO_A0181 | Unknown | 0.9 | | |
| m | hypothetical | HVO_0702 | C/M | 0.9 | | |
| c | cysteine desulfurase | HVO_0109 | C | 0.9 | hvo00450 - Selenoamino acid metabolism; hvo00730 - Thiamine metabolism | |
| c | phenylalanyl-tRNA synthetase subunit alpha | HVO_2948 | C | 0.9 | hvo00970 - Aminoacyl-tRNA biosynthesis | |
| c M S | ABC-type transport system periplasmic substrate-binding protein (probable substrate dipeptides/oligopeptides) | HVO_A0380 | Cellwall | 0.9 | hvo02010 - ABC transporters | |
| m | Na+/H+ antiporter NhaC | HVO_1394 | C/M | 0.8 | | OG1757 Code C Na+/H+ antiporter |
| m | sugar transporter | HVO_2097 | C/M | 0.8 | | COG2223 Code P Nitrate/nitrite transporter |
| c | hypothetical protein | HVO_1803 | Unknown | 0.8 | | |
| c | imidazoleglycerol phosphate synthase, cyclase subunit | HVO_1044 | C | 0.8 | hvo00340 - Histidine metabolism; hvo01110 - Biosynthesis of secondary metabolites | |
| c | putative yjeF family carbohydrate kinase | HVO_1348 | C | 0.8 | | COG0062 Code S Uncharacterized conserved protein |
| c | thioredoxin-disulfide reductase | HVO_1123 | C | 0.8 | hvo00240 - Pyrimidine metabolism; hvo00450 - Selenoamino acid metabolism | |
| c | 2-oxo-3-methylvalerate dehydrogenase E1 component subunit alpha | HVO_2958 | C | 0.8 | hvo00010 - Glycolysis / Gluconeogenesis; hvo00020 - Citrate cycle (TCA cycle); hvo00290 - Valine, leucine and isoleucine biosynthesis; hvo00620 - Pyruvate metabolism; hvo00650 - Butanoate metabolism; hvo01110 - Biosynthesis of secondary metabolites | |
| c | phosphoenolpyruvate-protein phosphotransferase | HVO_1496 | C | 0.8 | hvo02060 - Phosphotransferase system (PTS) | |
| m | putative glutamine ABC transporter permease | HVO_2431 | C/M | 0.8 | | COG0765 Code E ABC-type amino acid transport system, permease component |
| c | pyruvate--ferredoxin oxidoreductase subunit alpha | HVO_1305 | C | 0.8 | hvo00020 - Citrate cycle (TCA cycle); hvo01120 - Microbial metabolism in diverse environments | |
| c | metal-dependent carboxypeptidase | HVO_0417 | C | 0.8 | | COG2317 Code E Zn-dependent carboxypeptidase |

**Supplementary Table 3. Strains**

| **Strain** | **Genotype** | **Source** |
|---|---|---|
| DH5α | $F^-$, φ80d*lacZ*ΔM15, Δ(*lacZYA-argF*)U169, *deoR*, *recA*1, *endA*1, *hsdR*17($r_k^-$, $m_k^+$), *phoA*, *supE*44,$\lambda^-$, *thi*-1, *gyrA*96, *relA*1 | Invitrogen |
| H119 | DS70 (ΔpHV2), Δ*pyrE*2, Δ*trpA*, Δ*leuB* | Allers et al., 2004 |
| H66 | DS70 (ΔpHV2), Δ*pyrE*2, Δ*leuB* | Allers et al., 2004 |
| Δ*s479* | DS70 (ΔpHV2), Δ*pyrE2*, Δ*trpA*, Δ*leuB*, Δ*s479::trpA*$^+$ | Jaschinski et al., 2014 |
| Δ*cas6* | Δ*pyr*E2, Δ*trp*A, Δ*leu*B, Δ*cas6* | Brendel et al., 2014 |
| Δ*cas7* | Δ*pyr*E2, Δ*trp*A, Δ*leu*B, Δ*cas*7 | Brendel et al., 2014 |

**Supplementary Table 4. Oligonucleotides and Sequencing Barcodes**

| Primer | Sequence | Used for |
|---|---|---|
| TruSeq_Sense_primer | 5'-AATGATACGGCGACCACCGAGATCTACAC-NNNNNNNN-ACACTCTTTCCCTACACGACGCTCTTCCGATCT-3'; NNN=i5 Barcode | RNA-Seq |
| TruSeq_Antisense_primer | 5'-CAAGCAGAAGACGGCATACGAGAT-NNNNNNNN-GTGACTGGAGTTCAGACGTGTGCTCTTCCGATCT-3'; NNN= i7 Index | RNA-Seq |
| Delta-s479-Nr1 | i5 Barcode AGGCTATA | RNA-Seq |
| | i7 Index ATTACTCG | |
| Delta-s479-Nr2 | i5 Barcode GCCTCTAT | RNA-Seq |
| | i7 Index ATTACTCG | |
| Delta-s479-Nr3 | i5 Barcode GTCAGTAC | RNA-Seq |
| | i7 Index ATTACTCG | |
| H66-Nr1 | i5 Barcode AGGATAGG | RNA-Seq |
| | i7 Index ATTACTCG | |
| H66-Nr2 | i5 Barcode TCAGAGCC | RNA-Seq |
| | i7 Index ATTACTCG | |
| H66-Nr3 | i5 Barcode ATAGAGAG | RNA-Seq |
| | i7 Index ATTACTCG | |
| probe znuC1 Hvo_2398fw | CCGCGAGGTCGTGAAGATGGGTCGG | northern blot |
| probe znuC1 Hvo_2398rev | AGGTCGTGTTCGATGAGGAGAATCG | northern blot |
| 16Sseqf | CGCTAGGTGTGACACAGGCTACG | northern blot |
| 16Sseqrev | GTGAGATGTCCGGCGTTGAGTCC | northern blot |
| Hvo5Sp | CGCAGGTGAGCTTAACTTCCGTGTTCGGG | northern blot |
| s479spacerpart | CGCTTCGGCTGATGGGTGTCATGAGGTTTCCCTTGTCAATACTT | northern blot |
| s479 RNA | AAGUAUUGACAAGGGAAACCUCAUGACACCCAUCAGCCGAAGCGACAAACUGGGU | EMSA |
| Znucmid RNA | GGUCGUGAAGAUGGGUCGGUUCCCCCACGUCGGCUUCGGCCGGCUCUCGGCCGAAGACCACCGCAUCGUCGACGAGGCGC | EMSA |
| Znucmid mutant RNA | GGUCGUGAAGAUGGGUCGGUUCCCCCAC-CGGCUCUCGGCCGAAGACCACCGCAUCGUCGACGAGGCGC | EMSA |

**Supplementary Table 5. Data and calculations for growth experiments at standard zinc concentration.** Supplementary Table 5A1. and A2. list the data used to compile the growth curves in Figure 4. Supplementary Table 5B. shows the calculations of growth rate and doubling time for wildtype and deletion strain (Δ*s479*). Supplementary Table 5C1. and C2. list all the raw data collected for the growth experiments. For better readability, all numbers were cut to three decimals or less. **Supplementary Table 5A1. Mean growth and standard deviation used to compile the wildtype strain (H66) growth curve in Figure 4.** The mean $OD_{650nm}$ is given for all three biological replicates of wildtype strain H66 (H66-1 to H66-3) alongside the overall mean $OD_{650nm}$ and standard deviation per time interval.

| **H66-1** | **H66-2** | **H66-3** | **H66** | **H66** | |
|---|---|---|---|---|---|
| **Mean OD650** | **Mean OD650** | **Mean OD650** | **Mean OD650** | **SD** | **time [h]** |
| 0.066 | 0.068 | 0.065 | **0.066** | 0.001 | **0.5** |
| 0.100 | 0.099 | 0.094 | **0.098** | 0.003 | **2.5** |
| 0.147 | 0.143 | 0.130 | **0.140** | 0.007 | **4.5** |
| 0.201 | 0.196 | 0.175 | **0.190** | 0.011 | **6.5** |
| 0.246 | 0.248 | 0.229 | **0.241** | 0.009 | **8.5** |
| 0.289 | 0.297 | 0.287 | **0.291** | 0.004 | **10.5** |
| 0.336 | 0.342 | 0.344 | **0.340** | 0.003 | **12.5** |
| 0.372 | 0.385 | 0.391 | **0.383** | 0.008 | **14.5** |
| 0.405 | 0.431 | 0.430 | **0.422** | 0.012 | **16.5** |
| 0.441 | 0.477 | 0.462 | **0.460** | 0.015 | **18.5** |
| 0.480 | 0.516 | 0.490 | **0.496** | 0.015 | **20.5** |
| 0.520 | 0.549 | 0.514 | **0.528** | 0.015 | **22.5** |
| 0.551 | 0.575 | 0.534 | **0.554** | 0.017 | **24.5** |
| 0.574 | 0.595 | 0.552 | **0.574** | 0.017 | **26.5** |
| 0.592 | 0.611 | 0.567 | **0.590** | 0.018 | **28.5** |
| 0.607 | 0.622 | 0.579 | **0.603** | 0.018 | **30.5** |
| 0.618 | 0.632 | 0.591 | **0.614** | 0.017 | **32.5** |
| 0.630 | 0.642 | 0.601 | **0.624** | 0.017 | **34.5** |
| 0.640 | 0.650 | 0.610 | **0.633** | 0.017 | **36.5** |
| 0.649 | 0.659 | 0.619 | **0.642** | 0.017 | **38.5** |
| 0.657 | 0.665 | 0.628 | **0.650** | 0.016 | **40.5** |
| 0.663 | 0.671 | 0.636 | **0.656** | 0.015 | **42.5** |
| 0.668 | 0.676 | 0.643 | **0.662** | 0.014 | **44.5** |
| 0.668 | 0.678 | 0.649 | **0.665** | 0.012 | **46.5** |
| 0.666 | 0.681 | 0.655 | **0.667** | 0.011 | **48.5** |
| 0.669 | 0.681 | 0.660 | **0.670** | 0.009 | **50.5** |
| 0.672 | 0.684 | 0.664 | **0.673** | 0.008 | **52.5** |
| 0.675 | 0.688 | 0.667 | **0.677** | 0.008 | **54.5** |
| 0.677 | 0.689 | 0.671 | **0.679** | 0.008 | **56.5** |
| 0.680 | 0.691 | 0.673 | **0.681** | 0.007 | **58.5** |
| 0.684 | 0.696 | 0.677 | **0.685** | 0.008 | **60.5** |
| 0.686 | 0.698 | 0.679 | **0.688** | 0.008 | **62.5** |
| 0.689 | 0.700 | 0.683 | **0.691** | 0.007 | **64.5** |
| 0.692 | 0.703 | 0.686 | **0.694** | 0.007 | **66.5** |
| 0.696 | 0.705 | 0.690 | **0.697** | 0.006 | **68.5** |
| 0.700 | 0.706 | 0.693 | **0.700** | 0.005 | **70.5** |
| 0.697 | 0.707 | 0.695 | **0.700** | 0.005 | **72.5** |

**Supplementary Table 5A2. Mean growth and standard deviation used to compile the deletion strain (Δ*s479*) growth curve in Figure 4.** The mean $OD_{650nm}$ is given for all three biological replicates of deletion strain Δ*s479* (s479-1 to s479-3) alongside the overall mean $OD_{650nm}$ and standard deviation per time interval.

| s479-1 | s479-2 | s479-3 | s479 | s479 SD | |
|---|---|---|---|---|---|
| **Mean OD650** | **Mean OD650** | **Mean OD650** | **Mean OD650** | | **time [h]** |
| 0.065 | 0.055 | 0.064 | **0.061** | 0.004 | **0.5** |
| 0.101 | 0.086 | 0.100 | **0.096** | 0.007 | **2.5** |
| 0.142 | 0.120 | 0.138 | **0.133** | 0.010 | **4.5** |
| 0.192 | 0.162 | 0.187 | **0.180** | 0.013 | **6.5** |
| 0.251 | 0.210 | 0.243 | **0.235** | 0.018 | **8.5** |
| 0.306 | 0.260 | 0.295 | **0.287** | 0.020 | **10.5** |
| 0.352 | 0.305 | 0.337 | **0.331** | 0.020 | **12.5** |
| 0.387 | 0.341 | 0.371 | **0.366** | 0.019 | **14.5** |
| 0.413 | 0.373 | 0.401 | **0.396** | 0.017 | **16.5** |
| 0.437 | 0.401 | 0.429 | **0.422** | 0.015 | **18.5** |
| 0.457 | 0.426 | 0.457 | **0.447** | 0.015 | **20.5** |
| 0.478 | 0.448 | 0.481 | **0.469** | 0.015 | **22.5** |
| 0.495 | 0.468 | 0.501 | **0.488** | 0.014 | **24.5** |
| 0.510 | 0.484 | 0.519 | **0.505** | 0.015 | **26.5** |
| 0.522 | 0.498 | 0.535 | **0.518** | 0.015 | **28.5** |
| 0.534 | 0.511 | 0.549 | **0.531** | 0.015 | **30.5** |
| 0.543 | 0.523 | 0.561 | **0.542** | 0.015 | **32.5** |
| 0.553 | 0.534 | 0.572 | **0.553** | 0.015 | **34.5** |
| 0.561 | 0.543 | 0.583 | **0.562** | 0.016 | **36.5** |
| 0.568 | 0.553 | 0.592 | **0.571** | 0.016 | **38.5** |
| 0.577 | 0.561 | 0.602 | **0.580** | 0.017 | **40.5** |
| 0.584 | 0.568 | 0.610 | **0.587** | 0.017 | **42.5** |
| 0.590 | 0.576 | 0.624 | **0.597** | 0.020 | **44.5** |
| 0.597 | 0.582 | 0.635 | **0.605** | 0.022 | **46.5** |
| 0.602 | 0.590 | 0.639 | **0.610** | 0.021 | **48.5** |
| 0.608 | 0.597 | 0.640 | **0.615** | 0.018 | **50.5** |
| 0.612 | 0.601 | 0.638 | **0.617** | 0.015 | **52.5** |
| 0.615 | 0.604 | 0.642 | **0.620** | 0.016 | **54.5** |
| 0.617 | 0.607 | 0.648 | **0.624** | 0.017 | **56.5** |
| 0.619 | 0.610 | 0.654 | **0.628** | 0.019 | **58.5** |
| 0.619 | 0.613 | 0.659 | **0.630** | 0.020 | **60.5** |
| 0.621 | 0.614 | 0.666 | **0.634** | 0.023 | **62.5** |
| 0.623 | 0.615 | 0.670 | **0.636** | 0.024 | **64.5** |
| 0.623 | 0.617 | 0.672 | **0.637** | 0.024 | **66.5** |
| 0.625 | 0.619 | 0.670 | **0.638** | 0.023 | **68.5** |
| 0.626 | 0.620 | 0.668 | **0.638** | 0.021 | **70.5** |
| 0.626 | 0.621 | 0.667 | **0.638** | 0.021 | **72.5** |

**Supplementary Table 5B. Calculation of growth rate and doubling time at standard zinc concentration.** Growth rate and doubling time were calculated as growth rate μ = (ln(xt) – ln (x0)) / (t – t0) and doubling time d= ln(2) / μ. Calculations were carried out separately for all replicates before calculating mean value (red) and standard deviation (red). Phases of exponential growth were identified using fitted trendlines and corresponding $R^2$-values (Supplementary Figure 5C.). Values are given separately for H66 (grey) and Δ*s479* (orange) and for each of the two phases.

**phase1 0.5 to 10.5 h**

| growth rate μ(H66) [$h^{-1}$] time interval 0.5 h to 10.5 h with R² = 0.983 | | | | |
|---|---|---|---|---|
| **H66-1** | **H66-2** | **H66-3** | **μ(H66) [$h^{-1}$]** | **SD(μ(H66)) [$h^{-1}$]** |
| 0.147 | 0.148 | 0.149 | **0.148** | **0.001** |

| doubling time / generation time d(H66) [h] | | | | |
|---|---|---|---|---|
| **H66-1** | **H66-2** | **H66-3** | **d(H66) [h]** | **SD(d(H66)) [h]** |
| 4.7 | 4.7 | 4.7 | **4.7** | **0.0** |

| growth rate μ(s479) [$h^{-1}$] time interval 0.5 h to 10.5 h with R² = 0.9836 | | | | |
|---|---|---|---|---|
| **s479-1** | **s479-2** | **s479-3** | **μ(s479) [$h^{-1}$]** | **SD(μ(s479)) [$h^{-1}$]** |
| 0.155 | 0.155 | 0.153 | **0.154** | **0.001** |

| doubling time / generation time d(H66) [h] | | | | |
|---|---|---|---|---|
| **s479-1** | **s479-2** | **s479-3** | **d(s479) [h]** | **SD(d(s479)) [h]** |
| 4.5 | 4.5 | 4.5 | **4.5** | **0.0** |

**phase2 14.5 to 24.5 h**

| growth rate μ(H66) [$h^{-1}$] time interval 14.5 h to 24.5 h with R² = 0.9857 | | | | |
|---|---|---|---|---|
| **H66-1** | **H66-2** | **H66-3** | **μ(H66) [$h^{-1}$]** | **SD(μ(H66)) [$h^{-1}$]** |
| 0.039 | 0.040 | 0.031 | **0.037** | **0.004** |

| doubling time / generation time d(H66) [h] | | | | |
|---|---|---|---|---|
| **H66-1** | **H66-2** | **H66-3** | **d(H66) [h]** | **SD(d(H66)) [h]** |
| 17.6 | 17.3 | 22.1 | **19.0** | **2.2** |

| growth rate μ(s479) [$h^{-1}$] time interval 14.5 h to 24.5 h with R² = 0.9866 | | | | |
|---|---|---|---|---|
| **s479-1** | **s479-2** | **s479-3** | **μ(s479) [$h^{-1}$]** | **SD(μ(s479)) [$h^{-1}$]** |
| 0.025 | 0.032 | 0.030 | **0.029** | **0.003** |

| doubling time / generation time d(H66) [h] | | | | |
|---|---|---|---|---|
| **s479-1** | **s479-2** | **s479-3** | **d(s479) [h]** | **SD(d(s479)) [h]** |
| 28.1 | 21.8 | 23.1 | **24.4** | **2.7** |

**Supplementary Table 5C1. Raw growth data measured for the wildtype strain (H66) at standard zinc concentration.** For each time point, $OD_{650nm}$ of three biological (replicate 1-3; H66-1 to H66-3), with three technical replicates (A-C) each, are given. Additionally, the mean $OD_{650nm}$ over all technical replicates is given for each biological replicate.

| | replicate 1 | | | | replicate 2 | | | | replicate 3 | | | |
|---|---|---|---|---|---|---|---|---|---|---|---|---|
| | **H66-1A** | **H66-1B** | **H66-1C** | **H66-1** | **H66-2A** | **H66-2B** | **H66-2C** | **H66-2** | **H66-3A** | **H66-3B** | **H66-3C** | **H66-3** |
| **time [h]** | OD650 | OD650 | OD650 | **Mean OD650** | OD650 | OD650 | OD650 | **Mean OD650** | OD650 | OD650 | OD650 | **Mean OD650** |
| **0.5** | 0.068 | 0.061 | 0.07 | 0.066 | 0.065 | 0.066 | 0.073 | 0.068 | 0.069 | 0.066 | 0.059 | 0.065 |
| **1** | 0.074 | 0.065 | 0.075 | 0.071 | 0.068 | 0.071 | 0.079 | 0.073 | 0.073 | 0.071 | 0.063 | 0.069 |
| **1.5** | 0.083 | 0.074 | 0.084 | 0.080 | 0.079 | 0.079 | 0.087 | 0.082 | 0.082 | 0.08 | 0.071 | 0.078 |
| **2** | 0.094 | 0.084 | 0.093 | 0.090 | 0.088 | 0.089 | 0.097 | 0.091 | 0.092 | 0.089 | 0.079 | 0.087 |
| **2.5** | 0.101 | 0.095 | 0.103 | 0.100 | 0.097 | 0.097 | 0.104 | 0.099 | 0.099 | 0.097 | 0.086 | 0.094 |
| **3** | 0.112 | 0.106 | 0.111 | 0.110 | 0.106 | 0.107 | 0.113 | 0.109 | 0.108 | 0.106 | 0.094 | 0.103 |
| **3.5** | 0.124 | 0.119 | 0.121 | 0.121 | 0.117 | 0.119 | 0.122 | 0.119 | 0.117 | 0.115 | 0.101 | 0.111 |
| **4** | 0.137 | 0.132 | 0.133 | 0.134 | 0.129 | 0.131 | 0.132 | 0.131 | 0.127 | 0.125 | 0.109 | 0.120 |
| **4.5** | 0.15 | 0.146 | 0.146 | 0.147 | 0.142 | 0.144 | 0.144 | 0.143 | 0.135 | 0.136 | 0.119 | 0.130 |
| **5** | 0.163 | 0.159 | 0.16 | 0.161 | 0.155 | 0.158 | 0.154 | 0.156 | 0.146 | 0.147 | 0.128 | 0.140 |
| **5.5** | 0.178 | 0.174 | 0.174 | 0.175 | 0.17 | 0.171 | 0.165 | 0.169 | 0.156 | 0.159 | 0.138 | 0.151 |
| **6** | 0.191 | 0.186 | 0.186 | 0.188 | 0.184 | 0.185 | 0.178 | 0.182 | 0.168 | 0.171 | 0.148 | 0.162 |
| **6.5** | 0.204 | 0.198 | 0.201 | 0.201 | 0.199 | 0.197 | 0.191 | 0.196 | 0.18 | 0.184 | 0.16 | 0.175 |
| **7** | 0.217 | 0.207 | 0.214 | 0.213 | 0.214 | 0.21 | 0.205 | 0.210 | 0.192 | 0.197 | 0.17 | 0.186 |
| **7.5** | 0.23 | 0.215 | 0.228 | 0.224 | 0.228 | 0.221 | 0.219 | 0.223 | 0.205 | 0.211 | 0.183 | 0.200 |
| **8** | 0.242 | 0.223 | 0.244 | 0.236 | 0.241 | 0.231 | 0.233 | 0.235 | 0.219 | 0.228 | 0.195 | 0.214 |
| **8.5** | 0.255 | 0.229 | 0.254 | 0.246 | 0.255 | 0.241 | 0.249 | 0.248 | 0.234 | 0.243 | 0.209 | 0.229 |
| **9** | 0.267 | 0.234 | 0.271 | 0.257 | 0.268 | 0.252 | 0.264 | 0.261 | 0.247 | 0.26 | 0.224 | 0.244 |
| **9.5** | 0.28 | 0.239 | 0.285 | 0.268 | 0.28 | 0.261 | 0.282 | 0.274 | 0.26 | 0.276 | 0.237 | 0.258 |
| **10** | 0.294 | 0.243 | 0.298 | 0.278 | 0.294 | 0.268 | 0.294 | 0.285 | 0.275 | 0.292 | 0.251 | 0.273 |
| **10.5** | 0.309 | 0.247 | 0.312 | 0.289 | 0.305 | 0.277 | 0.31 | 0.297 | 0.288 | 0.308 | 0.265 | 0.287 |
| **11** | 0.325 | 0.253 | 0.326 | 0.301 | 0.314 | 0.284 | 0.323 | 0.307 | 0.3 | 0.323 | 0.28 | 0.301 |
| **11.5** | 0.344 | 0.257 | 0.341 | 0.314 | 0.324 | 0.293 | 0.336 | 0.318 | 0.313 | 0.337 | 0.294 | 0.315 |
| **12** | 0.363 | 0.261 | 0.352 | 0.325 | 0.338 | 0.303 | 0.351 | 0.331 | 0.328 | 0.354 | 0.307 | 0.330 |
| **12.5** | 0.379 | 0.263 | 0.366 | 0.336 | 0.35 | 0.311 | 0.364 | 0.342 | 0.342 | 0.368 | 0.321 | 0.344 |
| **13** | 0.392 | 0.268 | 0.376 | 0.345 | 0.361 | 0.321 | 0.376 | 0.353 | 0.353 | 0.381 | 0.333 | 0.356 |
| **13.5** | 0.404 | 0.27 | 0.389 | 0.354 | 0.371 | 0.331 | 0.388 | 0.363 | 0.366 | 0.393 | 0.348 | 0.369 |
| **14** | 0.415 | 0.275 | 0.397 | 0.362 | 0.383 | 0.341 | 0.398 | 0.374 | 0.376 | 0.403 | 0.36 | 0.380 |
| **14.5** | 0.426 | 0.281 | 0.408 | 0.372 | 0.394 | 0.352 | 0.41 | 0.385 | 0.387 | 0.414 | 0.371 | 0.391 |
| **15** | 0.436 | 0.287 | 0.416 | 0.380 | 0.407 | 0.366 | 0.419 | 0.397 | 0.396 | 0.425 | 0.381 | 0.401 |
| **15.5** | 0.442 | 0.294 | 0.427 | 0.388 | 0.419 | 0.379 | 0.43 | 0.409 | 0.408 | 0.433 | 0.391 | 0.411 |
| **16** | 0.451 | 0.302 | 0.434 | 0.396 | 0.43 | 0.39 | 0.437 | 0.419 | 0.418 | 0.441 | 0.402 | 0.420 |
| **16.5** | 0.458 | 0.312 | 0.445 | 0.405 | 0.441 | 0.404 | 0.449 | 0.431 | 0.427 | 0.45 | 0.412 | 0.430 |
| **17** | 0.466 | 0.323 | 0.452 | 0.414 | 0.451 | 0.418 | 0.458 | 0.442 | 0.438 | 0.456 | 0.422 | 0.439 |
| **17.5** | 0.47 | 0.337 | 0.461 | 0.423 | 0.464 | 0.432 | 0.465 | 0.454 | 0.446 | 0.464 | 0.431 | 0.447 |
| **18** | 0.476 | 0.352 | 0.469 | 0.432 | 0.475 | 0.448 | 0.472 | 0.465 | 0.455 | 0.473 | 0.439 | 0.456 |
| **18.5** | 0.479 | 0.366 | 0.477 | 0.441 | 0.486 | 0.464 | 0.48 | 0.477 | 0.461 | 0.479 | 0.446 | 0.462 |
| **19** | 0.484 | 0.38 | 0.485 | 0.450 | 0.496 | 0.477 | 0.488 | 0.487 | 0.469 | 0.486 | 0.455 | 0.470 |
| **19.5** | 0.492 | 0.397 | 0.493 | 0.461 | 0.506 | 0.492 | 0.496 | 0.498 | 0.476 | 0.493 | 0.462 | 0.477 |
| **20** | 0.497 | 0.411 | 0.499 | 0.469 | 0.515 | 0.502 | 0.501 | 0.506 | 0.482 | 0.5 | 0.469 | 0.484 |

| | | | | | | | | | | | | | | |
|---|---|---|---|---|---|---|---|---|---|---|---|---|---|---|
| **20.5** | 0.503 | 0.431 | 0.507 | 0.480 | | 0.524 | 0.516 | 0.508 | 0.516 | | 0.489 | 0.506 | 0.476 | 0.490 |
| **21** | 0.507 | 0.453 | 0.513 | 0.491 | | 0.531 | 0.526 | 0.513 | 0.523 | | 0.497 | 0.511 | 0.481 | 0.496 |
| **21.5** | 0.512 | 0.472 | 0.52 | 0.501 | | 0.541 | 0.538 | 0.521 | 0.533 | | 0.502 | 0.517 | 0.488 | 0.502 |
| **22** | 0.517 | 0.493 | 0.527 | 0.512 | | 0.549 | 0.55 | 0.526 | 0.542 | | 0.508 | 0.523 | 0.494 | 0.508 |
| **22.5** | 0.521 | 0.508 | 0.532 | 0.520 | | 0.557 | 0.559 | 0.532 | 0.549 | | 0.515 | 0.528 | 0.5 | 0.514 |
| **23** | 0.525 | 0.523 | 0.538 | 0.529 | | 0.564 | 0.568 | 0.537 | 0.556 | | 0.519 | 0.534 | 0.504 | 0.519 |
| **23.5** | 0.532 | 0.532 | 0.544 | 0.536 | | 0.569 | 0.574 | 0.544 | 0.562 | | 0.525 | 0.538 | 0.512 | 0.525 |
| **24** | 0.538 | 0.544 | 0.55 | 0.544 | | 0.576 | 0.583 | 0.548 | 0.569 | | 0.529 | 0.543 | 0.516 | 0.529 |
| **24.5** | 0.543 | 0.556 | 0.555 | 0.551 | | 0.581 | 0.591 | 0.554 | 0.575 | | 0.534 | 0.548 | 0.521 | 0.534 |
| **25** | 0.548 | 0.566 | 0.56 | 0.558 | | 0.586 | 0.597 | 0.557 | 0.580 | | 0.539 | 0.553 | 0.526 | 0.539 |
| **25.5** | 0.552 | 0.572 | 0.565 | 0.563 | | 0.591 | 0.603 | 0.563 | 0.586 | | 0.544 | 0.558 | 0.531 | 0.544 |
| **26** | 0.558 | 0.577 | 0.569 | 0.568 | | 0.596 | 0.607 | 0.567 | 0.590 | | 0.547 | 0.561 | 0.535 | 0.548 |
| **26.5** | 0.563 | 0.585 | 0.574 | 0.574 | | 0.599 | 0.613 | 0.572 | 0.595 | | 0.552 | 0.565 | 0.539 | 0.552 |
| **27** | 0.568 | 0.59 | 0.579 | 0.579 | | 0.604 | 0.619 | 0.576 | 0.600 | | 0.556 | 0.569 | 0.543 | 0.556 |
| **27.5** | 0.572 | 0.596 | 0.583 | 0.584 | | 0.608 | 0.623 | 0.577 | 0.603 | | 0.56 | 0.573 | 0.547 | 0.560 |
| **28** | 0.576 | 0.6 | 0.586 | 0.587 | | 0.611 | 0.625 | 0.583 | 0.606 | | 0.563 | 0.575 | 0.552 | 0.563 |
| **28.5** | 0.58 | 0.605 | 0.591 | 0.592 | | 0.615 | 0.632 | 0.586 | 0.611 | | 0.567 | 0.579 | 0.555 | 0.567 |
| **29** | 0.584 | 0.609 | 0.595 | 0.596 | | 0.617 | 0.633 | 0.589 | 0.613 | | 0.57 | 0.581 | 0.559 | 0.570 |
| **29.5** | 0.587 | 0.613 | 0.599 | 0.600 | | 0.619 | 0.637 | 0.593 | 0.616 | | 0.573 | 0.584 | 0.561 | 0.573 |
| **30** | 0.592 | 0.618 | 0.602 | 0.604 | | 0.623 | 0.641 | 0.597 | 0.620 | | 0.576 | 0.586 | 0.566 | 0.576 |
| **30.5** | 0.595 | 0.62 | 0.606 | 0.607 | | 0.624 | 0.642 | 0.6 | 0.622 | | 0.578 | 0.589 | 0.569 | 0.579 |
| **31** | 0.599 | 0.623 | 0.609 | 0.610 | | 0.629 | 0.646 | 0.604 | 0.626 | | 0.581 | 0.592 | 0.573 | 0.582 |
| **31.5** | 0.601 | 0.625 | 0.613 | 0.613 | | 0.63 | 0.648 | 0.606 | 0.628 | | 0.584 | 0.596 | 0.576 | 0.585 |
| **32** | 0.606 | 0.627 | 0.616 | 0.616 | | 0.633 | 0.651 | 0.611 | 0.632 | | 0.587 | 0.599 | 0.58 | 0.589 |
| **32.5** | 0.608 | 0.627 | 0.62 | 0.618 | | 0.633 | 0.651 | 0.613 | 0.632 | | 0.588 | 0.601 | 0.583 | 0.591 |
| **33** | 0.612 | 0.628 | 0.622 | 0.621 | | 0.636 | 0.652 | 0.616 | 0.635 | | 0.59 | 0.603 | 0.585 | 0.593 |
| **33.5** | 0.614 | 0.631 | 0.626 | 0.624 | | 0.639 | 0.656 | 0.618 | 0.638 | | 0.592 | 0.606 | 0.587 | 0.595 |
| **34** | 0.617 | 0.634 | 0.629 | 0.627 | | 0.64 | 0.657 | 0.621 | 0.639 | | 0.595 | 0.607 | 0.59 | 0.597 |
| **34.5** | 0.621 | 0.638 | 0.631 | 0.630 | | 0.642 | 0.66 | 0.624 | 0.642 | | 0.598 | 0.611 | 0.593 | 0.601 |
| **35** | 0.623 | 0.639 | 0.634 | 0.632 | | 0.644 | 0.661 | 0.627 | 0.644 | | 0.601 | 0.614 | 0.595 | 0.603 |
| **35.5** | 0.624 | 0.642 | 0.638 | 0.635 | | 0.646 | 0.663 | 0.63 | 0.646 | | 0.603 | 0.615 | 0.597 | 0.605 |
| **36** | 0.626 | 0.646 | 0.64 | 0.637 | | 0.647 | 0.665 | 0.632 | 0.648 | | 0.605 | 0.617 | 0.599 | 0.607 |
| **36.5** | 0.628 | 0.649 | 0.643 | 0.640 | | 0.649 | 0.666 | 0.635 | 0.650 | | 0.608 | 0.619 | 0.603 | 0.610 |
| **37** | 0.631 | 0.652 | 0.646 | 0.643 | | 0.653 | 0.67 | 0.637 | 0.653 | | 0.61 | 0.622 | 0.604 | 0.612 |
| **37.5** | 0.632 | 0.654 | 0.649 | 0.645 | | 0.653 | 0.67 | 0.64 | 0.654 | | 0.612 | 0.624 | 0.608 | 0.615 |
| **38** | 0.634 | 0.655 | 0.651 | 0.647 | | 0.654 | 0.671 | 0.643 | 0.656 | | 0.614 | 0.626 | 0.61 | 0.617 |
| **38.5** | 0.636 | 0.658 | 0.653 | 0.649 | | 0.657 | 0.674 | 0.645 | 0.659 | | 0.617 | 0.627 | 0.612 | 0.619 |
| **39** | 0.637 | 0.661 | 0.656 | 0.651 | | 0.657 | 0.675 | 0.647 | 0.660 | | 0.619 | 0.63 | 0.615 | 0.621 |
| **39.5** | 0.639 | 0.663 | 0.658 | 0.653 | | 0.66 | 0.678 | 0.649 | 0.662 | | 0.623 | 0.632 | 0.618 | 0.624 |
| **40** | 0.641 | 0.666 | 0.661 | 0.656 | | 0.661 | 0.679 | 0.652 | 0.664 | | 0.624 | 0.634 | 0.619 | 0.626 |
| **40.5** | 0.642 | 0.666 | 0.663 | 0.657 | | 0.662 | 0.68 | 0.652 | 0.665 | | 0.626 | 0.635 | 0.622 | 0.628 |
| **41** | 0.643 | 0.667 | 0.664 | 0.658 | | 0.664 | 0.68 | 0.655 | 0.666 | | 0.629 | 0.638 | 0.625 | 0.631 |
| **41.5** | 0.644 | 0.668 | 0.666 | 0.659 | | 0.665 | 0.683 | 0.657 | 0.668 | | 0.632 | 0.639 | 0.626 | 0.632 |
| **42** | 0.647 | 0.67 | 0.669 | 0.662 | | 0.667 | 0.683 | 0.659 | 0.670 | | 0.633 | 0.641 | 0.627 | 0.634 |
| **42.5** | 0.647 | 0.67 | 0.671 | 0.663 | | 0.667 | 0.684 | 0.661 | 0.671 | | 0.636 | 0.642 | 0.63 | 0.636 |
| **43** | 0.648 | 0.671 | 0.671 | 0.663 | | 0.669 | 0.685 | 0.662 | 0.672 | | 0.637 | 0.643 | 0.63 | 0.637 |
| **43.5** | 0.65 | 0.672 | 0.674 | 0.665 | | 0.67 | 0.685 | 0.664 | 0.673 | | 0.639 | 0.645 | 0.634 | 0.639 |
| **44** | 0.651 | 0.672 | 0.676 | 0.666 | | 0.671 | 0.686 | 0.668 | 0.675 | | 0.641 | 0.647 | 0.634 | 0.641 |
| **44.5** | 0.653 | 0.672 | 0.678 | 0.668 | | 0.673 | 0.687 | 0.668 | 0.676 | | 0.643 | 0.649 | 0.637 | 0.643 |
| **45** | 0.654 | 0.67 | 0.68 | 0.668 | | 0.673 | 0.687 | 0.67 | 0.677 | | 0.645 | 0.65 | 0.638 | 0.644 |
| **45.5** | 0.655 | 0.669 | 0.68 | 0.668 | | 0.672 | 0.686 | 0.671 | 0.676 | | 0.647 | 0.652 | 0.64 | 0.646 |

| | | | | | | | | | | | | | | |
|---|---|---|---|---|---|---|---|---|---|---|---|---|---|---|
| **46** | 0.657 | 0.668 | 0.681 | 0.669 | | 0.674 | 0.687 | 0.672 | 0.678 | | 0.648 | 0.654 | 0.641 | 0.648 |
| **46.5** | 0.657 | 0.666 | 0.682 | 0.668 | | 0.675 | 0.685 | 0.673 | 0.678 | | 0.65 | 0.656 | 0.642 | 0.649 |
| **47** | 0.657 | 0.662 | 0.683 | 0.667 | | 0.676 | 0.686 | 0.674 | 0.679 | | 0.652 | 0.657 | 0.644 | 0.651 |
| **47.5** | 0.659 | 0.659 | 0.685 | 0.668 | | 0.676 | 0.687 | 0.676 | 0.680 | | 0.654 | 0.658 | 0.646 | 0.653 |
| **48** | 0.66 | 0.653 | 0.687 | 0.667 | | 0.676 | 0.686 | 0.676 | 0.679 | | 0.654 | 0.659 | 0.646 | 0.653 |
| **48.5** | 0.661 | 0.65 | 0.687 | 0.666 | | 0.677 | 0.687 | 0.679 | 0.681 | | 0.657 | 0.66 | 0.649 | 0.655 |
| **49** | 0.663 | 0.648 | 0.689 | 0.667 | | 0.678 | 0.686 | 0.678 | 0.681 | | 0.658 | 0.661 | 0.649 | 0.656 |
| **49.5** | 0.664 | 0.65 | 0.69 | 0.668 | | 0.679 | 0.687 | 0.68 | 0.682 | | 0.661 | 0.663 | 0.651 | 0.658 |
| **50** | 0.665 | 0.65 | 0.69 | 0.668 | | 0.679 | 0.686 | 0.681 | 0.682 | | 0.661 | 0.664 | 0.651 | 0.659 |
| **50.5** | 0.666 | 0.651 | 0.69 | 0.669 | | 0.679 | 0.685 | 0.68 | 0.681 | | 0.663 | 0.665 | 0.653 | 0.660 |
| **51** | 0.666 | 0.652 | 0.692 | 0.670 | | 0.679 | 0.685 | 0.684 | 0.683 | | 0.664 | 0.666 | 0.654 | 0.661 |
| **51.5** | 0.667 | 0.652 | 0.692 | 0.670 | | 0.68 | 0.686 | 0.681 | 0.682 | | 0.665 | 0.667 | 0.655 | 0.662 |
| **52** | 0.669 | 0.653 | 0.692 | 0.671 | | 0.682 | 0.685 | 0.684 | 0.684 | | 0.666 | 0.668 | 0.657 | 0.664 |
| **52.5** | 0.669 | 0.654 | 0.692 | 0.672 | | 0.682 | 0.686 | 0.684 | 0.684 | | 0.666 | 0.669 | 0.657 | 0.664 |
| **53** | 0.671 | 0.654 | 0.694 | 0.673 | | 0.683 | 0.687 | 0.684 | 0.685 | | 0.667 | 0.67 | 0.659 | 0.665 |
| **53.5** | 0.671 | 0.655 | 0.694 | 0.673 | | 0.685 | 0.686 | 0.686 | 0.686 | | 0.668 | 0.671 | 0.66 | 0.666 |
| **54** | 0.672 | 0.656 | 0.695 | 0.674 | | 0.684 | 0.685 | 0.685 | 0.685 | | 0.667 | 0.671 | 0.66 | 0.666 |
| **54.5** | 0.673 | 0.656 | 0.696 | 0.675 | | 0.687 | 0.688 | 0.688 | 0.688 | | 0.668 | 0.673 | 0.661 | 0.667 |
| **55** | 0.673 | 0.658 | 0.695 | 0.675 | | 0.687 | 0.688 | 0.686 | 0.687 | | 0.669 | 0.674 | 0.663 | 0.669 |
| **55.5** | 0.675 | 0.657 | 0.698 | 0.677 | | 0.688 | 0.688 | 0.688 | 0.688 | | 0.669 | 0.674 | 0.663 | 0.669 |
| **56** | 0.675 | 0.657 | 0.699 | 0.677 | | 0.688 | 0.688 | 0.69 | 0.689 | | 0.67 | 0.675 | 0.665 | 0.670 |
| **56.5** | 0.674 | 0.658 | 0.699 | 0.677 | | 0.689 | 0.689 | 0.69 | 0.689 | | 0.67 | 0.676 | 0.666 | 0.671 |
| **57** | 0.677 | 0.659 | 0.7 | 0.679 | | 0.69 | 0.691 | 0.692 | 0.691 | | 0.671 | 0.677 | 0.668 | 0.672 |
| **57.5** | 0.676 | 0.659 | 0.7 | 0.678 | | 0.69 | 0.69 | 0.691 | 0.690 | | 0.671 | 0.678 | 0.669 | 0.673 |
| **58** | 0.676 | 0.659 | 0.701 | 0.679 | | 0.691 | 0.691 | 0.692 | 0.691 | | 0.671 | 0.678 | 0.668 | 0.672 |
| **58.5** | 0.677 | 0.659 | 0.703 | 0.680 | | 0.691 | 0.69 | 0.692 | 0.691 | | 0.672 | 0.679 | 0.668 | 0.673 |
| **59** | 0.678 | 0.661 | 0.705 | 0.681 | | 0.693 | 0.691 | 0.694 | 0.693 | | 0.673 | 0.68 | 0.671 | 0.675 |
| **59.5** | 0.679 | 0.661 | 0.706 | 0.682 | | 0.694 | 0.693 | 0.695 | 0.694 | | 0.674 | 0.68 | 0.672 | 0.675 |
| **60** | 0.68 | 0.662 | 0.707 | 0.683 | | 0.695 | 0.693 | 0.696 | 0.695 | | 0.675 | 0.681 | 0.672 | 0.676 |
| **60.5** | 0.68 | 0.662 | 0.709 | 0.684 | | 0.694 | 0.695 | 0.698 | 0.696 | | 0.676 | 0.681 | 0.673 | 0.677 |
| **61** | 0.681 | 0.661 | 0.71 | 0.684 | | 0.696 | 0.695 | 0.698 | 0.696 | | 0.676 | 0.682 | 0.674 | 0.677 |
| **61.5** | 0.681 | 0.663 | 0.711 | 0.685 | | 0.695 | 0.696 | 0.698 | 0.696 | | 0.677 | 0.683 | 0.675 | 0.678 |
| **62** | 0.682 | 0.664 | 0.711 | 0.686 | | 0.696 | 0.697 | 0.7 | 0.698 | | 0.678 | 0.684 | 0.677 | 0.680 |
| **62.5** | 0.682 | 0.664 | 0.712 | 0.686 | | 0.697 | 0.697 | 0.699 | 0.698 | | 0.677 | 0.684 | 0.676 | 0.679 |
| **63** | 0.683 | 0.665 | 0.714 | 0.687 | | 0.697 | 0.698 | 0.702 | 0.699 | | 0.679 | 0.684 | 0.678 | 0.680 |
| **63.5** | 0.683 | 0.666 | 0.713 | 0.687 | | 0.699 | 0.699 | 0.7 | 0.699 | | 0.68 | 0.685 | 0.679 | 0.681 |
| **64** | 0.684 | 0.667 | 0.715 | 0.689 | | 0.698 | 0.699 | 0.703 | 0.700 | | 0.681 | 0.686 | 0.68 | 0.682 |
| **64.5** | 0.685 | 0.668 | 0.715 | 0.689 | | 0.698 | 0.699 | 0.704 | 0.700 | | 0.682 | 0.687 | 0.681 | 0.683 |
| **65** | 0.685 | 0.67 | 0.716 | 0.690 | | 0.699 | 0.7 | 0.704 | 0.701 | | 0.683 | 0.688 | 0.682 | 0.684 |
| **65.5** | 0.685 | 0.67 | 0.716 | 0.690 | | 0.701 | 0.699 | 0.706 | 0.702 | | 0.683 | 0.687 | 0.682 | 0.684 |
| **66** | 0.685 | 0.672 | 0.717 | 0.691 | | 0.7 | 0.701 | 0.707 | 0.703 | | 0.685 | 0.689 | 0.684 | 0.686 |
| **66.5** | 0.686 | 0.673 | 0.718 | 0.692 | | 0.7 | 0.701 | 0.707 | 0.703 | | 0.685 | 0.689 | 0.684 | 0.686 |
| **67** | 0.686 | 0.677 | 0.718 | 0.694 | | 0.702 | 0.702 | 0.708 | 0.704 | | 0.686 | 0.69 | 0.685 | 0.687 |
| **67.5** | 0.687 | 0.678 | 0.72 | 0.695 | | 0.701 | 0.703 | 0.707 | 0.704 | | 0.687 | 0.69 | 0.687 | 0.688 |
| **68** | 0.687 | 0.679 | 0.72 | 0.695 | | 0.702 | 0.703 | 0.709 | 0.705 | | 0.688 | 0.69 | 0.686 | 0.688 |
| **68.5** | 0.688 | 0.678 | 0.721 | 0.696 | | 0.702 | 0.704 | 0.71 | 0.705 | | 0.689 | 0.692 | 0.688 | 0.690 |
| **69** | 0.688 | 0.688 | 0.721 | 0.699 | | 0.702 | 0.703 | 0.71 | 0.705 | | 0.69 | 0.692 | 0.688 | 0.690 |
| **69.5** | 0.688 | 0.691 | 0.722 | 0.700 | | 0.702 | 0.703 | 0.711 | 0.705 | | 0.69 | 0.693 | 0.689 | 0.691 |
| **70** | 0.688 | 0.691 | 0.721 | 0.700 | | 0.703 | 0.703 | 0.711 | 0.706 | | 0.692 | 0.694 | 0.689 | 0.692 |
| **70.5** | 0.69 | 0.688 | 0.723 | 0.700 | | 0.703 | 0.702 | 0.713 | 0.706 | | 0.692 | 0.695 | 0.691 | 0.693 |
| **71** | 0.689 | 0.686 | 0.722 | 0.699 | | 0.704 | 0.702 | 0.713 | 0.706 | | 0.693 | 0.695 | 0.691 | 0.693 |
| **71.5** | 0.689 | 0.686 | 0.722 | 0.699 | | 0.704 | 0.701 | 0.714 | 0.706 | | 0.694 | 0.695 | 0.693 | 0.694 |

| | | | | | | | | | | | | | | |
|---|---|---|---|---|---|---|---|---|---|---|---|---|---|---|
| **72** | 0.689 | 0.683 | 0.722 | 0.698 | | 0.704 | 0.701 | 0.714 | 0.706 | | 0.694 | 0.697 | 0.693 | 0.695 |
| **72.5** | 0.69 | 0.677 | 0.723 | 0.697 | | 0.705 | 0.701 | 0.714 | 0.707 | | 0.695 | 0.697 | 0.694 | 0.695 |

**Supplementary Table 5C2. Raw growth data measured for the deletion strain (Δ*s479*) at standard zinc concentration.** For each time point, $OD_{650nm}$ of three biological (replicate 1-3; s479-1 to s479-3), with three technical replicates (A to C) each, are given. Additionally, the mean $OD_{650nm}$ over all technical replicates is given for each biological replicate.

| | **replicate 1** | | | | | **replicate 2** | | | | | **replicate 3** | | | |
|---|---|---|---|---|---|---|---|---|---|---|---|---|---|---|
| | **s479-1A** | **s479-1B** | **s479-1C** | **s479-1** | | **s479-2A** | **s479-2B** | **s479-2C** | **s479-2** | | **s479-3A** | **s479-3B** | **s479-3C** | **s479-3** |
| **time [h]** | OD650 | OD650 | OD650 | **Mean OD650** | | OD650 | OD650 | OD650 | **Mean OD650** | | OD650 | OD650 | OD650 | **Mean OD650** |
| **0.5** | 0.064 | 0.065 | 0.065 | 0.065 | | 0.056 | 0.054 | 0.056 | 0.055 | | 0.067 | 0.063 | 0.062 | 0.064 |
| **1** | 0.066 | 0.065 | 0.065 | 0.065 | | 0.061 | 0.059 | 0.061 | 0.060 | | 0.074 | 0.071 | 0.069 | 0.071 |
| **1.5** | 0.08 | 0.08 | 0.08 | 0.080 | | 0.069 | 0.067 | 0.068 | 0.068 | | 0.085 | 0.082 | 0.079 | 0.082 |
| **2** | 0.091 | 0.092 | 0.091 | 0.091 | | 0.077 | 0.074 | 0.077 | 0.076 | | 0.095 | 0.091 | 0.089 | 0.092 |
| **2.5** | 0.1 | 0.101 | 0.101 | 0.101 | | 0.087 | 0.084 | 0.087 | 0.086 | | 0.103 | 0.1 | 0.098 | 0.100 |
| **3** | 0.109 | 0.111 | 0.11 | 0.110 | | 0.095 | 0.092 | 0.095 | 0.094 | | 0.111 | 0.109 | 0.106 | 0.109 |
| **3.5** | 0.119 | 0.12 | 0.121 | 0.120 | | 0.102 | 0.099 | 0.103 | 0.101 | | 0.12 | 0.118 | 0.115 | 0.118 |
| **4** | 0.13 | 0.132 | 0.133 | 0.132 | | 0.11 | 0.107 | 0.111 | 0.109 | | 0.13 | 0.127 | 0.124 | 0.127 |
| **4.5** | 0.141 | 0.143 | 0.143 | 0.142 | | 0.12 | 0.117 | 0.122 | 0.120 | | 0.142 | 0.139 | 0.132 | 0.138 |
| **5** | 0.154 | 0.155 | 0.155 | 0.155 | | 0.13 | 0.128 | 0.131 | 0.130 | | 0.151 | 0.148 | 0.144 | 0.148 |
| **5.5** | 0.167 | 0.167 | 0.167 | 0.167 | | 0.14 | 0.139 | 0.142 | 0.140 | | 0.164 | 0.16 | 0.158 | 0.161 |
| **6** | 0.178 | 0.18 | 0.181 | 0.180 | | 0.151 | 0.148 | 0.155 | 0.151 | | 0.178 | 0.173 | 0.171 | 0.174 |
| **6.5** | 0.191 | 0.192 | 0.193 | 0.192 | | 0.161 | 0.16 | 0.166 | 0.162 | | 0.19 | 0.186 | 0.185 | 0.187 |
| **7** | 0.204 | 0.206 | 0.205 | 0.205 | | 0.173 | 0.171 | 0.179 | 0.174 | | 0.202 | 0.203 | 0.198 | 0.201 |
| **7.5** | 0.218 | 0.22 | 0.221 | 0.220 | | 0.186 | 0.183 | 0.191 | 0.187 | | 0.215 | 0.216 | 0.213 | 0.215 |
| **8** | 0.231 | 0.234 | 0.237 | 0.234 | | 0.197 | 0.195 | 0.203 | 0.198 | | 0.228 | 0.228 | 0.228 | 0.228 |
| **8.5** | 0.248 | 0.25 | 0.255 | 0.251 | | 0.208 | 0.207 | 0.215 | 0.210 | | 0.242 | 0.245 | 0.242 | 0.243 |
| **9** | 0.26 | 0.264 | 0.271 | 0.265 | | 0.219 | 0.219 | 0.229 | 0.222 | | 0.255 | 0.259 | 0.255 | 0.256 |
| **9.5** | 0.273 | 0.278 | 0.289 | 0.280 | | 0.231 | 0.231 | 0.242 | 0.235 | | 0.27 | 0.273 | 0.268 | 0.270 |
| **10** | 0.286 | 0.289 | 0.303 | 0.293 | | 0.242 | 0.244 | 0.254 | 0.247 | | 0.282 | 0.283 | 0.281 | 0.282 |
| **10.5** | 0.299 | 0.302 | 0.317 | 0.306 | | 0.255 | 0.257 | 0.267 | 0.260 | | 0.294 | 0.297 | 0.293 | 0.295 |
| **11** | 0.31 | 0.312 | 0.327 | 0.316 | | 0.267 | 0.267 | 0.279 | 0.271 | | 0.307 | 0.308 | 0.306 | 0.307 |
| **11.5** | 0.323 | 0.326 | 0.341 | 0.330 | | 0.278 | 0.28 | 0.29 | 0.283 | | 0.319 | 0.317 | 0.315 | 0.317 |
| **12** | 0.335 | 0.338 | 0.353 | 0.342 | | 0.289 | 0.292 | 0.301 | 0.294 | | 0.329 | 0.328 | 0.327 | 0.328 |
| **12.5** | 0.345 | 0.348 | 0.362 | 0.352 | | 0.3 | 0.302 | 0.312 | 0.305 | | 0.339 | 0.336 | 0.335 | 0.337 |
| **13** | 0.355 | 0.358 | 0.371 | 0.361 | | 0.311 | 0.312 | 0.322 | 0.315 | | 0.349 | 0.345 | 0.346 | 0.347 |
| **13.5** | 0.365 | 0.367 | 0.38 | 0.371 | | 0.321 | 0.322 | 0.331 | 0.325 | | 0.359 | 0.353 | 0.353 | 0.355 |
| **14** | 0.374 | 0.375 | 0.387 | 0.379 | | 0.329 | 0.33 | 0.339 | 0.333 | | 0.367 | 0.36 | 0.361 | 0.363 |
| **14.5** | 0.381 | 0.383 | 0.396 | 0.387 | | 0.337 | 0.338 | 0.347 | 0.341 | | 0.377 | 0.368 | 0.369 | 0.371 |
| **15** | 0.39 | 0.391 | 0.404 | 0.395 | | 0.347 | 0.347 | 0.356 | 0.350 | | 0.386 | 0.376 | 0.376 | 0.379 |
| **15.5** | 0.397 | 0.398 | 0.409 | 0.401 | | 0.354 | 0.355 | 0.363 | 0.357 | | 0.392 | 0.383 | 0.385 | 0.387 |
| **16** | 0.403 | 0.402 | 0.415 | 0.407 | | 0.362 | 0.363 | 0.372 | 0.366 | | 0.4 | 0.39 | 0.392 | 0.394 |
| **16.5** | 0.409 | 0.409 | 0.421 | 0.413 | | 0.37 | 0.371 | 0.379 | 0.373 | | 0.406 | 0.397 | 0.401 | 0.401 |
| **17** | 0.416 | 0.417 | 0.428 | 0.420 | | 0.378 | 0.377 | 0.387 | 0.381 | | 0.414 | 0.405 | 0.409 | 0.409 |
| **17.5** | 0.419 | 0.423 | 0.432 | 0.425 | | 0.385 | 0.385 | 0.394 | 0.388 | | 0.421 | 0.411 | 0.415 | 0.416 |
| **18** | 0.428 | 0.427 | 0.439 | 0.431 | | 0.392 | 0.393 | 0.399 | 0.395 | | 0.428 | 0.419 | 0.423 | 0.423 |
| **18.5** | 0.433 | 0.434 | 0.443 | 0.437 | | 0.398 | 0.399 | 0.406 | 0.401 | | 0.433 | 0.424 | 0.431 | 0.429 |
| **19** | 0.437 | 0.439 | 0.448 | 0.441 | | 0.405 | 0.406 | 0.413 | 0.408 | | 0.44 | 0.432 | 0.439 | 0.437 |
| **19.5** | 0.442 | 0.444 | 0.455 | 0.447 | | 0.411 | 0.412 | 0.419 | 0.414 | | 0.447 | 0.439 | 0.447 | 0.444 |
| **20** | 0.448 | 0.448 | 0.459 | 0.452 | | 0.417 | 0.417 | 0.424 | 0.419 | | 0.453 | 0.445 | 0.453 | 0.450 |
| **20.5** | 0.454 | 0.455 | 0.463 | 0.457 | | 0.423 | 0.424 | 0.43 | 0.426 | | 0.459 | 0.451 | 0.46 | 0.457 |

| | | | | | | | | | | | | | | |
|---|---|---|---|---|---|---|---|---|---|---|---|---|---|---|
| **21** | 0.458 | 0.459 | 0.468 | 0.462 | | 0.429 | 0.429 | 0.435 | 0.431 | | 0.464 | 0.459 | 0.467 | 0.463 |
| **21.5** | 0.463 | 0.465 | 0.474 | 0.467 | | 0.434 | 0.435 | 0.441 | 0.437 | | 0.47 | 0.463 | 0.473 | 0.469 |
| **22** | 0.469 | 0.472 | 0.48 | 0.474 | | 0.44 | 0.442 | 0.447 | 0.443 | | 0.476 | 0.471 | 0.48 | 0.476 |
| **22.5** | 0.474 | 0.476 | 0.483 | 0.478 | | 0.446 | 0.446 | 0.451 | 0.448 | | 0.481 | 0.476 | 0.485 | 0.481 |
| **23** | 0.477 | 0.48 | 0.488 | 0.482 | | 0.451 | 0.451 | 0.457 | 0.453 | | 0.486 | 0.481 | 0.491 | 0.486 |
| **23.5** | 0.482 | 0.485 | 0.493 | 0.487 | | 0.455 | 0.456 | 0.462 | 0.458 | | 0.491 | 0.486 | 0.497 | 0.491 |
| **24** | 0.486 | 0.489 | 0.496 | 0.490 | | 0.46 | 0.461 | 0.467 | 0.463 | | 0.495 | 0.491 | 0.502 | 0.496 |
| **24.5** | 0.491 | 0.493 | 0.5 | 0.495 | | 0.466 | 0.466 | 0.472 | 0.468 | | 0.499 | 0.496 | 0.508 | 0.501 |
| **25** | 0.495 | 0.498 | 0.504 | 0.499 | | 0.47 | 0.47 | 0.476 | 0.472 | | 0.503 | 0.501 | 0.513 | 0.506 |
| **25.5** | 0.499 | 0.502 | 0.507 | 0.503 | | 0.474 | 0.474 | 0.48 | 0.476 | | 0.508 | 0.506 | 0.518 | 0.511 |
| **26** | 0.502 | 0.505 | 0.511 | 0.506 | | 0.477 | 0.476 | 0.484 | 0.479 | | 0.512 | 0.51 | 0.523 | 0.515 |
| **26.5** | 0.507 | 0.509 | 0.515 | 0.510 | | 0.482 | 0.482 | 0.489 | 0.484 | | 0.514 | 0.515 | 0.528 | 0.519 |
| **27** | 0.511 | 0.512 | 0.519 | 0.514 | | 0.485 | 0.485 | 0.492 | 0.487 | | 0.519 | 0.519 | 0.532 | 0.523 |
| **27.5** | 0.514 | 0.516 | 0.521 | 0.517 | | 0.489 | 0.488 | 0.495 | 0.491 | | 0.521 | 0.523 | 0.536 | 0.527 |
| **28** | 0.517 | 0.518 | 0.523 | 0.519 | | 0.492 | 0.492 | 0.497 | 0.494 | | 0.525 | 0.526 | 0.54 | 0.530 |
| **28.5** | 0.52 | 0.521 | 0.526 | 0.522 | | 0.497 | 0.496 | 0.501 | 0.498 | | 0.53 | 0.53 | 0.545 | 0.535 |
| **29** | 0.523 | 0.524 | 0.528 | 0.525 | | 0.5 | 0.498 | 0.504 | 0.501 | | 0.533 | 0.533 | 0.548 | 0.538 |
| **29.5** | 0.526 | 0.527 | 0.531 | 0.528 | | 0.503 | 0.502 | 0.507 | 0.504 | | 0.536 | 0.537 | 0.552 | 0.542 |
| **30** | 0.529 | 0.529 | 0.532 | 0.530 | | 0.508 | 0.506 | 0.51 | 0.508 | | 0.54 | 0.539 | 0.556 | 0.545 |
| **30.5** | 0.533 | 0.533 | 0.535 | 0.534 | | 0.511 | 0.51 | 0.513 | 0.511 | | 0.543 | 0.543 | 0.56 | 0.549 |
| **31** | 0.536 | 0.535 | 0.538 | 0.536 | | 0.514 | 0.513 | 0.515 | 0.514 | | 0.546 | 0.546 | 0.562 | 0.551 |
| **31.5** | 0.538 | 0.537 | 0.54 | 0.538 | | 0.517 | 0.516 | 0.518 | 0.517 | | 0.548 | 0.549 | 0.567 | 0.555 |
| **32** | 0.541 | 0.541 | 0.543 | 0.542 | | 0.521 | 0.519 | 0.522 | 0.521 | | 0.552 | 0.554 | 0.57 | 0.559 |
| **32.5** | 0.543 | 0.542 | 0.543 | 0.543 | | 0.524 | 0.522 | 0.524 | 0.523 | | 0.555 | 0.554 | 0.573 | 0.561 |
| **33** | 0.546 | 0.545 | 0.546 | 0.546 | | 0.526 | 0.525 | 0.525 | 0.525 | | 0.556 | 0.557 | 0.575 | 0.563 |
| **33.5** | 0.548 | 0.547 | 0.548 | 0.548 | | 0.529 | 0.527 | 0.528 | 0.528 | | 0.559 | 0.56 | 0.579 | 0.566 |
| **34** | 0.55 | 0.55 | 0.55 | 0.550 | | 0.533 | 0.53 | 0.531 | 0.531 | | 0.561 | 0.563 | 0.581 | 0.568 |
| **34.5** | 0.553 | 0.552 | 0.553 | 0.553 | | 0.536 | 0.533 | 0.534 | 0.534 | | 0.564 | 0.566 | 0.585 | 0.572 |
| **35** | 0.556 | 0.555 | 0.555 | 0.555 | | 0.538 | 0.535 | 0.535 | 0.536 | | 0.567 | 0.568 | 0.588 | 0.574 |
| **35.5** | 0.557 | 0.555 | 0.557 | 0.556 | | 0.541 | 0.538 | 0.538 | 0.539 | | 0.569 | 0.571 | 0.592 | 0.577 |
| **36** | 0.558 | 0.558 | 0.559 | 0.558 | | 0.544 | 0.539 | 0.539 | 0.541 | | 0.572 | 0.574 | 0.594 | 0.580 |
| **36.5** | 0.562 | 0.561 | 0.56 | 0.561 | | 0.546 | 0.542 | 0.542 | 0.543 | | 0.573 | 0.577 | 0.598 | 0.583 |
| **37** | 0.564 | 0.563 | 0.562 | 0.563 | | 0.548 | 0.544 | 0.544 | 0.545 | | 0.576 | 0.58 | 0.601 | 0.586 |
| **37.5** | 0.566 | 0.565 | 0.564 | 0.565 | | 0.55 | 0.546 | 0.546 | 0.547 | | 0.577 | 0.582 | 0.602 | 0.587 |
| **38** | 0.568 | 0.567 | 0.566 | 0.567 | | 0.553 | 0.548 | 0.548 | 0.550 | | 0.582 | 0.584 | 0.606 | 0.591 |
| **38.5** | 0.57 | 0.567 | 0.566 | 0.568 | | 0.555 | 0.551 | 0.552 | 0.553 | | 0.581 | 0.586 | 0.61 | 0.592 |
| **39** | 0.572 | 0.57 | 0.569 | 0.570 | | 0.557 | 0.553 | 0.553 | 0.554 | | 0.584 | 0.588 | 0.613 | 0.595 |
| **39.5** | 0.575 | 0.573 | 0.572 | 0.573 | | 0.559 | 0.555 | 0.555 | 0.556 | | 0.587 | 0.59 | 0.616 | 0.598 |
| **40** | 0.577 | 0.575 | 0.572 | 0.575 | | 0.561 | 0.557 | 0.557 | 0.558 | | 0.588 | 0.594 | 0.619 | 0.600 |
| **40.5** | 0.579 | 0.576 | 0.575 | 0.577 | | 0.563 | 0.559 | 0.56 | 0.561 | | 0.589 | 0.595 | 0.623 | 0.602 |
| **41** | 0.579 | 0.576 | 0.576 | 0.577 | | 0.564 | 0.561 | 0.562 | 0.562 | | 0.593 | 0.596 | 0.629 | 0.606 |
| **41.5** | 0.583 | 0.58 | 0.577 | 0.580 | | 0.565 | 0.563 | 0.565 | 0.564 | | 0.592 | 0.602 | 0.635 | 0.610 |
| **42** | 0.585 | 0.581 | 0.58 | 0.582 | | 0.567 | 0.564 | 0.567 | 0.566 | | 0.596 | 0.601 | 0.625 | 0.607 |
| **42.5** | 0.588 | 0.583 | 0.581 | 0.584 | | 0.569 | 0.566 | 0.569 | 0.568 | | 0.597 | 0.603 | 0.629 | 0.610 |
| **43** | 0.589 | 0.583 | 0.583 | 0.585 | | 0.571 | 0.569 | 0.571 | 0.570 | | 0.599 | 0.608 | 0.636 | 0.614 |
| **43.5** | 0.591 | 0.586 | 0.582 | 0.586 | | 0.573 | 0.569 | 0.572 | 0.571 | | 0.601 | 0.609 | 0.641 | 0.617 |
| **44** | 0.592 | 0.588 | 0.584 | 0.588 | | 0.575 | 0.572 | 0.574 | 0.574 | | 0.603 | 0.609 | 0.646 | 0.619 |
| **44.5** | 0.595 | 0.589 | 0.587 | 0.590 | | 0.576 | 0.574 | 0.577 | 0.576 | | 0.605 | 0.613 | 0.654 | 0.624 |
| **45** | 0.596 | 0.591 | 0.588 | 0.592 | | 0.578 | 0.575 | 0.577 | 0.577 | | 0.607 | 0.613 | 0.661 | 0.627 |
| **45.5** | 0.599 | 0.593 | 0.591 | 0.594 | | 0.58 | 0.577 | 0.579 | 0.579 | | 0.607 | 0.614 | 0.668 | 0.630 |
| **46** | 0.601 | 0.595 | 0.593 | 0.596 | | 0.581 | 0.579 | 0.581 | 0.580 | | 0.607 | 0.616 | 0.674 | 0.632 |

| | | | | | | | | | | | | | | |
|---|---|---|---|---|---|---|---|---|---|---|---|---|---|---|
| **46.5** | 0.602 | 0.596 | 0.592 | 0.597 | | 0.584 | 0.58 | 0.583 | 0.582 | | 0.608 | 0.619 | 0.677 | 0.635 |
| **47** | 0.604 | 0.597 | 0.595 | 0.599 | | 0.586 | 0.582 | 0.585 | 0.584 | | 0.61 | 0.619 | 0.679 | 0.636 |
| **47.5** | 0.605 | 0.598 | 0.596 | 0.600 | | 0.587 | 0.585 | 0.587 | 0.586 | | 0.611 | 0.621 | 0.68 | 0.637 |
| **48** | 0.607 | 0.601 | 0.598 | 0.602 | | 0.589 | 0.586 | 0.588 | 0.588 | | 0.611 | 0.621 | 0.681 | 0.638 |
| **48.5** | 0.607 | 0.601 | 0.599 | 0.602 | | 0.59 | 0.588 | 0.591 | 0.590 | | 0.613 | 0.62 | 0.684 | 0.639 |
| **49** | 0.609 | 0.603 | 0.601 | 0.604 | | 0.593 | 0.591 | 0.592 | 0.592 | | 0.615 | 0.622 | 0.685 | 0.641 |
| **49.5** | 0.611 | 0.605 | 0.602 | 0.606 | | 0.595 | 0.592 | 0.594 | 0.594 | | 0.617 | 0.621 | 0.685 | 0.641 |
| **50** | 0.612 | 0.606 | 0.603 | 0.607 | | 0.596 | 0.594 | 0.594 | 0.595 | | 0.618 | 0.624 | 0.683 | 0.642 |
| **50.5** | 0.613 | 0.608 | 0.604 | 0.608 | | 0.598 | 0.596 | 0.596 | 0.597 | | 0.619 | 0.624 | 0.678 | 0.640 |
| **51** | 0.613 | 0.607 | 0.606 | 0.609 | | 0.6 | 0.597 | 0.596 | 0.598 | | 0.62 | 0.626 | 0.673 | 0.640 |
| **51.5** | 0.615 | 0.609 | 0.607 | 0.610 | | 0.601 | 0.598 | 0.597 | 0.599 | | 0.622 | 0.625 | 0.666 | 0.638 |
| **52** | 0.616 | 0.61 | 0.608 | 0.611 | | 0.603 | 0.599 | 0.598 | 0.600 | | 0.621 | 0.627 | 0.662 | 0.637 |
| **52.5** | 0.617 | 0.611 | 0.608 | 0.612 | | 0.604 | 0.6 | 0.599 | 0.601 | | 0.624 | 0.626 | 0.663 | 0.638 |
| **53** | 0.618 | 0.612 | 0.61 | 0.613 | | 0.605 | 0.601 | 0.599 | 0.602 | | 0.624 | 0.627 | 0.664 | 0.638 |
| **53.5** | 0.618 | 0.612 | 0.61 | 0.613 | | 0.606 | 0.602 | 0.601 | 0.603 | | 0.625 | 0.627 | 0.668 | 0.640 |
| **54** | 0.618 | 0.612 | 0.61 | 0.613 | | 0.607 | 0.603 | 0.6 | 0.603 | | 0.626 | 0.628 | 0.668 | 0.641 |
| **54.5** | 0.62 | 0.614 | 0.611 | 0.615 | | 0.607 | 0.604 | 0.602 | 0.604 | | 0.627 | 0.63 | 0.668 | 0.642 |
| **55** | 0.62 | 0.614 | 0.612 | 0.615 | | 0.608 | 0.605 | 0.603 | 0.605 | | 0.627 | 0.629 | 0.669 | 0.642 |
| **55.5** | 0.62 | 0.615 | 0.612 | 0.616 | | 0.609 | 0.606 | 0.603 | 0.606 | | 0.628 | 0.63 | 0.674 | 0.644 |
| **56** | 0.622 | 0.615 | 0.613 | 0.617 | | 0.609 | 0.607 | 0.603 | 0.606 | | 0.63 | 0.63 | 0.678 | 0.646 |
| **56.5** | 0.622 | 0.616 | 0.613 | 0.617 | | 0.61 | 0.607 | 0.604 | 0.607 | | 0.63 | 0.631 | 0.682 | 0.648 |
| **57** | 0.622 | 0.616 | 0.615 | 0.618 | | 0.612 | 0.608 | 0.605 | 0.608 | | 0.631 | 0.632 | 0.685 | 0.649 |
| **57.5** | 0.621 | 0.616 | 0.614 | 0.617 | | 0.612 | 0.609 | 0.606 | 0.609 | | 0.632 | 0.632 | 0.689 | 0.651 |
| **58** | 0.622 | 0.617 | 0.615 | 0.618 | | 0.612 | 0.609 | 0.607 | 0.609 | | 0.632 | 0.634 | 0.693 | 0.653 |
| **58.5** | 0.623 | 0.618 | 0.615 | 0.619 | | 0.613 | 0.609 | 0.608 | 0.610 | | 0.634 | 0.632 | 0.696 | 0.654 |
| **59** | 0.623 | 0.617 | 0.616 | 0.619 | | 0.613 | 0.61 | 0.609 | 0.611 | | 0.634 | 0.634 | 0.698 | 0.655 |
| **59.5** | 0.622 | 0.618 | 0.616 | 0.619 | | 0.613 | 0.611 | 0.61 | 0.611 | | 0.635 | 0.635 | 0.701 | 0.657 |
| **60** | 0.622 | 0.619 | 0.616 | 0.619 | | 0.614 | 0.612 | 0.61 | 0.612 | | 0.636 | 0.637 | 0.704 | 0.659 |
| **60.5** | 0.622 | 0.619 | 0.617 | 0.619 | | 0.614 | 0.613 | 0.611 | 0.613 | | 0.636 | 0.638 | 0.702 | 0.659 |
| **61** | 0.622 | 0.619 | 0.617 | 0.619 | | 0.615 | 0.613 | 0.611 | 0.613 | | 0.637 | 0.64 | 0.688 | 0.655 |
| **61.5** | 0.623 | 0.62 | 0.617 | 0.620 | | 0.615 | 0.613 | 0.611 | 0.613 | | 0.638 | 0.64 | 0.705 | 0.661 |
| **62** | 0.623 | 0.62 | 0.617 | 0.620 | | 0.615 | 0.614 | 0.612 | 0.614 | | 0.638 | 0.64 | 0.714 | 0.664 |
| **62.5** | 0.624 | 0.621 | 0.619 | 0.621 | | 0.615 | 0.613 | 0.613 | 0.614 | | 0.639 | 0.64 | 0.72 | 0.666 |
| **63** | 0.624 | 0.621 | 0.619 | 0.621 | | 0.616 | 0.614 | 0.613 | 0.614 | | 0.64 | 0.64 | 0.725 | 0.668 |
| **63.5** | 0.624 | 0.622 | 0.619 | 0.622 | | 0.616 | 0.614 | 0.613 | 0.614 | | 0.641 | 0.642 | 0.725 | 0.669 |
| **64** | 0.624 | 0.623 | 0.62 | 0.622 | | 0.615 | 0.615 | 0.614 | 0.615 | | 0.641 | 0.643 | 0.724 | 0.669 |
| **64.5** | 0.625 | 0.623 | 0.62 | 0.623 | | 0.616 | 0.616 | 0.614 | 0.615 | | 0.641 | 0.643 | 0.725 | 0.670 |
| **65** | 0.625 | 0.624 | 0.62 | 0.623 | | 0.617 | 0.616 | 0.615 | 0.616 | | 0.643 | 0.645 | 0.726 | 0.671 |
| **65.5** | 0.625 | 0.623 | 0.621 | 0.623 | | 0.617 | 0.616 | 0.615 | 0.616 | | 0.643 | 0.646 | 0.725 | 0.671 |
| **66** | 0.625 | 0.624 | 0.621 | 0.623 | | 0.618 | 0.617 | 0.616 | 0.617 | | 0.643 | 0.645 | 0.725 | 0.671 |
| **66.5** | 0.626 | 0.624 | 0.62 | 0.623 | | 0.618 | 0.617 | 0.616 | 0.617 | | 0.644 | 0.647 | 0.724 | 0.672 |
| **67** | 0.626 | 0.625 | 0.622 | 0.624 | | 0.619 | 0.617 | 0.616 | 0.617 | | 0.645 | 0.649 | 0.722 | 0.672 |
| **67.5** | 0.626 | 0.624 | 0.622 | 0.624 | | 0.619 | 0.618 | 0.617 | 0.618 | | 0.644 | 0.648 | 0.72 | 0.671 |
| **68** | 0.626 | 0.625 | 0.621 | 0.624 | | 0.62 | 0.619 | 0.617 | 0.619 | | 0.645 | 0.647 | 0.72 | 0.671 |
| **68.5** | 0.627 | 0.625 | 0.623 | 0.625 | | 0.62 | 0.619 | 0.618 | 0.619 | | 0.645 | 0.649 | 0.717 | 0.670 |
| **69** | 0.627 | 0.626 | 0.622 | 0.625 | | 0.62 | 0.619 | 0.617 | 0.619 | | 0.646 | 0.648 | 0.715 | 0.670 |
| **69.5** | 0.627 | 0.626 | 0.623 | 0.625 | | 0.621 | 0.62 | 0.618 | 0.620 | | 0.646 | 0.646 | 0.714 | 0.669 |
| **70** | 0.628 | 0.626 | 0.623 | 0.626 | | 0.622 | 0.62 | 0.619 | 0.620 | | 0.646 | 0.645 | 0.713 | 0.668 |
| **70.5** | 0.628 | 0.627 | 0.623 | 0.626 | | 0.622 | 0.621 | 0.618 | 0.620 | | 0.645 | 0.645 | 0.713 | 0.668 |
| **71** | 0.628 | 0.627 | 0.623 | 0.626 | | 0.622 | 0.621 | 0.619 | 0.621 | | 0.647 | 0.646 | 0.712 | 0.668 |
| **71.5** | 0.628 | 0.627 | 0.623 | 0.626 | | 0.623 | 0.622 | 0.619 | 0.621 | | 0.647 | 0.643 | 0.712 | 0.667 |
| **72** | 0.628 | 0.627 | 0.622 | 0.626 | | 0.624 | 0.622 | 0.62 | 0.622 | | 0.647 | 0.644 | 0.711 | 0.667 |

| | | | | | | | | | | | | | | |
|---|---|---|---|---|---|---|---|---|---|---|---|---|---|---|
| **72.5** | 0.628 | 0.627 | 0.624 | 0.626 | | 0.624 | 0.621 | 0.619 | 0.621 | | 0.647 | 0.644 | 0.711 | 0.667 |

**Supplementary Table 6. Data and calculations for growth experiments at high zinc concentration.** Supplementary Table 6A1. and A2. list the data used to compile the growth curves in Supplementary Figure 5A. Supplementary Table 6B. shows the calculations of growth rate and doubling time for wildtype and deletion strain ($\Delta$*s479*). Supplementary Table 6C1. and C2. list all the raw data collected for the growth experiments at high zinc concentration. For better readability, all numbers were cut to three decimals or less. **Supplementary Table 6A1. Mean growth and standard deviation used to compile the wildtype strain (H66) growth curve in Supplementary Figure 5A.** The mean $OD_{650nm}$ is given for all three biological replicates of wildtype strain H66 (H66-1 to H66-3) alongside the overall mean $OD_{650nm}$ and standard deviation per time interval.

| **H66-1** | **H66-2** | **H66-3** | **H66** | **H66** | |
|---|---|---|---|---|---|
| **Mean OD650** | **Mean OD650** | **Mean OD650** | **Mean OD650** | **SD** | **time [h]** |
| 0.066 | 0.064 | 0.062 | **0.064** | 0.002 | **0.5** |
| 0.090 | 0.093 | 0.093 | **0.092** | 0.001 | **2.5** |
| 0.122 | 0.129 | 0.131 | **0.127** | 0.004 | **4.5** |
| 0.151 | 0.165 | 0.171 | **0.162** | 0.008 | **6.5** |
| 0.172 | 0.200 | 0.213 | **0.195** | 0.017 | **8.5** |
| 0.196 | 0.239 | 0.257 | **0.231** | 0.026 | **10.5** |
| 0.220 | 0.285 | 0.299 | **0.268** | 0.035 | **12.5** |
| 0.245 | 0.339 | 0.347 | **0.310** | 0.046 | **14.5** |
| 0.274 | 0.393 | 0.394 | **0.354** | 0.056 | **16.5** |
| 0.308 | 0.444 | 0.447 | **0.400** | 0.065 | **18.5** |
| 0.357 | 0.496 | 0.495 | **0.449** | 0.065 | **20.5** |
| 0.423 | 0.538 | 0.537 | **0.499** | 0.054 | **22.5** |
| 0.489 | 0.573 | 0.573 | **0.545** | 0.039 | **24.5** |
| 0.550 | 0.601 | 0.604 | **0.585** | 0.025 | **26.5** |
| 0.599 | 0.626 | 0.629 | **0.618** | 0.014 | **28.5** |
| 0.637 | 0.648 | 0.651 | **0.645** | 0.006 | **30.5** |
| 0.670 | 0.668 | 0.672 | **0.670** | 0.001 | **32.5** |
| 0.697 | 0.686 | 0.691 | **0.692** | 0.005 | **34.5** |
| 0.721 | 0.704 | 0.710 | **0.712** | 0.007 | **36.5** |
| 0.742 | 0.717 | 0.728 | **0.729** | 0.010 | **38.5** |
| 0.760 | 0.730 | 0.741 | **0.744** | 0.013 | **40.5** |
| 0.775 | 0.741 | 0.754 | **0.757** | 0.014 | **42.5** |
| 0.784 | 0.746 | 0.763 | **0.765** | 0.016 | **44.5** |
| 0.791 | 0.753 | 0.771 | **0.771** | 0.016 | **46.5** |
| 0.798 | 0.758 | 0.775 | **0.777** | 0.017 | **48.5** |
| 0.802 | 0.763 | 0.781 | **0.782** | 0.016 | **50.5** |
| 0.806 | 0.768 | 0.788 | **0.787** | 0.015 | **52.5** |
| 0.808 | 0.775 | 0.794 | **0.793** | 0.014 | **54.5** |
| 0.814 | 0.781 | 0.800 | **0.798** | 0.013 | **56.5** |
| 0.815 | 0.786 | 0.805 | **0.802** | 0.012 | **58.5** |
| 0.820 | 0.792 | 0.812 | **0.808** | 0.012 | **60.5** |
| 0.823 | 0.796 | 0.817 | **0.812** | 0.012 | **62.5** |
| 0.827 | 0.802 | 0.822 | **0.817** | 0.011 | **64.5** |
| 0.831 | 0.807 | 0.827 | **0.822** | 0.011 | **66.5** |
| 0.834 | 0.812 | 0.832 | **0.826** | 0.010 | **68.5** |
| 0.839 | 0.817 | 0.837 | **0.831** | 0.010 | **70.5** |

| | | | | | |
|---|---|---|---|---|---|
| | 0.821 | 0.842 | **0.831** | 0.010 | **72.5** |

**Supplementary Table 6A2. Mean growth and standard deviation used to compile the deletion strain (Δ*s479*) growth curve in Supplementary Figure 5A.** The mean $OD_{650nm}$ is given for all three biological replicates of deletion strain Δ*s479* (s479-1 to s479-3) alongside the overall mean $OD_{650nm}$ and standard deviation per time interval.

| **s479-1** | **s479-2** | **s479-3** | **s479** | **s479 SD** | |
|---|---|---|---|---|---|
| **Mean OD650** | **Mean OD650** | **Mean OD650** | **Mean OD650** | **SD** | **time [h]** |
| 0.066 | 0.063 | 0.061 | **0.063** | 0.002 | **0.5** |
| 0.078 | 0.075 | 0.075 | **0.076** | 0.001 | **2.5** |
| 0.095 | 0.094 | 0.091 | **0.093** | 0.002 | **4.5** |
| 0.112 | 0.116 | 0.107 | **0.112** | 0.003 | **6.5** |
| 0.129 | 0.136 | 0.125 | **0.130** | 0.004 | **8.5** |
| 0.143 | 0.156 | 0.143 | **0.147** | 0.006 | **10.5** |
| 0.162 | 0.176 | 0.162 | **0.167** | 0.007 | **12.5** |
| 0.183 | 0.197 | 0.183 | **0.188** | 0.007 | **14.5** |
| 0.205 | 0.223 | 0.204 | **0.210** | 0.009 | **16.5** |
| 0.227 | 0.247 | 0.226 | **0.233** | 0.010 | **18.5** |
| 0.252 | 0.273 | 0.250 | **0.259** | 0.010 | **20.5** |
| 0.281 | 0.297 | 0.272 | **0.283** | 0.010 | **22.5** |
| 0.308 | 0.322 | 0.295 | **0.308** | 0.011 | **24.5** |
| 0.353 | 0.348 | 0.319 | **0.340** | 0.015 | **26.5** |
| 0.402 | 0.375 | 0.345 | **0.374** | 0.023 | **28.5** |
| 0.445 | 0.405 | 0.378 | **0.409** | 0.027 | **30.5** |
| 0.489 | 0.435 | 0.412 | **0.446** | 0.032 | **32.5** |
| 0.527 | 0.464 | 0.443 | **0.478** | 0.036 | **34.5** |
| 0.561 | 0.495 | 0.474 | **0.510** | 0.037 | **36.5** |
| 0.589 | 0.523 | 0.499 | **0.537** | 0.038 | **38.5** |
| 0.609 | 0.546 | 0.522 | **0.559** | 0.037 | **40.5** |
| 0.631 | 0.560 | 0.543 | **0.578** | 0.038 | **42.5** |
| 0.647 | 0.577 | 0.563 | **0.596** | 0.037 | **44.5** |
| 0.673 | 0.590 | 0.582 | **0.615** | 0.041 | **46.5** |
| 0.688 | 0.603 | 0.598 | **0.630** | 0.042 | **48.5** |
| 0.693 | 0.610 | 0.609 | **0.637** | 0.040 | **50.5** |
| 0.703 | 0.621 | 0.621 | **0.648** | 0.039 | **52.5** |
| 0.708 | 0.630 | 0.630 | **0.656** | 0.037 | **54.5** |
| 0.718 | 0.640 | 0.641 | **0.666** | 0.036 | **56.5** |
| 0.722 | 0.648 | 0.649 | **0.673** | 0.034 | **58.5** |
| 0.728 | 0.655 | 0.658 | **0.681** | 0.034 | **60.5** |
| 0.730 | 0.661 | 0.669 | **0.687** | 0.031 | **62.5** |
| 0.738 | 0.667 | 0.673 | **0.693** | 0.032 | **64.5** |
| 0.741 | 0.673 | 0.679 | **0.698** | 0.031 | **66.5** |
| 0.745 | 0.680 | 0.687 | **0.704** | 0.029 | **68.5** |
| 0.750 | 0.682 | 0.694 | **0.709** | 0.029 | **70.5** |
| 0.758 | 0.688 | 0.702 | **0.716** | 0.030 | **72.5** |

**Supplementary Table 6B. Calculation of growth rate and doubling time at high zinc concentration.** Growth rate and doubling time were calculated as growth rate μ = (ln(xt) – ln (x0)) / (t – t0) and doubling time d= ln(2) / μ. Calculations were carried out separately for all replicates before calculating mean value (red) and standard deviation (red). Phases of exponential growth were identified using fitted trendlines and corresponding $R^2$r-values (Supplementary Figure 5B.). Values are given separately for H66 (grey) and Δ*s479* (orange) and for each of the two phases.

**Phase 1: 0.5 h-8.5 h**

| growth rate μ(H66) [$h^{-1}$] time interval 0.5 h to 8.5 h with R² = | | | | |
|---|---|---|---|---|
| **H66-1** | **H66-2** | **H66-3** | **μ(H66) [$h^{-1}$]** | **SD(μ(H66)) [$h^{-1}$]** |
| 0.121 | 0.142 | 0.155 | **0.139** | **0.014** |

| doubling time / generation time d(H66) [h] | | | | |
|---|---|---|---|---|
| **H66-1** | **H66-2** | **H66-3** | **d(H66) [h]** | **SD(d(H66)) [h]** |
| 5.7 | 4.9 | 4.5 | **5.0** | **0.5** |

| growth rate μ(s479) [$h^{-1}$] time interval 0.5 h to 8.5 h with R² = | | | | |
|---|---|---|---|---|
| **s479-1** | **s479-2** | **s479-3** | **μ(s479) [$h^{-1}$]** | **SD(μ(s479)) [$h^{-1}$]** |
| 0.084 | 0.097 | 0.089 | **0.090** | **0.005** |

| doubling time / generation time d(H66) [h] | | | | |
|---|---|---|---|---|
| **s479-1** | **s479-2** | **s479-3** | **d(s479) [h]** | **SD(d(s479)) [h]** |
| 8.2 | 7.2 | 7.8 | **7.7** | **0.4** |

**Phase2: 10.5 h-24.5 h**

| growth rate μ(H66) [$h^{-1}$] time interval 10.5 h to 24.5 h with R² = 0.9936 | | | | |
|---|---|---|---|---|
| **H66-1** | **H66-2** | **H66-3** | **μ(H66) [$h^{-1}$]** | **SD(μ(H66)) [$h^{-1}$]** |
| 0.065 | 0.062 | 0.057 | **0.062** | **0.003** |

| doubling time / generation time d(H66) [h] | | | | |
|---|---|---|---|---|
| **H66-1** | **H66-2** | **H66-3** | **d(H66) [h]** | **SD(d(H66)) [h]** |
| 10.6 | 11.1 | 12.1 | **11.3** | **0.6** |

| growth rate μ(s479) [$h^{-1}$] time interval 10.5 h to 24.5 h with R² = 0.9962 | | | | |
|---|---|---|---|---|
| **s479-1** | **s479-2** | **s479-3** | **μ(s479) [$h^{-1}$]** | **SD(μ(s479)) [$h^{-1}$]** |
| 0.055 | 0.052 | 0.051 | **0.053** | **0.001** |

| doubling time / generation time d(H66) [h] | | | | |
|---|---|---|---|---|
| **s479-1** | **s479-2** | **s479-3** | **d(s479) [h]** | **SD(d(s479)) [h]** |
| 12.7 | 13.3 | 13.5 | **13.2** | **0.3** |

**Supplementary Table 6C1. Raw growth data measured for the wildtype strain (H66) at high zinc concentration.** For each time point, $OD_{650nm}$ of three biological (replicate 1-3; H66-1 to H66-3), with three technical replicates (A-C) each, are given. Additionally, the mean $OD_{650nm}$ over all technical replicates is given for each biological replicate.

| | **replicate 1** | | | | | **replicate 2** | | | | | **replicate 3** | | | |
|---|---|---|---|---|---|---|---|---|---|---|---|---|---|---|
| | **H66-1A** | **H66-1B** | **H66-1C** | **H66-1** | | **H66-2A** | **H66-2B** | **H66-2C** | **H66-2** | | **H66-3A** | **H66-3B** | **H66-3C** | **H66-3** |
| **time [h]** | OD650 | OD650 | OD650 | **Mean OD650** | | OD650 | OD650 | OD650 | **Mean OD650** | | OD650 | OD650 | OD650 | **Mean OD650** |
| **0.5** | 0.063 | 0.066 | 0.068 | 0.066 | | 0.055 | 0.064 | 0.073 | 0.064 | | 0.059 | 0.059 | 0.067 | 0.062 |
| **1** | 0.065 | 0.069 | 0.073 | 0.069 | | 0.059 | 0.069 | 0.081 | 0.070 | | 0.065 | 0.064 | 0.073 | 0.067 |
| **1.5** | 0.07 | 0.075 | 0.078 | 0.074 | | 0.065 | 0.077 | 0.088 | 0.077 | | 0.073 | 0.071 | 0.081 | 0.075 |
| **2** | 0.077 | 0.083 | 0.086 | 0.082 | | 0.072 | 0.084 | 0.096 | 0.084 | | 0.081 | 0.079 | 0.089 | 0.083 |
| **2.5** | 0.084 | 0.091 | 0.095 | 0.090 | | 0.08 | 0.093 | 0.105 | 0.093 | | 0.091 | 0.088 | 0.099 | 0.093 |
| **3** | 0.091 | 0.099 | 0.103 | 0.098 | | 0.087 | 0.102 | 0.115 | 0.101 | | 0.1 | 0.097 | 0.108 | 0.102 |
| **3.5** | 0.098 | 0.106 | 0.111 | 0.105 | | 0.096 | 0.113 | 0.124 | 0.111 | | 0.109 | 0.106 | 0.117 | 0.111 |
| **4** | 0.105 | 0.115 | 0.12 | 0.113 | | 0.104 | 0.122 | 0.133 | 0.120 | | 0.12 | 0.115 | 0.127 | 0.121 |
| **4.5** | 0.113 | 0.124 | 0.13 | 0.122 | | 0.113 | 0.131 | 0.143 | 0.129 | | 0.13 | 0.126 | 0.136 | 0.131 |
| **5** | 0.121 | 0.133 | 0.138 | 0.131 | | 0.122 | 0.141 | 0.151 | 0.138 | | 0.142 | 0.136 | 0.145 | 0.141 |
| **5.5** | 0.129 | 0.141 | 0.147 | 0.139 | | 0.131 | 0.151 | 0.158 | 0.147 | | 0.154 | 0.146 | 0.154 | 0.151 |
| **6** | 0.135 | 0.148 | 0.153 | 0.145 | | 0.14 | 0.161 | 0.166 | 0.156 | | 0.166 | 0.156 | 0.162 | 0.161 |
| **6.5** | 0.141 | 0.155 | 0.158 | 0.151 | | 0.151 | 0.169 | 0.174 | 0.165 | | 0.177 | 0.167 | 0.169 | 0.171 |
| **7** | 0.147 | 0.161 | 0.164 | 0.157 | | 0.16 | 0.178 | 0.18 | 0.173 | | 0.19 | 0.178 | 0.177 | 0.182 |
| **7.5** | 0.153 | 0.167 | 0.17 | 0.163 | | 0.17 | 0.189 | 0.187 | 0.182 | | 0.202 | 0.189 | 0.185 | 0.192 |
| **8** | 0.159 | 0.173 | 0.174 | 0.169 | | 0.182 | 0.197 | 0.194 | 0.191 | | 0.217 | 0.202 | 0.192 | 0.204 |
| **8.5** | 0.164 | 0.176 | 0.177 | 0.172 | | 0.193 | 0.206 | 0.2 | 0.200 | | 0.23 | 0.213 | 0.197 | 0.213 |
| **9** | 0.171 | 0.183 | 0.183 | 0.179 | | 0.205 | 0.216 | 0.208 | 0.210 | | 0.243 | 0.224 | 0.204 | 0.224 |
| **9.5** | 0.174 | 0.191 | 0.188 | 0.184 | | 0.213 | 0.226 | 0.213 | 0.217 | | 0.256 | 0.236 | 0.21 | 0.234 |
| **10** | 0.182 | 0.196 | 0.193 | 0.190 | | 0.225 | 0.237 | 0.221 | 0.228 | | 0.269 | 0.25 | 0.218 | 0.246 |
| **10.5** | 0.188 | 0.203 | 0.198 | 0.196 | | 0.239 | 0.249 | 0.229 | 0.239 | | 0.283 | 0.263 | 0.226 | 0.257 |
| **11** | 0.192 | 0.208 | 0.205 | 0.202 | | 0.249 | 0.26 | 0.239 | 0.249 | | 0.294 | 0.276 | 0.229 | 0.266 |
| **11.5** | 0.198 | 0.214 | 0.21 | 0.207 | | 0.26 | 0.273 | 0.248 | 0.260 | | 0.305 | 0.288 | 0.238 | 0.277 |
| **12** | 0.206 | 0.221 | 0.217 | 0.215 | | 0.273 | 0.285 | 0.258 | 0.272 | | 0.317 | 0.302 | 0.245 | 0.288 |
| **12.5** | 0.21 | 0.228 | 0.222 | 0.220 | | 0.285 | 0.301 | 0.269 | 0.285 | | 0.327 | 0.317 | 0.254 | 0.299 |
| **13** | 0.215 | 0.235 | 0.228 | 0.226 | | 0.297 | 0.314 | 0.279 | 0.297 | | 0.337 | 0.33 | 0.262 | 0.310 |
| **13.5** | 0.222 | 0.241 | 0.233 | 0.232 | | 0.311 | 0.328 | 0.292 | 0.310 | | 0.35 | 0.343 | 0.273 | 0.322 |
| **14** | 0.227 | 0.249 | 0.24 | 0.239 | | 0.325 | 0.342 | 0.307 | 0.325 | | 0.362 | 0.358 | 0.285 | 0.335 |
| **14.5** | 0.235 | 0.255 | 0.245 | 0.245 | | 0.339 | 0.354 | 0.324 | 0.339 | | 0.372 | 0.37 | 0.299 | 0.347 |
| **15** | 0.243 | 0.264 | 0.254 | 0.254 | | 0.349 | 0.368 | 0.338 | 0.352 | | 0.38 | 0.383 | 0.313 | 0.359 |
| **15.5** | 0.25 | 0.271 | 0.259 | 0.260 | | 0.362 | 0.381 | 0.354 | 0.366 | | 0.388 | 0.395 | 0.328 | 0.370 |
| **16** | 0.259 | 0.279 | 0.265 | 0.268 | | 0.372 | 0.394 | 0.37 | 0.379 | | 0.398 | 0.408 | 0.343 | 0.383 |
| **16.5** | 0.266 | 0.285 | 0.27 | 0.274 | | 0.385 | 0.406 | 0.387 | 0.393 | | 0.405 | 0.418 | 0.36 | 0.394 |
| **17** | 0.274 | 0.294 | 0.277 | 0.282 | | 0.396 | 0.42 | 0.406 | 0.407 | | 0.417 | 0.432 | 0.378 | 0.409 |
| **17.5** | 0.282 | 0.303 | 0.284 | 0.290 | | 0.406 | 0.433 | 0.423 | 0.421 | | 0.427 | 0.443 | 0.393 | 0.421 |
| **18** | 0.29 | 0.313 | 0.291 | 0.298 | | 0.421 | 0.446 | 0.438 | 0.435 | | 0.435 | 0.456 | 0.406 | 0.432 |
| **18.5** | 0.3 | 0.325 | 0.3 | 0.308 | | 0.426 | 0.456 | 0.451 | 0.444 | | 0.445 | 0.468 | 0.428 | 0.447 |
| **19** | 0.311 | 0.336 | 0.31 | 0.319 | | 0.443 | 0.47 | 0.469 | 0.461 | | 0.455 | 0.48 | 0.441 | 0.459 |
| **19.5** | 0.322 | 0.349 | 0.321 | 0.331 | | 0.45 | 0.479 | 0.482 | 0.470 | | 0.466 | 0.492 | 0.459 | 0.472 |
| **20** | 0.334 | 0.364 | 0.333 | 0.344 | | 0.465 | 0.493 | 0.495 | 0.484 | | 0.474 | 0.504 | 0.473 | 0.484 |

| | | | | | | | | | | | | | | |
|---|---|---|---|---|---|---|---|---|---|---|---|---|---|---|
| **20.5** | 0.347 | 0.377 | 0.347 | 0.357 | | 0.477 | 0.502 | 0.509 | 0.496 | | 0.485 | 0.514 | 0.487 | 0.495 |
| **21** | 0.362 | 0.395 | 0.359 | 0.372 | | 0.486 | 0.514 | 0.522 | 0.507 | | 0.492 | 0.524 | 0.502 | 0.506 |
| **21.5** | 0.377 | 0.413 | 0.375 | 0.388 | | 0.498 | 0.525 | 0.532 | 0.518 | | 0.502 | 0.535 | 0.514 | 0.517 |
| **22** | 0.394 | 0.43 | 0.392 | 0.405 | | 0.508 | 0.531 | 0.544 | 0.528 | | 0.512 | 0.545 | 0.526 | 0.528 |
| **22.5** | 0.41 | 0.448 | 0.41 | 0.423 | | 0.514 | 0.544 | 0.555 | 0.538 | | 0.52 | 0.554 | 0.537 | 0.537 |
| **23** | 0.428 | 0.465 | 0.426 | 0.440 | | 0.526 | 0.552 | 0.565 | 0.548 | | 0.53 | 0.565 | 0.547 | 0.547 |
| **23.5** | 0.446 | 0.481 | 0.444 | 0.457 | | 0.535 | 0.561 | 0.575 | 0.557 | | 0.538 | 0.573 | 0.559 | 0.557 |
| **24** | 0.462 | 0.496 | 0.461 | 0.473 | | 0.544 | 0.568 | 0.585 | 0.566 | | 0.545 | 0.581 | 0.566 | 0.564 |
| **24.5** | 0.476 | 0.513 | 0.479 | 0.489 | | 0.55 | 0.576 | 0.592 | 0.573 | | 0.553 | 0.589 | 0.577 | 0.573 |
| **25** | 0.492 | 0.529 | 0.494 | 0.505 | | 0.558 | 0.584 | 0.601 | 0.581 | | 0.563 | 0.597 | 0.586 | 0.582 |
| **25.5** | 0.506 | 0.543 | 0.514 | 0.521 | | 0.566 | 0.592 | 0.608 | 0.589 | | 0.569 | 0.605 | 0.593 | 0.589 |
| **26** | 0.519 | 0.557 | 0.531 | 0.536 | | 0.573 | 0.598 | 0.616 | 0.596 | | 0.575 | 0.61 | 0.601 | 0.595 |
| **26.5** | 0.535 | 0.57 | 0.546 | 0.550 | | 0.578 | 0.603 | 0.623 | 0.601 | | 0.584 | 0.618 | 0.61 | 0.604 |
| **27** | 0.546 | 0.583 | 0.561 | 0.563 | | 0.586 | 0.611 | 0.63 | 0.609 | | 0.589 | 0.625 | 0.616 | 0.610 |
| **27.5** | 0.559 | 0.595 | 0.576 | 0.577 | | 0.59 | 0.617 | 0.636 | 0.614 | | 0.596 | 0.631 | 0.622 | 0.616 |
| **28** | 0.568 | 0.605 | 0.59 | 0.588 | | 0.596 | 0.623 | 0.642 | 0.620 | | 0.603 | 0.636 | 0.629 | 0.623 |
| **28.5** | 0.576 | 0.616 | 0.604 | 0.599 | | 0.601 | 0.629 | 0.647 | 0.626 | | 0.61 | 0.641 | 0.637 | 0.629 |
| **29** | 0.587 | 0.626 | 0.616 | 0.610 | | 0.607 | 0.634 | 0.654 | 0.632 | | 0.616 | 0.646 | 0.642 | 0.635 |
| **29.5** | 0.595 | 0.634 | 0.628 | 0.619 | | 0.613 | 0.64 | 0.66 | 0.638 | | 0.621 | 0.652 | 0.648 | 0.640 |
| **30** | 0.602 | 0.643 | 0.636 | 0.627 | | 0.617 | 0.645 | 0.665 | 0.642 | | 0.626 | 0.656 | 0.652 | 0.645 |
| **30.5** | 0.61 | 0.651 | 0.649 | 0.637 | | 0.624 | 0.651 | 0.67 | 0.648 | | 0.633 | 0.662 | 0.659 | 0.651 |
| **31** | 0.618 | 0.661 | 0.66 | 0.646 | | 0.63 | 0.655 | 0.675 | 0.653 | | 0.639 | 0.667 | 0.664 | 0.657 |
| **31.5** | 0.625 | 0.667 | 0.67 | 0.654 | | 0.635 | 0.66 | 0.679 | 0.658 | | 0.645 | 0.672 | 0.666 | 0.661 |
| **32** | 0.631 | 0.675 | 0.679 | 0.662 | | 0.64 | 0.665 | 0.683 | 0.663 | | 0.651 | 0.677 | 0.673 | 0.667 |
| **32.5** | 0.638 | 0.682 | 0.69 | 0.670 | | 0.647 | 0.67 | 0.688 | 0.668 | | 0.658 | 0.681 | 0.677 | 0.672 |
| **33** | 0.644 | 0.689 | 0.698 | 0.677 | | 0.651 | 0.674 | 0.691 | 0.672 | | 0.662 | 0.687 | 0.682 | 0.677 |
| **33.5** | 0.65 | 0.695 | 0.706 | 0.684 | | 0.656 | 0.679 | 0.696 | 0.677 | | 0.666 | 0.692 | 0.685 | 0.681 |
| **34** | 0.655 | 0.701 | 0.712 | 0.689 | | 0.661 | 0.684 | 0.7 | 0.682 | | 0.673 | 0.697 | 0.689 | 0.686 |
| **34.5** | 0.662 | 0.708 | 0.722 | 0.697 | | 0.666 | 0.689 | 0.704 | 0.686 | | 0.678 | 0.701 | 0.694 | 0.691 |
| **35** | 0.667 | 0.713 | 0.731 | 0.704 | | 0.671 | 0.693 | 0.708 | 0.691 | | 0.684 | 0.706 | 0.699 | 0.696 |
| **35.5** | 0.673 | 0.719 | 0.737 | 0.710 | | 0.677 | 0.697 | 0.712 | 0.695 | | 0.688 | 0.71 | 0.703 | 0.700 |
| **36** | 0.679 | 0.725 | 0.743 | 0.716 | | 0.681 | 0.702 | 0.716 | 0.700 | | 0.694 | 0.716 | 0.707 | 0.706 |
| **36.5** | 0.684 | 0.73 | 0.75 | 0.721 | | 0.687 | 0.707 | 0.719 | 0.704 | | 0.7 | 0.719 | 0.711 | 0.710 |
| **37** | 0.689 | 0.735 | 0.755 | 0.726 | | 0.692 | 0.71 | 0.72 | 0.707 | | 0.706 | 0.725 | 0.716 | 0.716 |
| **37.5** | 0.694 | 0.742 | 0.761 | 0.732 | | 0.695 | 0.713 | 0.724 | 0.711 | | 0.71 | 0.731 | 0.719 | 0.720 |
| **38** | 0.7 | 0.746 | 0.766 | 0.737 | | 0.699 | 0.716 | 0.725 | 0.713 | | 0.714 | 0.735 | 0.722 | 0.724 |
| **38.5** | 0.705 | 0.754 | 0.767 | 0.742 | | 0.704 | 0.719 | 0.729 | 0.717 | | 0.719 | 0.739 | 0.725 | 0.728 |
| **39** | 0.708 | 0.756 | 0.774 | 0.746 | | 0.706 | 0.721 | 0.731 | 0.719 | | 0.722 | 0.743 | 0.727 | 0.731 |
| **39.5** | 0.712 | 0.761 | 0.778 | 0.750 | | 0.707 | 0.726 | 0.734 | 0.722 | | 0.725 | 0.747 | 0.73 | 0.734 |
| **40** | 0.719 | 0.767 | 0.781 | 0.756 | | 0.712 | 0.729 | 0.738 | 0.726 | | 0.728 | 0.752 | 0.733 | 0.738 |
| **40.5** | 0.724 | 0.771 | 0.786 | 0.760 | | 0.717 | 0.732 | 0.741 | 0.730 | | 0.731 | 0.754 | 0.737 | 0.741 |
| **41** | 0.729 | 0.776 | 0.788 | 0.764 | | 0.72 | 0.736 | 0.745 | 0.734 | | 0.733 | 0.758 | 0.738 | 0.743 |
| **41.5** | 0.733 | 0.78 | 0.792 | 0.768 | | 0.723 | 0.737 | 0.747 | 0.736 | | 0.737 | 0.761 | 0.742 | 0.747 |
| **42** | 0.736 | 0.783 | 0.795 | 0.771 | | 0.725 | 0.741 | 0.749 | 0.738 | | 0.74 | 0.763 | 0.743 | 0.749 |
| **42.5** | 0.739 | 0.788 | 0.799 | 0.775 | | 0.727 | 0.744 | 0.752 | 0.741 | | 0.744 | 0.768 | 0.75 | 0.754 |
| **43** | 0.743 | 0.79 | 0.8 | 0.778 | | 0.73 | 0.746 | 0.752 | 0.743 | | 0.748 | 0.771 | 0.749 | 0.756 |
| **43.5** | 0.745 | 0.792 | 0.802 | 0.780 | | 0.732 | 0.748 | 0.751 | 0.744 | | 0.75 | 0.774 | 0.753 | 0.759 |
| **44** | 0.747 | 0.794 | 0.803 | 0.781 | | 0.735 | 0.75 | 0.754 | 0.746 | | 0.753 | 0.778 | 0.754 | 0.762 |
| **44.5** | 0.751 | 0.796 | 0.806 | 0.784 | | 0.737 | 0.75 | 0.752 | 0.746 | | 0.756 | 0.779 | 0.754 | 0.763 |
| **45** | 0.756 | 0.799 | 0.806 | 0.787 | | 0.74 | 0.751 | 0.756 | 0.749 | | 0.758 | 0.782 | 0.753 | 0.764 |

| | | | | | | | | | | | | | | |
|---|---|---|---|---|---|---|---|---|---|---|---|---|---|---|
| **45.5** | 0.757 | 0.797 | 0.804 | 0.786 | | 0.741 | 0.752 | 0.757 | 0.750 | | 0.76 | 0.783 | 0.754 | 0.766 |
| **46** | 0.76 | 0.797 | 0.807 | 0.788 | | 0.743 | 0.754 | 0.757 | 0.751 | | 0.763 | 0.786 | 0.758 | 0.769 |
| **46.5** | 0.765 | 0.8 | 0.807 | 0.791 | | 0.744 | 0.755 | 0.759 | 0.753 | | 0.765 | 0.788 | 0.759 | 0.771 |
| **47** | 0.764 | 0.8 | 0.807 | 0.790 | | 0.744 | 0.756 | 0.76 | 0.753 | | 0.764 | 0.789 | 0.758 | 0.770 |
| **47.5** | 0.77 | 0.803 | 0.81 | 0.794 | | 0.747 | 0.757 | 0.761 | 0.755 | | 0.767 | 0.79 | 0.761 | 0.773 |
| **48** | 0.77 | 0.803 | 0.809 | 0.794 | | 0.748 | 0.759 | 0.762 | 0.756 | | 0.768 | 0.792 | 0.762 | 0.774 |
| **48.5** | 0.775 | 0.806 | 0.813 | 0.798 | | 0.749 | 0.76 | 0.764 | 0.758 | | 0.769 | 0.792 | 0.763 | 0.775 |
| **49** | 0.773 | 0.804 | 0.813 | 0.797 | | 0.75 | 0.76 | 0.763 | 0.758 | | 0.772 | 0.793 | 0.762 | 0.776 |
| **49.5** | 0.773 | 0.807 | 0.813 | 0.798 | | 0.752 | 0.762 | 0.765 | 0.760 | | 0.773 | 0.794 | 0.767 | 0.778 |
| **50** | 0.775 | 0.81 | 0.814 | 0.800 | | 0.752 | 0.764 | 0.767 | 0.761 | | 0.774 | 0.795 | 0.765 | 0.778 |
| **50.5** | 0.778 | 0.811 | 0.816 | 0.802 | | 0.753 | 0.766 | 0.77 | 0.763 | | 0.777 | 0.798 | 0.769 | 0.781 |
| **51** | 0.78 | 0.812 | 0.817 | 0.803 | | 0.751 | 0.767 | 0.768 | 0.762 | | 0.776 | 0.797 | 0.773 | 0.782 |
| **51.5** | 0.785 | 0.815 | 0.814 | 0.805 | | 0.754 | 0.769 | 0.771 | 0.765 | | 0.779 | 0.8 | 0.775 | 0.785 |
| **52** | 0.783 | 0.813 | 0.814 | 0.803 | | 0.754 | 0.77 | 0.773 | 0.766 | | 0.78 | 0.801 | 0.777 | 0.786 |
| **52.5** | 0.786 | 0.814 | 0.817 | 0.806 | | 0.759 | 0.771 | 0.775 | 0.768 | | 0.782 | 0.803 | 0.778 | 0.788 |
| **53** | 0.787 | 0.816 | 0.821 | 0.808 | | 0.762 | 0.772 | 0.776 | 0.770 | | 0.783 | 0.804 | 0.781 | 0.789 |
| **53.5** | 0.787 | 0.815 | 0.82 | 0.807 | | 0.762 | 0.774 | 0.779 | 0.772 | | 0.785 | 0.806 | 0.781 | 0.791 |
| **54** | 0.788 | 0.815 | 0.82 | 0.808 | | 0.763 | 0.775 | 0.779 | 0.772 | | 0.785 | 0.807 | 0.78 | 0.791 |
| **54.5** | 0.789 | 0.816 | 0.82 | 0.808 | | 0.766 | 0.777 | 0.782 | 0.775 | | 0.788 | 0.808 | 0.787 | 0.794 |
| **55** | 0.786 | 0.816 | 0.824 | 0.809 | | 0.766 | 0.778 | 0.782 | 0.775 | | 0.789 | 0.809 | 0.783 | 0.794 |
| **55.5** | 0.79 | 0.817 | 0.824 | 0.810 | | 0.77 | 0.779 | 0.785 | 0.778 | | 0.79 | 0.812 | 0.79 | 0.797 |
| **56** | 0.793 | 0.819 | 0.828 | 0.813 | | 0.771 | 0.781 | 0.784 | 0.779 | | 0.792 | 0.814 | 0.788 | 0.798 |
| **56.5** | 0.794 | 0.82 | 0.827 | 0.814 | | 0.773 | 0.783 | 0.788 | 0.781 | | 0.793 | 0.816 | 0.792 | 0.800 |
| **57** | 0.793 | 0.821 | 0.83 | 0.815 | | 0.774 | 0.784 | 0.788 | 0.782 | | 0.797 | 0.817 | 0.788 | 0.801 |
| **57.5** | 0.795 | 0.823 | 0.829 | 0.816 | | 0.775 | 0.785 | 0.79 | 0.783 | | 0.797 | 0.816 | 0.793 | 0.802 |
| **58** | 0.795 | 0.821 | 0.83 | 0.815 | | 0.776 | 0.787 | 0.791 | 0.785 | | 0.799 | 0.819 | 0.795 | 0.804 |
| **58.5** | 0.794 | 0.82 | 0.83 | 0.815 | | 0.777 | 0.788 | 0.793 | 0.786 | | 0.801 | 0.82 | 0.795 | 0.805 |
| **59** | 0.794 | 0.82 | 0.833 | 0.816 | | 0.779 | 0.79 | 0.795 | 0.788 | | 0.803 | 0.821 | 0.796 | 0.807 |
| **59.5** | 0.795 | 0.821 | 0.835 | 0.817 | | 0.78 | 0.791 | 0.795 | 0.789 | | 0.803 | 0.823 | 0.796 | 0.807 |
| **60** | 0.798 | 0.822 | 0.835 | 0.818 | | 0.782 | 0.793 | 0.796 | 0.790 | | 0.806 | 0.825 | 0.801 | 0.811 |
| **60.5** | 0.797 | 0.825 | 0.837 | 0.820 | | 0.784 | 0.793 | 0.798 | 0.792 | | 0.807 | 0.828 | 0.802 | 0.812 |
| **61** | 0.8 | 0.824 | 0.837 | 0.820 | | 0.785 | 0.793 | 0.798 | 0.792 | | 0.808 | 0.826 | 0.802 | 0.812 |
| **61.5** | 0.799 | 0.825 | 0.838 | 0.821 | | 0.786 | 0.795 | 0.8 | 0.794 | | 0.81 | 0.828 | 0.803 | 0.814 |
| **62** | 0.803 | 0.827 | 0.84 | 0.823 | | 0.787 | 0.797 | 0.803 | 0.796 | | 0.812 | 0.829 | 0.807 | 0.816 |
| **62.5** | 0.801 | 0.828 | 0.84 | 0.823 | | 0.788 | 0.797 | 0.803 | 0.796 | | 0.813 | 0.831 | 0.807 | 0.817 |
| **63** | 0.805 | 0.831 | 0.841 | 0.826 | | 0.791 | 0.799 | 0.804 | 0.798 | | 0.815 | 0.832 | 0.805 | 0.817 |
| **63.5** | 0.804 | 0.83 | 0.843 | 0.826 | | 0.792 | 0.8 | 0.807 | 0.800 | | 0.816 | 0.834 | 0.809 | 0.820 |
| **64** | 0.803 | 0.831 | 0.843 | 0.826 | | 0.793 | 0.801 | 0.808 | 0.801 | | 0.818 | 0.836 | 0.81 | 0.821 |
| **64.5** | 0.804 | 0.832 | 0.844 | 0.827 | | 0.795 | 0.803 | 0.809 | 0.802 | | 0.819 | 0.836 | 0.811 | 0.822 |
| **65** | 0.806 | 0.833 | 0.844 | 0.828 | | 0.794 | 0.805 | 0.811 | 0.803 | | 0.819 | 0.836 | 0.814 | 0.823 |
| **65.5** | 0.806 | 0.833 | 0.848 | 0.829 | | 0.796 | 0.805 | 0.812 | 0.804 | | 0.823 | 0.838 | 0.814 | 0.825 |
| **66** | 0.803 | 0.833 | 0.848 | 0.828 | | 0.799 | 0.807 | 0.812 | 0.806 | | 0.823 | 0.84 | 0.816 | 0.826 |
| **66.5** | 0.806 | 0.837 | 0.85 | 0.831 | | 0.799 | 0.808 | 0.813 | 0.807 | | 0.825 | 0.841 | 0.816 | 0.827 |
| **67** | 0.808 | 0.835 | 0.848 | 0.830 | | 0.8 | 0.809 | 0.815 | 0.808 | | 0.826 | 0.843 | 0.819 | 0.829 |
| **67.5** | 0.806 | 0.835 | 0.849 | 0.830 | | 0.801 | 0.81 | 0.817 | 0.809 | | 0.828 | 0.842 | 0.82 | 0.830 |
| **68** | 0.815 | 0.84 | 0.853 | 0.836 | | 0.804 | 0.812 | 0.819 | 0.812 | | 0.83 | 0.844 | 0.823 | 0.832 |
| **68.5** | 0.81 | 0.838 | 0.855 | 0.834 | | 0.804 | 0.813 | 0.82 | 0.812 | | 0.83 | 0.845 | 0.822 | 0.832 |
| **69** | 0.815 | 0.841 | 0.855 | 0.837 | | 0.806 | 0.814 | 0.821 | 0.814 | | 0.834 | 0.847 | 0.826 | 0.836 |
| **69.5** | 0.816 | 0.837 | 0.854 | 0.836 | | 0.805 | 0.816 | 0.822 | 0.814 | | 0.833 | 0.847 | 0.824 | 0.835 |
| **70** | 0.814 | 0.841 | 0.856 | 0.837 | | 0.807 | 0.815 | 0.824 | 0.815 | | 0.834 | 0.848 | 0.828 | 0.837 |
| **70.5** | 0.818 | 0.841 | 0.858 | 0.839 | | 0.808 | 0.818 | 0.826 | 0.817 | | 0.835 | 0.849 | 0.826 | 0.837 |
| **71** | 0.82 | 0.843 | 0.859 | 0.841 | | 0.809 | 0.819 | 0.826 | 0.818 | | 0.838 | 0.849 | 0.83 | 0.839 |

| | | | | | | | | | | | | | | |
|---|---|---|---|---|---|---|---|---|---|---|---|---|---|---|
| **71.5** | 0.818 | 0.839 | 0.86 | 0.839 | | 0.81 | 0.818 | 0.826 | 0.818 | | 0.838 | 0.849 | 0.829 | 0.839 |
| **72** | 0.818 | 0.839 | 0.86 | 0.839 | | 0.812 | 0.821 | 0.829 | 0.821 | | 0.84 | 0.851 | 0.831 | 0.841 |
| **72.5** | 0.82 | 0.841 | 0.863 | 0.841 | | 0.813 | 0.821 | 0.829 | 0.821 | | 0.841 | 0.851 | 0.833 | 0.842 |

**Supplementary Table 6C2. Raw growth data measured for the deletion strain (Δ*s479*) at high zinc concentration.** For each time point, $OD_{650nm}$ of three biological (replicate 1-3; s479-1 to s479-3), with three technical replicates (A to C) each, are given. Additionally, the mean $OD_{650nm}$ over all technical replicates is given for each biological replicate.

| | **replicate 1** | | | | | **replicate 2** | | | | | **replicate 3** | | | |
|---|---|---|---|---|---|---|---|---|---|---|---|---|---|---|
| | s479-1A | s479-1B | **s479-1C** | **s479-1** | | **s479-2A** | **s479-2B** | **s479-2C** | **s479-2** | | **s479-3A** | **s479-3B** | **s479-3C** | **s479-3** |
| **time [h]** | OD650 | OD650 | OD650 | **Mean OD650** | | OD650 | OD650 | OD650 | **Mean OD650** | | OD650 | OD650 | OD650 | **Mean OD650** |
| **0.5** | 0.058 | 0.071 | 0.068 | 0.066 | | 0.066 | 0.062 | 0.06 | 0.063 | | 0.061 | 0.06 | 0.063 | 0.061 |
| **1** | 0.062 | 0.07 | 0.067 | 0.066 | | 0.069 | 0.064 | 0.063 | 0.065 | | 0.064 | 0.063 | 0.066 | 0.064 |
| **1.5** | 0.064 | 0.076 | 0.072 | 0.071 | | 0.072 | 0.068 | 0.066 | 0.069 | | 0.064 | 0.065 | 0.069 | 0.066 |
| **2** | 0.067 | 0.081 | 0.076 | 0.075 | | 0.075 | 0.071 | 0.071 | 0.072 | | 0.068 | 0.068 | 0.075 | 0.070 |
| **2.5** | 0.07 | 0.084 | 0.079 | 0.078 | | 0.077 | 0.074 | 0.074 | 0.075 | | 0.072 | 0.071 | 0.081 | 0.075 |
| **3** | 0.074 | 0.086 | 0.083 | 0.081 | | 0.082 | 0.078 | 0.079 | 0.080 | | 0.073 | 0.075 | 0.087 | 0.078 |
| **3.5** | 0.079 | 0.091 | 0.087 | 0.086 | | 0.085 | 0.082 | 0.084 | 0.084 | | 0.077 | 0.079 | 0.093 | 0.083 |
| **4** | 0.083 | 0.097 | 0.092 | 0.091 | | 0.091 | 0.087 | 0.09 | 0.089 | | 0.08 | 0.082 | 0.097 | 0.086 |
| **4.5** | 0.087 | 0.101 | 0.097 | 0.095 | | 0.096 | 0.091 | 0.094 | 0.094 | | 0.083 | 0.087 | 0.102 | 0.091 |
| **5** | 0.092 | 0.106 | 0.102 | 0.100 | | 0.101 | 0.098 | 0.1 | 0.100 | | 0.086 | 0.091 | 0.108 | 0.095 |
| **5.5** | 0.097 | 0.11 | 0.108 | 0.105 | | 0.105 | 0.102 | 0.105 | 0.104 | | 0.089 | 0.096 | 0.113 | 0.099 |
| **6** | 0.1 | 0.115 | 0.111 | 0.109 | | 0.113 | 0.108 | 0.112 | 0.111 | | 0.092 | 0.1 | 0.118 | 0.103 |
| **6.5** | 0.103 | 0.118 | 0.114 | 0.112 | | 0.12 | 0.111 | 0.116 | 0.116 | | 0.095 | 0.105 | 0.122 | 0.107 |
| **7** | 0.107 | 0.123 | 0.119 | 0.116 | | 0.121 | 0.117 | 0.121 | 0.120 | | 0.098 | 0.11 | 0.126 | 0.111 |
| **7.5** | 0.111 | 0.126 | 0.123 | 0.120 | | 0.13 | 0.121 | 0.128 | 0.126 | | 0.101 | 0.115 | 0.131 | 0.116 |
| **8** | 0.118 | 0.129 | 0.126 | 0.124 | | 0.133 | 0.125 | 0.132 | 0.130 | | 0.103 | 0.121 | 0.136 | 0.120 |
| **8.5** | 0.122 | 0.134 | 0.13 | 0.129 | | 0.141 | 0.13 | 0.137 | 0.136 | | 0.107 | 0.127 | 0.142 | 0.125 |
| **9** | 0.124 | 0.138 | 0.134 | 0.132 | | 0.146 | 0.137 | 0.144 | 0.142 | | 0.11 | 0.133 | 0.147 | 0.130 |
| **9.5** | 0.127 | 0.144 | 0.136 | 0.136 | | 0.149 | 0.139 | 0.147 | 0.145 | | 0.113 | 0.136 | 0.151 | 0.133 |
| **10** | 0.131 | 0.146 | 0.142 | 0.140 | | 0.156 | 0.143 | 0.153 | 0.151 | | 0.117 | 0.143 | 0.156 | 0.139 |
| **10.5** | 0.134 | 0.149 | 0.146 | 0.143 | | 0.161 | 0.148 | 0.158 | 0.156 | | 0.12 | 0.149 | 0.161 | 0.143 |
| **11** | 0.138 | 0.154 | 0.151 | 0.148 | | 0.166 | 0.152 | 0.164 | 0.161 | | 0.123 | 0.154 | 0.167 | 0.148 |
| **11.5** | 0.141 | 0.16 | 0.156 | 0.152 | | 0.172 | 0.157 | 0.169 | 0.166 | | 0.127 | 0.16 | 0.171 | 0.153 |
| **12** | 0.146 | 0.164 | 0.162 | 0.157 | | 0.175 | 0.161 | 0.174 | 0.170 | | 0.13 | 0.166 | 0.176 | 0.157 |
| **12.5** | 0.149 | 0.169 | 0.167 | 0.162 | | 0.181 | 0.167 | 0.18 | 0.176 | | 0.134 | 0.172 | 0.181 | 0.162 |
| **13** | 0.152 | 0.174 | 0.171 | 0.166 | | 0.185 | 0.17 | 0.183 | 0.179 | | 0.137 | 0.178 | 0.186 | 0.167 |
| **13.5** | 0.155 | 0.18 | 0.176 | 0.170 | | 0.192 | 0.178 | 0.19 | 0.187 | | 0.142 | 0.186 | 0.191 | 0.173 |
| **14** | 0.161 | 0.187 | 0.182 | 0.177 | | 0.197 | 0.184 | 0.195 | 0.192 | | 0.147 | 0.192 | 0.197 | 0.179 |
| **14.5** | 0.167 | 0.193 | 0.189 | 0.183 | | 0.203 | 0.188 | 0.201 | 0.197 | | 0.149 | 0.197 | 0.202 | 0.183 |
| **15** | 0.171 | 0.199 | 0.193 | 0.188 | | 0.209 | 0.194 | 0.208 | 0.204 | | 0.154 | 0.203 | 0.205 | 0.187 |
| **15.5** | 0.176 | 0.204 | 0.197 | 0.192 | | 0.216 | 0.201 | 0.212 | 0.210 | | 0.156 | 0.211 | 0.211 | 0.193 |
| **16** | 0.181 | 0.21 | 0.205 | 0.199 | | 0.222 | 0.206 | 0.218 | 0.215 | | 0.161 | 0.216 | 0.216 | 0.198 |
| **16.5** | 0.186 | 0.215 | 0.213 | 0.205 | | 0.231 | 0.212 | 0.225 | 0.223 | | 0.169 | 0.221 | 0.221 | 0.204 |
| **17** | 0.193 | 0.221 | 0.22 | 0.211 | | 0.237 | 0.219 | 0.231 | 0.229 | | 0.175 | 0.228 | 0.227 | 0.210 |
| **17.5** | 0.199 | 0.226 | 0.225 | 0.217 | | 0.241 | 0.223 | 0.238 | 0.234 | | 0.176 | 0.233 | 0.231 | 0.213 |
| **18** | 0.204 | 0.23 | 0.231 | 0.222 | | 0.251 | 0.229 | 0.243 | 0.241 | | 0.186 | 0.239 | 0.236 | 0.220 |
| **18.5** | 0.208 | 0.235 | 0.237 | 0.227 | | 0.256 | 0.235 | 0.25 | 0.247 | | 0.192 | 0.245 | 0.241 | 0.226 |
| **19** | 0.219 | 0.24 | 0.24 | 0.233 | | 0.262 | 0.241 | 0.256 | 0.253 | | 0.198 | 0.251 | 0.246 | 0.232 |
| **19.5** | 0.226 | 0.246 | 0.246 | 0.239 | | 0.267 | 0.248 | 0.261 | 0.259 | | 0.204 | 0.257 | 0.251 | 0.237 |
| **20** | 0.233 | 0.251 | 0.249 | 0.244 | | 0.276 | 0.255 | 0.268 | 0.266 | | 0.211 | 0.264 | 0.257 | 0.244 |
| **20.5** | 0.247 | 0.256 | 0.254 | 0.252 | | 0.283 | 0.262 | 0.275 | 0.273 | | 0.218 | 0.27 | 0.262 | 0.250 |

| | | | | | | | | | | | | | | |
|---|---|---|---|---|---|---|---|---|---|---|---|---|---|---|
| **21** | 0.257 | 0.264 | 0.261 | 0.261 | | 0.288 | 0.267 | 0.281 | 0.279 | | 0.224 | 0.276 | 0.267 | 0.256 |
| **21.5** | 0.265 | 0.272 | 0.267 | 0.268 | | 0.294 | 0.271 | 0.286 | 0.284 | | 0.228 | 0.282 | 0.273 | 0.261 |
| **22** | 0.275 | 0.278 | 0.271 | 0.275 | | 0.301 | 0.277 | 0.294 | 0.291 | | 0.234 | 0.289 | 0.278 | 0.267 |
| **22.5** | 0.278 | 0.287 | 0.277 | 0.281 | | 0.307 | 0.284 | 0.299 | 0.297 | | 0.239 | 0.295 | 0.283 | 0.272 |
| **23** | 0.284 | 0.296 | 0.283 | 0.288 | | 0.314 | 0.29 | 0.306 | 0.303 | | 0.244 | 0.302 | 0.288 | 0.278 |
| **23.5** | 0.286 | 0.305 | 0.29 | 0.294 | | 0.32 | 0.296 | 0.313 | 0.310 | | 0.248 | 0.308 | 0.294 | 0.283 |
| **24** | 0.289 | 0.314 | 0.299 | 0.301 | | 0.326 | 0.301 | 0.318 | 0.315 | | 0.255 | 0.314 | 0.3 | 0.290 |
| **24.5** | 0.292 | 0.323 | 0.308 | 0.308 | | 0.334 | 0.307 | 0.326 | 0.322 | | 0.26 | 0.319 | 0.305 | 0.295 |
| **25** | 0.298 | 0.335 | 0.319 | 0.317 | | 0.339 | 0.313 | 0.332 | 0.328 | | 0.266 | 0.326 | 0.31 | 0.301 |
| **25.5** | 0.306 | 0.346 | 0.329 | 0.327 | | 0.346 | 0.319 | 0.338 | 0.334 | | 0.271 | 0.332 | 0.315 | 0.306 |
| **26** | 0.319 | 0.357 | 0.342 | 0.339 | | 0.354 | 0.325 | 0.345 | 0.341 | | 0.278 | 0.34 | 0.319 | 0.312 |
| **26.5** | 0.337 | 0.367 | 0.355 | 0.353 | | 0.362 | 0.332 | 0.351 | 0.348 | | 0.284 | 0.346 | 0.327 | 0.319 |
| **27** | 0.352 | 0.379 | 0.367 | 0.366 | | 0.37 | 0.337 | 0.357 | 0.355 | | 0.289 | 0.353 | 0.331 | 0.324 |
| **27.5** | 0.359 | 0.39 | 0.381 | 0.377 | | 0.378 | 0.343 | 0.363 | 0.361 | | 0.296 | 0.36 | 0.338 | 0.331 |
| **28** | 0.369 | 0.405 | 0.395 | 0.390 | | 0.387 | 0.351 | 0.37 | 0.369 | | 0.303 | 0.368 | 0.343 | 0.338 |
| **28.5** | 0.38 | 0.417 | 0.409 | 0.402 | | 0.394 | 0.356 | 0.375 | 0.375 | | 0.311 | 0.375 | 0.35 | 0.345 |
| **29** | 0.389 | 0.429 | 0.42 | 0.413 | | 0.404 | 0.362 | 0.383 | 0.383 | | 0.32 | 0.385 | 0.357 | 0.354 |
| **29.5** | 0.399 | 0.442 | 0.433 | 0.425 | | 0.414 | 0.37 | 0.387 | 0.390 | | 0.331 | 0.392 | 0.363 | 0.362 |
| **30** | 0.408 | 0.455 | 0.444 | 0.436 | | 0.422 | 0.376 | 0.394 | 0.397 | | 0.34 | 0.399 | 0.37 | 0.370 |
| **30.5** | 0.416 | 0.465 | 0.454 | 0.445 | | 0.433 | 0.383 | 0.399 | 0.405 | | 0.352 | 0.407 | 0.376 | 0.378 |
| **31** | 0.426 | 0.473 | 0.466 | 0.455 | | 0.441 | 0.389 | 0.406 | 0.412 | | 0.365 | 0.414 | 0.384 | 0.388 |
| **31.5** | 0.438 | 0.484 | 0.473 | 0.465 | | 0.45 | 0.397 | 0.412 | 0.420 | | 0.375 | 0.421 | 0.391 | 0.396 |
| **32** | 0.466 | 0.492 | 0.484 | 0.481 | | 0.46 | 0.407 | 0.418 | 0.428 | | 0.385 | 0.428 | 0.398 | 0.404 |
| **32.5** | 0.476 | 0.499 | 0.493 | 0.489 | | 0.467 | 0.413 | 0.426 | 0.435 | | 0.396 | 0.434 | 0.406 | 0.412 |
| **33** | 0.48 | 0.507 | 0.503 | 0.497 | | 0.473 | 0.419 | 0.432 | 0.441 | | 0.405 | 0.441 | 0.411 | 0.419 |
| **33.5** | 0.509 | 0.515 | 0.512 | 0.512 | | 0.481 | 0.427 | 0.44 | 0.449 | | 0.415 | 0.448 | 0.421 | 0.428 |
| **34** | 0.522 | 0.523 | 0.52 | 0.522 | | 0.49 | 0.438 | 0.449 | 0.459 | | 0.426 | 0.453 | 0.428 | 0.436 |
| **34.5** | 0.524 | 0.529 | 0.528 | 0.527 | | 0.493 | 0.443 | 0.456 | 0.464 | | 0.435 | 0.459 | 0.435 | 0.443 |
| **35** | 0.537 | 0.537 | 0.538 | 0.537 | | 0.5 | 0.454 | 0.465 | 0.473 | | 0.446 | 0.464 | 0.444 | 0.451 |
| **35.5** | 0.546 | 0.545 | 0.546 | 0.546 | | 0.505 | 0.464 | 0.473 | 0.481 | | 0.455 | 0.471 | 0.45 | 0.459 |
| **36** | 0.555 | 0.554 | 0.553 | 0.554 | | 0.513 | 0.473 | 0.481 | 0.489 | | 0.465 | 0.474 | 0.461 | 0.467 |
| **36.5** | 0.56 | 0.561 | 0.561 | 0.561 | | 0.516 | 0.481 | 0.488 | 0.495 | | 0.474 | 0.48 | 0.467 | 0.474 |
| **37** | 0.567 | 0.566 | 0.568 | 0.567 | | 0.521 | 0.488 | 0.497 | 0.502 | | 0.48 | 0.487 | 0.476 | 0.481 |
| **37.5** | 0.572 | 0.579 | 0.576 | 0.576 | | 0.527 | 0.496 | 0.506 | 0.510 | | 0.49 | 0.491 | 0.483 | 0.488 |
| **38** | 0.577 | 0.584 | 0.581 | 0.581 | | 0.532 | 0.505 | 0.513 | 0.517 | | 0.495 | 0.495 | 0.491 | 0.494 |
| **38.5** | 0.585 | 0.594 | 0.589 | 0.589 | | 0.54 | 0.511 | 0.519 | 0.523 | | 0.499 | 0.501 | 0.496 | 0.499 |
| **39** | 0.591 | 0.605 | 0.596 | 0.597 | | 0.542 | 0.519 | 0.527 | 0.529 | | 0.51 | 0.505 | 0.504 | 0.506 |
| **39.5** | 0.589 | 0.61 | 0.602 | 0.600 | | 0.549 | 0.526 | 0.533 | 0.536 | | 0.518 | 0.51 | 0.513 | 0.514 |
| **40** | 0.59 | 0.617 | 0.61 | 0.606 | | 0.551 | 0.531 | 0.538 | 0.540 | | 0.513 | 0.516 | 0.518 | 0.516 |
| **40.5** | 0.59 | 0.619 | 0.617 | 0.609 | | 0.555 | 0.538 | 0.544 | 0.546 | | 0.518 | 0.523 | 0.525 | 0.522 |
| **41** | 0.601 | 0.624 | 0.625 | 0.617 | | 0.555 | 0.543 | 0.55 | 0.549 | | 0.525 | 0.529 | 0.528 | 0.527 |
| **41.5** | 0.606 | 0.635 | 0.632 | 0.624 | | 0.561 | 0.547 | 0.552 | 0.553 | | 0.539 | 0.533 | 0.532 | 0.535 |
| **42** | 0.61 | 0.642 | 0.637 | 0.630 | | 0.564 | 0.549 | 0.554 | 0.556 | | 0.541 | 0.537 | 0.537 | 0.538 |
| **42.5** | 0.604 | 0.645 | 0.643 | 0.631 | | 0.567 | 0.554 | 0.559 | 0.560 | | 0.547 | 0.541 | 0.541 | 0.543 |
| **43** | 0.61 | 0.655 | 0.649 | 0.638 | | 0.573 | 0.559 | 0.564 | 0.565 | | 0.558 | 0.546 | 0.545 | 0.550 |
| **43.5** | 0.608 | 0.654 | 0.653 | 0.638 | | 0.577 | 0.562 | 0.567 | 0.569 | | 0.568 | 0.55 | 0.549 | 0.556 |
| **44** | 0.611 | 0.651 | 0.664 | 0.642 | | 0.584 | 0.567 | 0.57 | 0.574 | | 0.567 | 0.55 | 0.553 | 0.557 |
| **44.5** | 0.615 | 0.66 | 0.666 | 0.647 | | 0.586 | 0.571 | 0.574 | 0.577 | | 0.574 | 0.557 | 0.557 | 0.563 |
| **45** | 0.63 | 0.668 | 0.677 | 0.658 | | 0.59 | 0.573 | 0.577 | 0.580 | | 0.582 | 0.56 | 0.561 | 0.568 |
| **45.5** | 0.635 | 0.669 | 0.679 | 0.661 | | 0.595 | 0.579 | 0.581 | 0.585 | | 0.587 | 0.566 | 0.564 | 0.572 |

| | | | | | | | | | | | | | | |
|---|---|---|---|---|---|---|---|---|---|---|---|---|---|---|
| **46** | 0.64 | 0.673 | 0.685 | 0.666 | | 0.598 | 0.583 | 0.584 | 0.588 | | 0.591 | 0.571 | 0.568 | 0.577 |
| **46.5** | 0.652 | 0.677 | 0.69 | 0.673 | | 0.599 | 0.586 | 0.586 | 0.590 | | 0.597 | 0.576 | 0.573 | 0.582 |
| **47** | 0.65 | 0.676 | 0.693 | 0.673 | | 0.603 | 0.59 | 0.587 | 0.593 | | 0.6 | 0.579 | 0.574 | 0.584 |
| **47.5** | 0.666 | 0.681 | 0.696 | 0.681 | | 0.608 | 0.593 | 0.591 | 0.597 | | 0.606 | 0.583 | 0.579 | 0.589 |
| **48** | 0.67 | 0.685 | 0.698 | 0.684 | | 0.608 | 0.596 | 0.594 | 0.599 | | 0.609 | 0.588 | 0.582 | 0.593 |
| **48.5** | 0.674 | 0.689 | 0.702 | 0.688 | | 0.612 | 0.6 | 0.596 | 0.603 | | 0.619 | 0.59 | 0.585 | 0.598 |
| **49** | 0.674 | 0.694 | 0.703 | 0.690 | | 0.614 | 0.601 | 0.598 | 0.604 | | 0.617 | 0.594 | 0.588 | 0.600 |
| **49.5** | 0.683 | 0.694 | 0.703 | 0.693 | | 0.616 | 0.603 | 0.6 | 0.606 | | 0.621 | 0.598 | 0.59 | 0.603 |
| **50** | 0.675 | 0.692 | 0.705 | 0.691 | | 0.619 | 0.604 | 0.603 | 0.609 | | 0.623 | 0.6 | 0.593 | 0.605 |
| **50.5** | 0.68 | 0.693 | 0.707 | 0.693 | | 0.622 | 0.604 | 0.605 | 0.610 | | 0.627 | 0.603 | 0.596 | 0.609 |
| **51** | 0.681 | 0.7 | 0.708 | 0.696 | | 0.632 | 0.608 | 0.606 | 0.615 | | 0.633 | 0.604 | 0.599 | 0.612 |
| **51.5** | 0.688 | 0.701 | 0.712 | 0.700 | | 0.631 | 0.612 | 0.609 | 0.617 | | 0.637 | 0.606 | 0.602 | 0.615 |
| **52** | 0.687 | 0.7 | 0.713 | 0.700 | | 0.635 | 0.612 | 0.611 | 0.619 | | 0.638 | 0.61 | 0.604 | 0.617 |
| **52.5** | 0.691 | 0.702 | 0.717 | 0.703 | | 0.636 | 0.615 | 0.612 | 0.621 | | 0.644 | 0.612 | 0.606 | 0.621 |
| **53** | 0.689 | 0.703 | 0.716 | 0.703 | | 0.637 | 0.616 | 0.615 | 0.623 | | 0.644 | 0.614 | 0.609 | 0.622 |
| **53.5** | 0.691 | 0.704 | 0.716 | 0.704 | | 0.641 | 0.621 | 0.617 | 0.626 | | 0.648 | 0.615 | 0.612 | 0.625 |
| **54** | 0.693 | 0.706 | 0.721 | 0.707 | | 0.646 | 0.623 | 0.619 | 0.629 | | 0.648 | 0.621 | 0.614 | 0.628 |
| **54.5** | 0.696 | 0.706 | 0.721 | 0.708 | | 0.646 | 0.623 | 0.621 | 0.630 | | 0.655 | 0.619 | 0.616 | 0.630 |
| **55** | 0.692 | 0.707 | 0.722 | 0.707 | | 0.648 | 0.629 | 0.623 | 0.633 | | 0.658 | 0.624 | 0.618 | 0.633 |
| **55.5** | 0.701 | 0.714 | 0.725 | 0.713 | | 0.652 | 0.629 | 0.624 | 0.635 | | 0.666 | 0.624 | 0.621 | 0.637 |
| **56** | 0.706 | 0.714 | 0.728 | 0.716 | | 0.653 | 0.634 | 0.627 | 0.638 | | 0.668 | 0.628 | 0.624 | 0.640 |
| **56.5** | 0.705 | 0.718 | 0.73 | 0.718 | | 0.657 | 0.634 | 0.629 | 0.640 | | 0.67 | 0.628 | 0.626 | 0.641 |
| **57** | 0.709 | 0.722 | 0.73 | 0.720 | | 0.66 | 0.637 | 0.631 | 0.643 | | 0.678 | 0.63 | 0.629 | 0.646 |
| **57.5** | 0.704 | 0.721 | 0.732 | 0.719 | | 0.661 | 0.639 | 0.632 | 0.644 | | 0.674 | 0.635 | 0.63 | 0.646 |
| **58** | 0.708 | 0.723 | 0.733 | 0.721 | | 0.665 | 0.641 | 0.633 | 0.646 | | 0.673 | 0.637 | 0.633 | 0.648 |
| **58.5** | 0.707 | 0.725 | 0.733 | 0.722 | | 0.668 | 0.642 | 0.635 | 0.648 | | 0.676 | 0.637 | 0.635 | 0.649 |
| **59** | 0.709 | 0.725 | 0.732 | 0.722 | | 0.669 | 0.642 | 0.637 | 0.649 | | 0.677 | 0.642 | 0.638 | 0.652 |
| **59.5** | 0.711 | 0.726 | 0.734 | 0.724 | | 0.671 | 0.644 | 0.639 | 0.651 | | 0.677 | 0.644 | 0.64 | 0.654 |
| **60** | 0.718 | 0.733 | 0.74 | 0.730 | | 0.673 | 0.647 | 0.641 | 0.654 | | 0.686 | 0.645 | 0.641 | 0.657 |
| **60.5** | 0.716 | 0.732 | 0.736 | 0.728 | | 0.675 | 0.648 | 0.643 | 0.655 | | 0.684 | 0.647 | 0.644 | 0.658 |
| **61** | 0.715 | 0.733 | 0.74 | 0.729 | | 0.674 | 0.649 | 0.644 | 0.656 | | 0.684 | 0.649 | 0.645 | 0.659 |
| **61.5** | 0.712 | 0.732 | 0.739 | 0.728 | | 0.679 | 0.652 | 0.645 | 0.659 | | 0.683 | 0.651 | 0.647 | 0.660 |
| **62** | 0.722 | 0.736 | 0.742 | 0.733 | | 0.681 | 0.654 | 0.649 | 0.661 | | 0.689 | 0.652 | 0.649 | 0.663 |
| **62.5** | 0.716 | 0.736 | 0.739 | 0.730 | | 0.679 | 0.655 | 0.649 | 0.661 | | 0.7 | 0.655 | 0.651 | 0.669 |
| **63** | 0.719 | 0.741 | 0.745 | 0.735 | | 0.683 | 0.657 | 0.651 | 0.664 | | 0.702 | 0.655 | 0.652 | 0.670 |
| **63.5** | 0.722 | 0.734 | 0.744 | 0.733 | | 0.684 | 0.658 | 0.653 | 0.665 | | 0.698 | 0.659 | 0.654 | 0.670 |
| **64** | 0.726 | 0.737 | 0.744 | 0.736 | | 0.684 | 0.66 | 0.653 | 0.666 | | 0.692 | 0.659 | 0.657 | 0.669 |
| **64.5** | 0.723 | 0.744 | 0.746 | 0.738 | | 0.685 | 0.662 | 0.655 | 0.667 | | 0.698 | 0.662 | 0.658 | 0.673 |
| **65** | 0.725 | 0.743 | 0.746 | 0.738 | | 0.683 | 0.663 | 0.655 | 0.667 | | 0.7 | 0.663 | 0.66 | 0.674 |
| **65.5** | 0.73 | 0.74 | 0.743 | 0.738 | | 0.689 | 0.664 | 0.658 | 0.670 | | 0.705 | 0.666 | 0.661 | 0.677 |
| **66** | 0.729 | 0.744 | 0.748 | 0.740 | | 0.688 | 0.666 | 0.658 | 0.671 | | 0.704 | 0.665 | 0.663 | 0.677 |
| **66.5** | 0.732 | 0.746 | 0.746 | 0.741 | | 0.689 | 0.669 | 0.661 | 0.673 | | 0.705 | 0.668 | 0.665 | 0.679 |
| **67** | 0.734 | 0.748 | 0.752 | 0.745 | | 0.691 | 0.669 | 0.66 | 0.673 | | 0.713 | 0.67 | 0.666 | 0.683 |
| **67.5** | 0.733 | 0.749 | 0.75 | 0.744 | | 0.691 | 0.671 | 0.662 | 0.675 | | 0.717 | 0.671 | 0.668 | 0.685 |
| **68** | 0.741 | 0.747 | 0.75 | 0.746 | | 0.698 | 0.672 | 0.664 | 0.678 | | 0.708 | 0.673 | 0.67 | 0.684 |
| **68.5** | 0.741 | 0.748 | 0.747 | 0.745 | | 0.7 | 0.674 | 0.665 | 0.680 | | 0.715 | 0.675 | 0.671 | 0.687 |
| **69** | 0.739 | 0.753 | 0.754 | 0.749 | | 0.696 | 0.675 | 0.666 | 0.679 | | 0.723 | 0.675 | 0.672 | 0.690 |
| **69.5** | 0.737 | 0.751 | 0.752 | 0.747 | | 0.697 | 0.677 | 0.668 | 0.681 | | 0.726 | 0.677 | 0.674 | 0.692 |
| **70** | 0.74 | 0.752 | 0.754 | 0.749 | | 0.698 | 0.677 | 0.668 | 0.681 | | 0.72 | 0.68 | 0.676 | 0.692 |
| **70.5** | 0.745 | 0.749 | 0.756 | 0.750 | | 0.698 | 0.68 | 0.669 | 0.682 | | 0.724 | 0.681 | 0.678 | 0.694 |
| **71** | 0.747 | 0.752 | 0.76 | 0.753 | | 0.7 | 0.681 | 0.673 | 0.685 | | 0.729 | 0.683 | 0.68 | 0.697 |
| **71.5** | 0.748 | 0.751 | 0.76 | 0.753 | | 0.702 | 0.681 | 0.672 | 0.685 | | 0.731 | 0.683 | 0.681 | 0.698 |

| | | | | | | | | | | | | | | |
|---|---|---|---|---|---|---|---|---|---|---|---|---|---|---|
| **72** | 0.752 | 0.753 | 0.759 | 0.755 | | 0.703 | 0.683 | 0.674 | 0.687 | | 0.736 | 0.685 | 0.683 | 0.701 |
| **72.5** | 0.755 | 0.755 | 0.764 | 0.758 | | 0.704 | 0.684 | 0.675 | 0.688 | | 0.737 | 0.685 | 0.685 | 0.702 |
